# Long-term optical imaging of the spinal cord in awake, behaving animals

**DOI:** 10.1101/2023.05.22.541477

**Authors:** Biafra Ahanonu, Andrew Crowther, Artur Kania, Mariela Rosa Casillas, Allan Basbaum

## Abstract

Advances in optical imaging approaches and fluorescent biosensors have enabled an understanding of the spatiotemporal and long-term neural dynamics in the brain of awake animals. However, methodological difficulties and the persistence of post-laminectomy fibrosis have greatly limited similar advances in the spinal cord. To overcome these technical obstacles, we combined *in vivo* application of fluoropolymer membranes that inhibit fibrosis; a redesigned, cost-effective implantable spinal imaging chamber; and improved motion correction methods that together permit imaging of the spinal cord in awake, behaving mice, for months to over a year. We also demonstrate a robust ability to monitor axons, identify a spinal cord somatotopic map, conduct Ca^2+^ imaging of neural dynamics in behaving animals responding to pain-provoking stimuli, and observe persistent microglial changes after nerve injury. The ability to couple neural activity and behavior at the spinal cord level will drive insights not previously possible at a key location for somatosensory transmission to the brain.

## MAIN TEXT

Understanding neural computation mechanisms and their relation to perception and behavior necessitates recording neural activity in awake, behaving animals, ideally along all points of the neuroaxis. To facilitate this, optical imaging approaches have rapidly advanced, as has the development of a suite of biosensors that allow researchers to detect biologically relevant molecules (Kim and Schnitzer, 2022). Although almost all studies of “pain processing” at the level of the spinal cord have relied on recordings in anesthetized preparations, spinal cord neural activity recordings in the awake behaving animal have clear and significant advantages (Cheng et al., 2019; Nelson et al., 2019; Iseppon et al., 2022). Anesthetized preparations are selected mainly due to the numerous technical challenges of awake state spinal imaging, which requires invasive non-terminal microsurgical procedures. For example, spinal cord flexibility during natural locomotion, exacerbated by breathing, produces motion artifacts in imaging data and hinders quantitative analysis. Not surprisingly, few studies have recorded spinal cord neural activity, long-term in awake animals (Cheng et al., 2016; Sekiguchi et al., 2016; Ju et al., 2022; Shekhtmeyster et al., 2023).

Awake state recording performed repeatedly over months makes it possible to track neural circuit activity before, during, and after the development of neurological disorders, such as the transition from acute to chronic pain. Long-term optical access to the spinal cord also allows one to observe disease progression in non-neural cells (e.g. microglia). Moreover, stimulus-behavior relationships help identify psychophysical thresholds that can be compared to the activity of single neurons, which can only be measured in the awake state. For example, animals do not always respond to every noxious stimulus or elicit the same behavior to the same stimulus. The ability to couple behavior to neural activity at the spinal cord level will provide important insights as to where this variability in pain perception and behavior arises and also introduces an ability to study a variety of analgesic manipulations that target the spinal circuit level.

A major impediment to long-term optical recording is the inevitable post-laminectomy fibrosis, which occurs regardless of whether the animal is anesthetized or awake. More recent long-term imaging methods have implanted a small metal chamber overlying the spinal cord segment of interest (Farrar et al., 2012; Figley et al., 2013). Following laminectomy and adherence of a cover glass over the exposed spinal cord, a continually accessible spinal cord window provided the ability to follow injury-associated axonal regeneration over time (Kerschensteiner et al., 2005; Farrar et al., 2012; Fenrich et al., 2012). Despite these advances, post-laminectomy fibrosis can recur and frequently necessitates repreparation to uncover the spinal cord before imaging, with the associated risk of spinal cord damage and undesired physiological responses.

Here we address experimental and computational challenges to enable reliable, long-term spinal cord imaging (**Extended Data Fig. 1a**). We demonstrate that polytetrafluoroethylene (PTFE)-based inhibition of the post-laminectomy fibrosis preserves long-term spinal cord window clarity, making it possible to record, for months from defined populations of spinal cord neuronal and non-neuronal (microglial) cells, without repreparation. Implanted mice displayed normal locomotion and sensitivity, without evidence of spinal cord pathology. To handle the extensive spinal cord motion during imaging and improve neural activity analysis, we combined deep-learning feature identification (Mathis et al., 2018) and control point registration along with non-rigid diffeomorphic (Vercauteren et al., 2009) and rigid image registration algorithms (Thevenaz et al., 1998). We demonstrate repeated imaging (over months) of single-cell neural activity of large numbers of genetically-defined lamina I dorsal horn projection neurons, as well as high-quality imaging of axons (Thy1-GFP mouse lines and tdTomato expressed in sensory neurons), cell bodies (GFP expression pan-neuronally in the CNS), and microglial (using CX3CR1-EYFP mice). Furthermore, this new method achieves cellular imaging in behaving mice and is fully compatible with the time course of long-term preclinical studies, such as those investigating the transition from acute to chronic pain, pharmacological interventions, learning, nerve regeneration after spinal cord injury, and neurodegeneration.

**Fig. 1.**
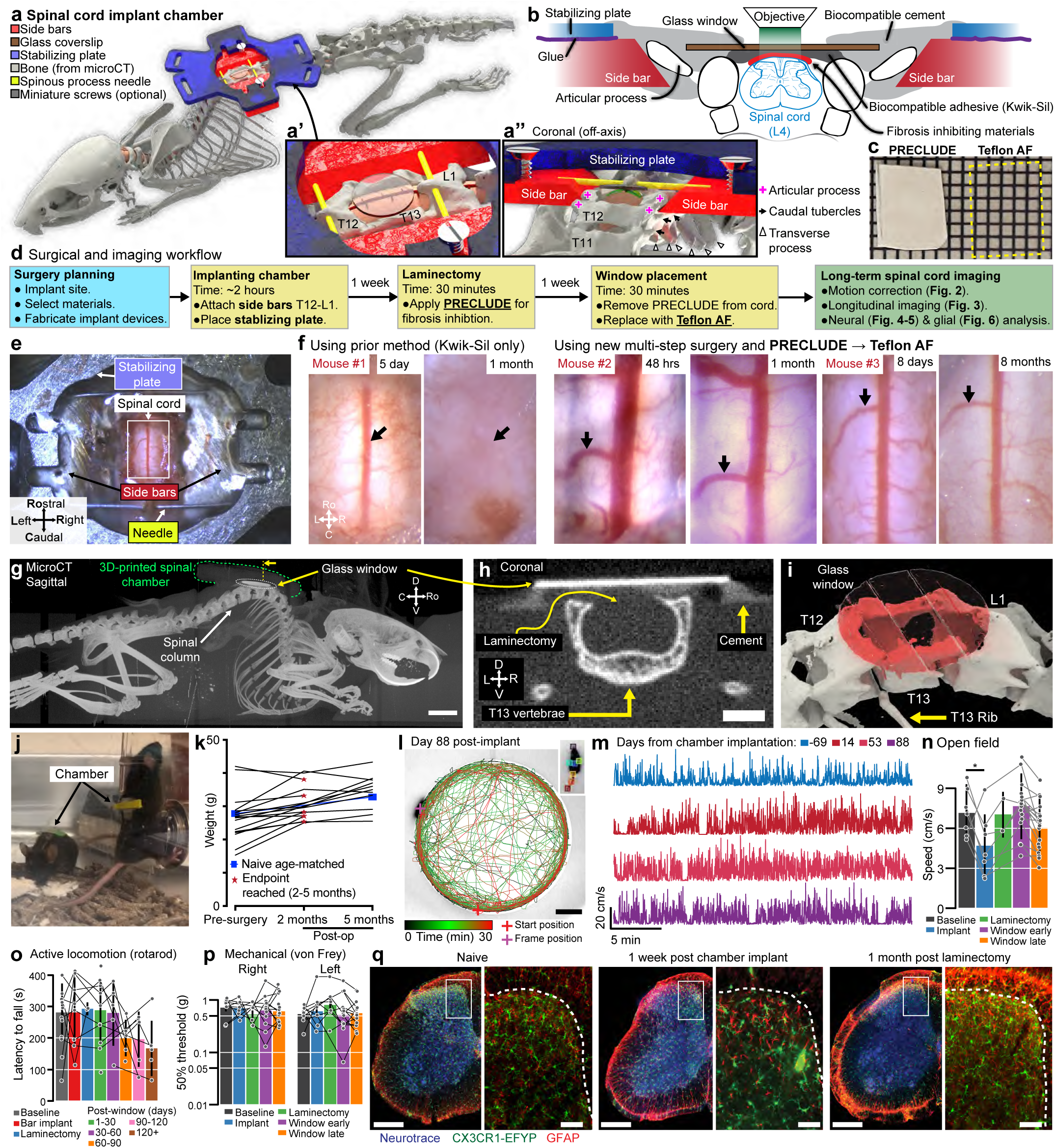
Surgical design and health validation for long-term spinal cord optical access in awake animals. **a.** 3D model of the recording chamber implanted at the T12-L1 vertebrae. Skeleton created using microCT data from a naive mouse (**Extended Data Fig. 4g**). Insets: **a’**, optical window (cover glass) provides access to the dorsal spinal cord post-T13 laminectomy and **a’’**, coronal view slightly off-axis shows a slice through side bars and highlights side bar placement adjacent to the articular processes and location of Teflon AF (green film). **b.** Diagram of chamber components that permit optical access to the lumbar (L4-L5) or lumbosacral (L5-S1) spinal cord. Super glue fixes the stabilizing plate to the two side bars. **c.** Two Teflon materials inhibit post-laminectomy fibrosis: opaque PRECLUDE is applied immediately after laminectomy (as in **d**), then removed and replaced subsequently by transparent Teflon AF fluoropolymer (as in **e**). **d.** Workflow for long-term spinal cord imaging includes three surgical steps. **e.** Overhead view of the spinal chamber one week after window placement. **f.** Using only silicone to cover the spinal cord leads to fibrosis and disappearance of the dorsal vein (black arrow) within a month (Mouse #1). Sequencing PRECLUDE and Teflon AF inhibits fibrosis, allowing visualization of the dorsal vein and ascending venules (black arrows) for months. Fluorescent imaging of mouse #2 and #3 in Fig. 3c and Fig. 5d, respectively. **g.** Whole body microCT sagittal max projection after chamber implantation. Chamber (green dashed line) fashioned using a 3D printed radiotransparent material (BioMed Clear). Scale bar, 5.0 mm. **h.** Coronal slice (yellow arrow and dashed line in **g**) from microCT, post laminectomy, shows intact surrounding bone in relation to the glass window. A surface layer of radiotransparent dental cement secures the glass window to the side bars. Scale bar, 1.0 mm. **i.** Multi-vertebral 3D reconstruction of microCT data in **g-h** confirms T13 window placement and bone integrity. **j.** Two mice exhibiting normal behaviors after chamber implant (see **Supplementary Video 1**). **k.** Body weight of chamber-implanted mice (n = 16) pre- and post-surgery compared to age-matched controls (n = 2). **l.** Tracking of open field locomotion (30 min) after chamber implant. Inset: DeepLabCut markers of individual body parts used for openfield tracking. Scale bar, 10 cm. **m.** Locomotor speed of the same mouse during 30-min sessions, pre- and post-surgery (Stage 1, side bar). **n.** Mean open field locomotor speed comparing naïve (n = 7) and post-surgery mice at different stages (n = 7, 2, 18, and 19, respectively) with “Window late” indicating beyond 30 days post window procedure. Most mice in **n-p** are used for imaging. Bar plot and error bars in **n**-**p** are mean ± SD, gray lines indicate animals measured across multiple stages. **o.** Mean latency to fall in final (3rd) trial on an accelerating rotarod comparing naïve (n = 14) and post-surgery mice at different stages (n = 12, 2, 10, 5, 5, 5, 5, respectively). **p.** Von Frey mechanical thresholds comparing naïve (n = 9) and post-surgery mice at different stages (n = 7, 6, 13, and 19, respectively) with “Window late” indicating beyond 30 days post window procedure. **q.** Microglia (CX3CR1-EYFP) and astrocyte (α-GFAP) immunofluorescence of 100-µm thick L4 sections before and after chamber implant. Scale bars, 300 µm and 50 µm (zoom).

## RESULTS

### Spinal chamber design and implant procedures

#### Design and fabrication of the implant components

To gain stable access to the lumbar spinal cord, we fabricated three low-cost implant components by laser cutting sheet metal (0.9-mm thickness, **Fig. 1a**) which together form a spinal chamber. These were either made of biocompatible metal (stainless steel, titanium, etc.) or 3D-printed biocompatible materials (FormLabs BioMed Clear or Surgical Guide) (**Extended Data Fig. 1b, d-h**). Specifically, we use two ∼9- × 10-mm metal pieces (side bars), one on each side of the vertebra and a third ∼20- × ∼24-mm piece (stabilizing plate) that connects the side bars over the top, forming an elevated platform across the spinal column (**Fig. 1b**). Additionally, we manually taper the side bars before surgery for paravertebral fitting (**Extended Data Fig. 2c**). To facilitate surgery on the lumbar spine, we remodeled a surgical table based on commercially available and 3D-printed components (Farrar et al., 2012) (**Extended Data Fig. 1c**). Adjustable posts on each side of the surgical table include small clamps that manipulate the implantable components. We provide detailed blueprints of the surgery set-up, side bars, and stabilizing plate (**Extended Data Fig. 1c-e**). The combined metal components weigh ∼1.3 g, comparable to material implants developed for head-mounted imaging (Ghosh et al., 2011), and are readily supported by the mouse (**Supplementary Video 1**).

**Fig. 2.**
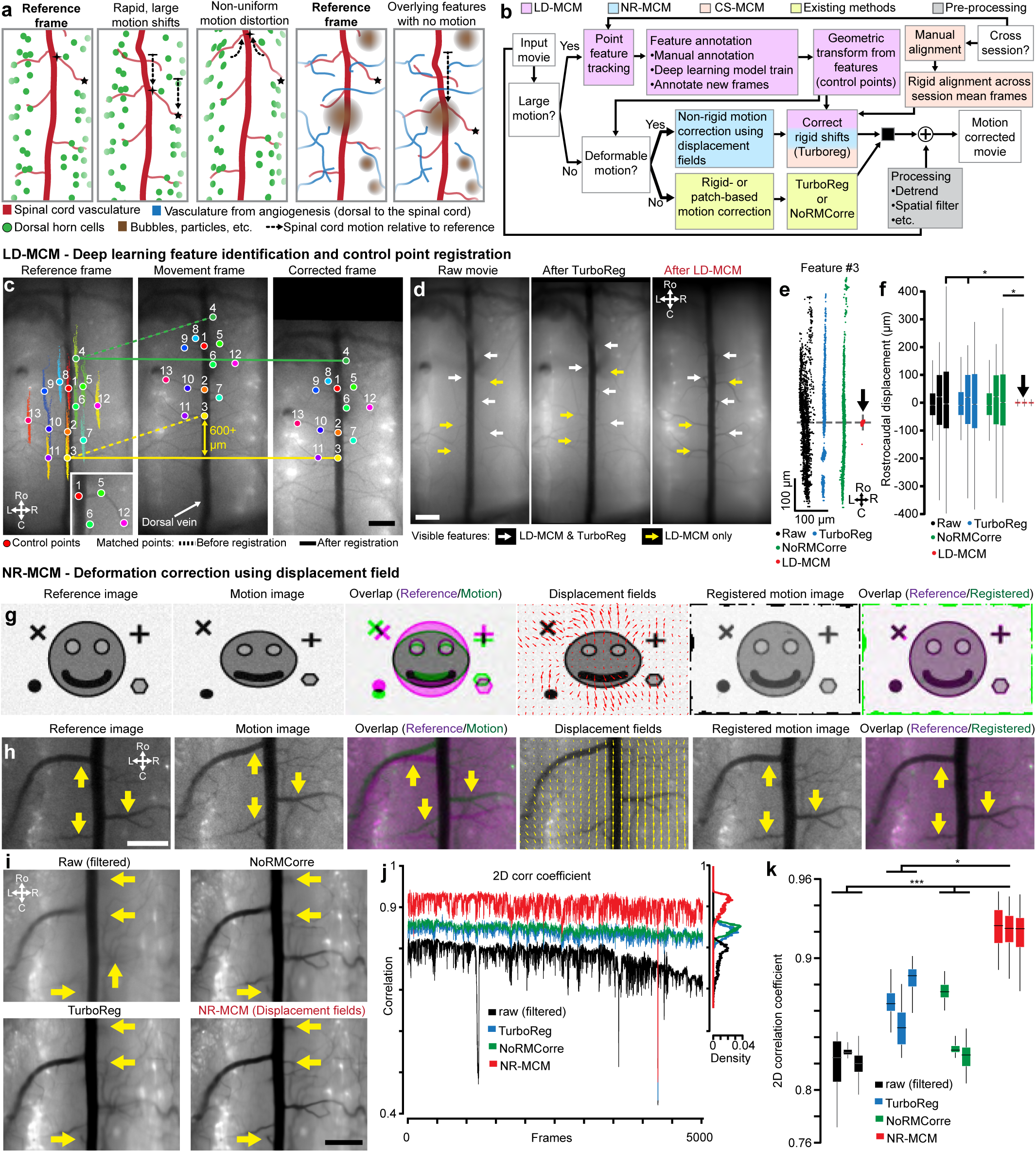
Computational correction of spinal cord motion in awake animals. **a.** Imaging a large spinal cord field is subject to several types of motion artifacts. **b.** Modular motion correction pipeline that addresses each of the issues outlined in **a**: features identified using deep learning followed by control point and rigid registration (large displacement motion correction method, **LD-MCM**), deformation correction using displacement fields followed by rigid registration (non-rigid motion correction method, **NR-MCM**), and manual or automated cross-session motion correction (**CS-MCM**). **c.** LD-MCM utilizes deep learning, here DeepLabCut (DLC), to identify vascular features after manual annotation and training. LD-MCM uses these features to transform and register frames to a reference frame’s features. Point clouds overlapping each feature in the reference frame show rostrocaudal and mediolateral extent of motion during each frame of an entire imaging session (2.31 mins, 20 Hz). Inset, zoomed in field of view shows overlap of markers with distinct vasculature features. Scale bar, 300 µm. **d.** Mean projection image across all movie frames for the raw movie and after TurboReg or LD-MCM demonstrates reduced motion with LD-MCM. Arrows indicate features seen only in LD-MCM (yellow) and others that are barely visible after TurboReg (white). Scale bar, 300 µm. **e.** Point clouds for rostrocaudal and mediolateral movement of feature #3 (as in **c**) after motion correction with TurboReg, NoRMCorre, and LD-MCM. Each dot represents the location of that feature on an individual frame during an imaging session (2.31 min, 20 Hz). Arrow indicates location of LD-MCM points, showing negligible post-procedure motion. **f.** Boxplots show rostrocaudal displacement of the spinal cord relative to the mean location in the raw movie and after motion correction with TurboReg, NoRMCorre, and LD-MCM over all features, for each of 3 movies (n = 2 mice). Features for all methods identified with DLC. Arrow highlights negligible post-LD-MCM motion (as in **e**). **g.** Synthetic image (116 × 77 px) before and after image alignment with NR-MCM. Vectors (red) indicate the displacement field orientation and magnitude at a given pixel location; the vector field is sub-sampled (5x) and magnitude scaled for display purposes. **h.** NR-MCM (as in **g**) on an example frame from one-photon fluorescence imaging of spinal cord GCaMP-expressing neurons (same mouse as Fig. 5d). Yellow arrows highlight features that are aligned after registration. Scale bar, 300 µm. **i.** Mean projection images of the first 5,000 frames in a movie (∼12.5 min, 20 Hz) show improved motion correction with NR-MCM compared to TurboReg and NoRMCorre. Yellow arrows highlight stable features in NR-MCM movies. Scale bar, 300 µm. **j.** 2D correlation coefficient of all frames to the mean frame of the movie (as in **i**) for NR-MCM compared to raw, TurboReg, and NoRMCorre. All movies were spatially filtered to remove large magnitude, low-frequency changes in fluorescence, which artificially enhances correlations. Right inset: histogram of correlation coefficients across all frames; vertical axis is aligned to that in the main graph. **k.** Boxplots summarize results, as in j, over 3 movies (n = 2 mice). Boxplots in all figures display the 1st, 2nd, and 3rd quartiles with whiskers indicating 1.5*IQR; outliers are omitted.

#### Paraspinal implant procedure

To achieve a continually accessible window to the spinal cord lumbar enlargement, we perform three separate surgical procedures (**Fig. 1d-e**). These surgeries occur over three weeks, totaling ∼5 hours of general anesthesia per animal. In the first surgery (2-3 hours) (**Supplementary Fig. 1**, **Supplementary Video 2**), we expose and fuse three successive vertebrae (T12-T13-L1) using the side bars and the stabilizing plate as structural supports. To access vertebral bone, we sever paraspinal musculature at the midline and retract laterally. To facilitate an intraoperative assembly of the metal components and improve the stability of the spinal chamber attachment, we bore thin needles (33G, 0.2-mm OD) perpendicularly through the dorsal spinous process (DSP) of the most rostral (T12) and caudal vertebra (L1) (see anatomy in **Extended Data Fig. 2a-i**). We then clamp and advance the side bars under the needles to rest against the vertebral wall above the intervertebral foramina (**Extended Data Fig. 1k**). Before freeing this structure from the clamps, we superglue the 33G needles to the side bars and lightly seal the edges of the incised tissue with VetBond. We then superglue the stabilizing plate over both side bars, which completes the metal assembly. Subsequent steps fix the bone and implant using resin polymethyl methacrylate (PMMA, C&B Metabond kit). Importantly, we uncover the laminar bone of the middle vertebra (T13) before the PMMA cement sets, allowing for the subsequent laminectomy, which is generally performed 1 week after the chamber implantation; see **Methods** and **Supplemental Fig. 1** for details.

#### Inhibiting post-laminectomy fibrosis with synthetic fluoropolymer materials

As emphasized above, a major impediment to chronic spinal cord recordings is the post-laminectomy fibrosis that rapidly obscures the spinal cord’s dorsal surface, drastically limiting optical access to the underlying tissue (**Fig. 1f**). Although filling the space directly above the spinal cord with transparent silicone-based plastic can abate fibrosis, the success rate is highly variable (Farrar et al., 2012; Figley et al., 2013). Despite using a variety of silicone adhesives (Kwik-Sil, PDMS, Q3-6575, Dowsil, and others), as previously reported, we found that regrowth directly over the spinal cord is inevitable. Interestingly, because synthetic dural substitutes are used effectively over the long term in human neurosurgical procedures, here in mice, after laminectomy, we tested a cohort of artificial dura plastic materials for their efficacy and permanence as a dural substitute (Duraseal, Gelfoam, ePTFE). What stood out from all these products was ePTFE, i.e. Teflon (GORE® PRECLUDE® Membrane, Gore medical), which completely inhibited fibrotic and dural regrowth for months.

A significant limitation, however, is that the PRECLUDE® membrane is opaque **(Fig. 1c)**, which is incompatible with optical imaging (**Extended Data Fig. 2j**). Thus, we identified a crystal-clear amorphous fluoropolymer, Teflon AF 2400 **(Fig. 1c)**, which also displayed long-term regrowth inhibitory properties (**Fig. 1f**). The extremely transparent properties of Teflon AF (>95% transmittance from λ=300–2000 nm) were critical to our studies, and other applications in preclinical research (Resnick and Buck, 2002; Anamelechi et al., 2005; Yang et al., 2008; Czolkos et al., 2012). Further, as Teflon AF has a visible and near-IR refractive index of *n*_TAF_ ≈ 1.28–1.29 compared to nervous system tissue of *n*_brain_ ≈ 1.36–1.38 (Binding et al., 2011), we predicted that Teflon AF would minimally affect optical access. We confirmed this by conducting one- and two-photon imaging of 1-µm fluorospheres on glass slides and found that the presence of Teflon AF did not affect optical imaging quality (**Extended Data Fig. 2k-m**).

#### Laminectomy, regrowth inhibition, and window placement

In the second surgical procedure (0.5-1 hr) (**Supplementary Fig. 2**, **Supplementary Video 3**), we expose the spinal cord (L4/5) by laminectomy of the middle vertebra (T13). Using fine forceps, we delicately separate dura from the center of the exposure (unless it is removed together with bone during the laminectomy), taking care to preserve the underlying leptomeninges (arachnoid and pia mater). We then place PRECLUDE® over the leptomeninges, surround it with a gelatin sponge, and set it in place with Kwik-sil. Although we typically proceed to the third step within a week, this procedure can be performed up to a month after placing the PRECLUDE® membrane.

In the third and final surgical procedure (0.5-1 hr) (**Supplementary Fig. 3**, **Supplementary Video 4**), we swap out the opaque PRECLUDE® Teflon for the transparent form, Teflon AF 2400. We then set a #0 glass coverslip over the Teflon AF using Kwik-Sil as an adhesive between these layers. Finally, we stabilize this configuration with bone cement, making a base on the metal assembly up to the rim of the coverslip. With the Teflon AF material set and the surrounding soft tissue sealed from the environment, the underneath leptomeningeal layer has a stable, clear quality for long-term optical access (**Fig. 1f**). We were able to retain optical access for months to over a year, which allowed visualization of the meninges as well as the dorsal spinal cord, including the midline dorsal vein and dorsal ascending venules (dAVs) **(Fig. 1f**, black arrows).

**Fig. 3.**
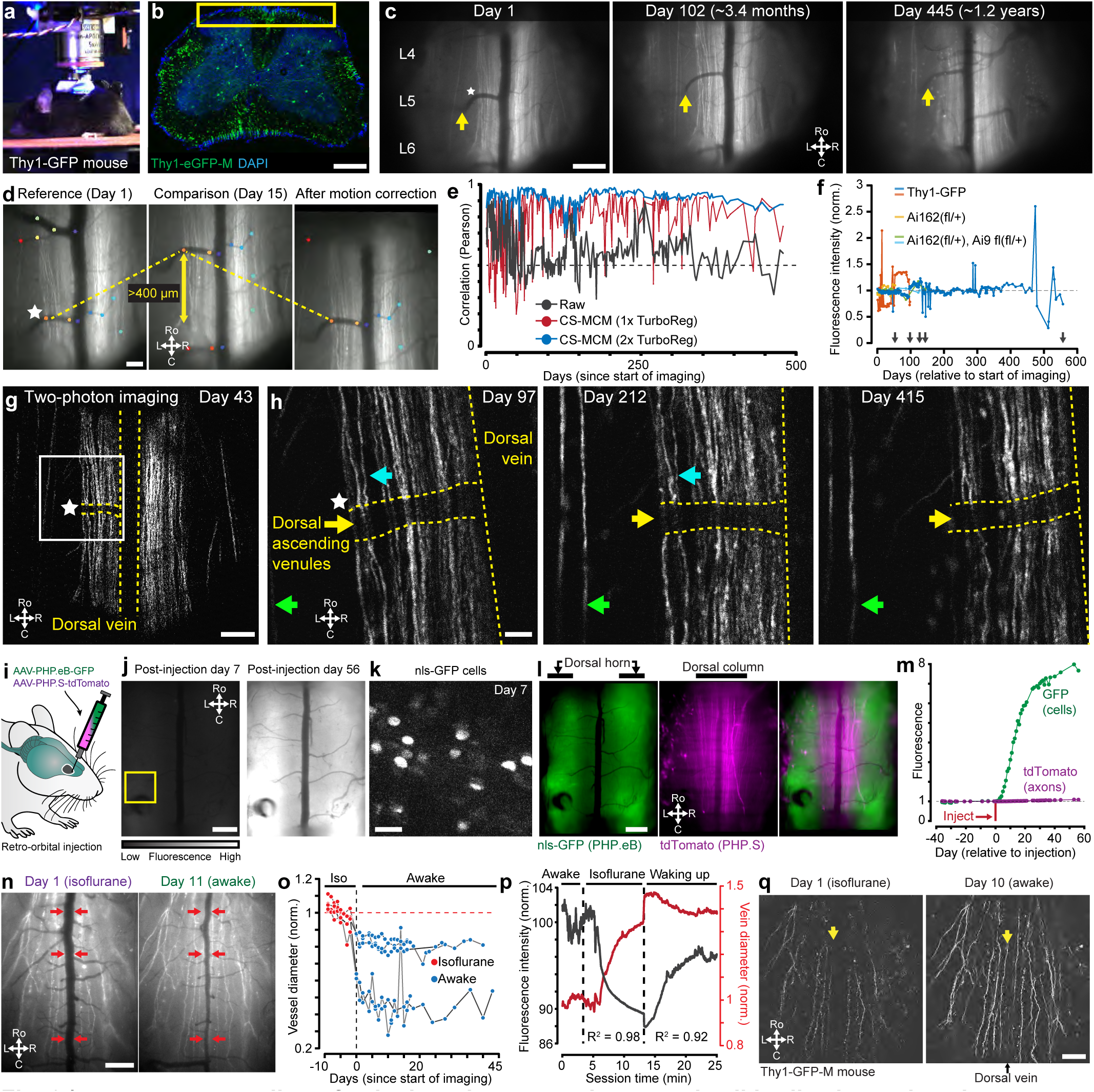
Long term recording of spinal cord axons and neuronal cell bodies in awake mice. **a.** Chamber implanted mouse in one- and two-photon imaging set up. **b.** Confocal fluorescence image of 100-µm-thick coronal tissue section from a Thy1-GFP mouse shows GFP+ dorsal column axons. Yellow box indicates the approximate region imaged in **c**, **d**, and **g**. Scale bar, 300 µm. **c.** One-photon imaging over 1.2 years in an awake, behaving Thy1-GFP mouse. Yellow arrows indicate the same dorsal ascending venule (dAV) observed across all imaging sessions. Scale bar, 300 µm. **d.** Frames from two different imaging sessions in a Thy1-GFP mouse (as in **c**) shows the large rostrocaudal shifts that can occur across sessions during long-term imaging. Control points (colored dots) are DLC-tracked vascular features used to correct for the large displacement (see Fig. 2c). Scale bar: 100 µm. **e.** Pearson correlation to the mean frame of raw movie shows improved cross-session alignment after CS-MCM, including manual alignment, plus one or two TurboReg iterations on different aspects of the cross- session movies. **f.** Fluorescence intensity from one-photon imaging sessions of Thy1-GFP and background (Ai162(fl/+); Ai9 fl(fl/+) with no signal) mice (n = 5) normalized by mean intensity across equivalent acquisition cameras, which allows comparison across CCD and sCMOS cameras used during long-term imaging. Gray lines indicate the last imaging session day for a given animal. **g.** Two-photon fluorescence imaging of GFP+ axons 43 days after start of imaging. Star in **g**-**h** indicates the same dAV as seen in one-photon data in **c**-**d**. Scale bar, 200 µm. **h.** Zoomed-in view of GFP+ axons near a dAV (white box in **g**) over time. Similar color arrows indicate individual axons common across imaging sessions. Scale bar, 50 µm. **i.** Retro-orbital injection of AAV-PHP.eB-CAG-nls-GFP and AAV-PHP.S-CAG-tdTomato enables concurrent imaging of spinal cord dorsal horn neurons (GFP) and ascending axons of primary sensory neurons (tdTomato). **j.** One-photon bilateral spinal cord imaging shows increasing GFP expression from 7 to 56 days after retro-orbital injection. Constant contrast and brightness between days. Scale bar, 300 µm. **k.** Two-photon imaging in the dorsal horn (yellow box in **j**) shows expected nuclear localization of the nls-GFP signal. Scale bar, 100 µm. **l.** One-photon imaging of nls-GFP in dorsal horn neurons and tdTomato in dorsal column axons in the same mouse. Scale bar, 300 µm. **m.** Near daily imaging of GFP and tdTomato fluorescence normalized to baseline (pre retro-orbital injection) from mouse in **j**-**k**. TdTomato fluorescence in axons is dimmer; see **Extended Data Fig. 7e** for expanded view. **n.** Imaging Thy1-GFP under 2% isoflurane anesthesia or while awake. Red arrows denote reduced dorsal vein diameter in the awake mouse. Scale bar, 300 µm. **o.** Vessel diameter (normalized per animal to isoflurane baseline sessions) in anesthetized vs awake mice (n = 5, gray lines) recorded over several weeks. **p.** Temporal change of vessel diameter and whole-frame fluorescence (normalized to 4-min awake baseline) within a single imaging session in a Thy1-GFP mouse (as in **n**) before and after anesthesia (2% isoflurane). R^2^ between fluorescence and dorsal vein diameter given for relevant session epochs. **q.** Two-photon imaging of GFP+ axons (Thy1-GFP mouse as in **n**) also shows increased signal of axons near the midline during awake imaging; these axons are obscured under anesthesia. Scale bars, 300 µm.

### Normal animal behavior and preserved spinal cord integrity after implant procedures

#### 3D printed spinal chamber for microCT imaging and surgery validation

There are certain studies in which non-invasive imaging modalities can be combined with fluorescence imaging. These include multi-scale neural dynamics with Ca^2+^ imaging and functional magnetic resonance imaging (fMRI) (Lake et al., 2020). However, existing spinal cord implant devices use metals or similar materials that are incompatible with or cause artifacts in MRI or micro X-ray computed tomography (microCT)—as can be seen in validation studies (**Extended Data Fig. 4a-d**) and prior papers by the lack of bone and soft tissue around the implant in the microCT reconstructions. Using advances in 3D printing of biocompatible materials, we found that FormLab’s Surgical Guide and BioMED Clear resins are both bio- and microCT-compatible (**Extended Data Fig. 4e-f**). Attaching stabilizing plates to the side bars with adhesives, instead of metallic miniature screws, as is used in other spinal cord preparations, also improves compatibility with microCT (**Extended Data Fig. 4h-i, Supplementary Video 4**). To verify the laminectomy and spine structural integrity, we performed the implant surgery and laminectomy, then conducted microCT imaging (**Fig. 1g-i, Supplementary Videos 5-6**). Importantly, the 3D-printed side bars and stabilizing plate did not occlude the laminectomy location nor other parts of the body. Future studies can use contrast agents or MRI to visualize the spinal cord and dorsal root health and anatomy over time.

**Fig. 4.**
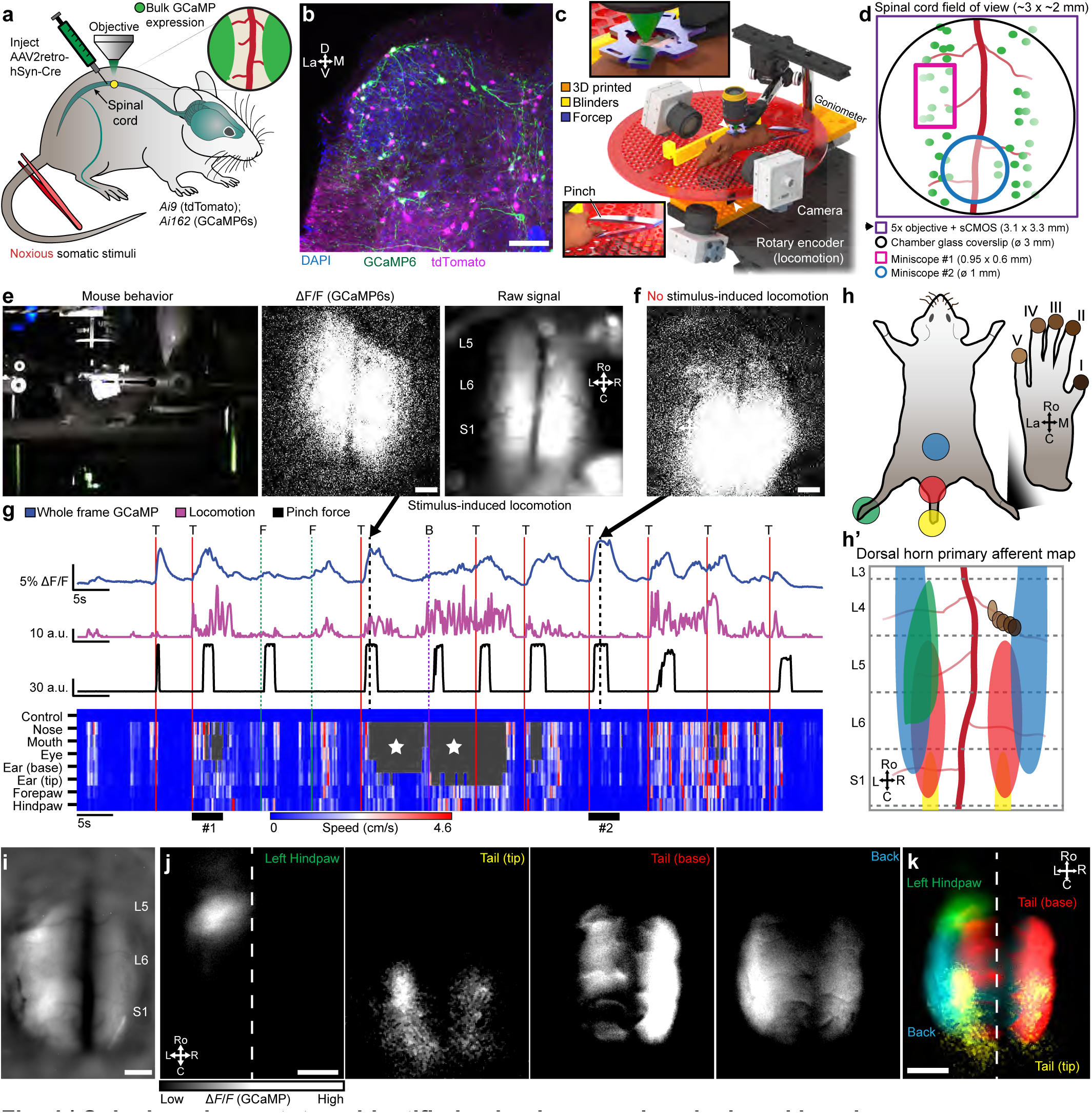
Spinal cord somatotopy identified using large-scale spinal cord imaging. **a.** Bulk expression of GCaMP6s throughout the spinal cord dorsal horn is achieved after intraspinal injection of AAV2retro-hSyn-Cre virus into Ai9 and Ai162 (Cre-dependent GCaMP6s) mice. **b.** GCaMP6s and tdTomato expression in neurons throughout the superficial and deep laminae of the sacral spinal cord after AAV intraspinal injection (as in **a**). Scale bar, 100 µm. **c.** 3D model of vertebral-fixed setup for imaging and stimulus delivery. Behavior measurement uses a rotary encoder and high-speed cameras. Similar setup is used for awake imaging (Fig. 3**-6**). **d.** Bilateral spinal cord imaging using widefield microscopy with low-magnification objectives and large-sensor sCMOS cameras. Miniscope #1 = Inscopix nVista, Miniscope #2 = Miniscope v4.4. **e.** Tail pinch (red bar) induces large increase in dorsal horn bulk GCaMP6s fluorescence, coincident with increased locomotion. Arrow points to the session time point in **g**. Scale bar, 300 µm. **f.** Widespread increase in stimulus-evoked dorsal horn GCaMP6s fluorescence can occur despite minimal locomotion. Arrow as in **e**. Scale bar, 300 µm. **g.** Neural activity (whole frame GCaMP6s fluorescence) aligned to locomotion, body part movement, and force of pinch applied to the tail (T) or back (B) or by innocuous tactile stimulation to the forepaw (F). Bottom heatmap is DLC-based detection of movement of individual body parts. Numbers below heat map highlight when: #1, neural activity correlated with increased behavior; #2, stimulus did not induce locomotion, but there was head movement. Gray portions of the heat map with stars are those where DLC had low confidence in predicting body part location, likely due to occlusion by the mouse moving out of the camera field of view. Pinch stimulus magnitude measured using force-sensitive resistor (values given in arbitrary units). **h.** Schematic illustrates location of body parts stimulated (recordings in **i**-**k**). **h’**, illustrates dorsal horn primary afferent terminal map based on anatomical tracing studies in mice and rats (Takahashi et al., 2002, 2003; Odagaki et al., 2019). **i.** One-photon fluorescence imaging field of view from GCaMP6s mouse (as in **a**) recorded in **j**-**k**. Mean projection image after bandpass filtering highlights vasculature. Scale bar, 200 µm. **j.** Neural activity (mean projection image of Δ*F*/*F*) in response to pinch of different parts of the body. Scale bar, 300 µm. **k.** Neural activity maps from **j** superimposed to show dorsal horn somatotopy. Scale bar, 300 µm.

#### Preserved sensitivity and locomotion

We regularly monitored mice with implants (**Fig. 1j**) and used weight as a quantifiable indicator of health. The postoperative weight did not decline at 2 or 5 months (n = 22 animals) and mirrored the weight gain of naive mice (**Fig. 1k** and **Extended Data Fig. 4k**). We also verified animal health post-surgery using physiological measures along with sensory and locomotor behavioral assays (**Fig. 1l-p**). Spinal-implanted mice move throughout the open field as well as naive animals (**Supplementary Video 7**). Quantitative data confirm that implanted animals after the final surgery achieve similar open field speeds as their pre-surgery baseline (**Fig. 1l-n**). Implanted animals can repeatedly perform the rotarod task, with a trend toward decreasing performance after 2 months (**Fig. 1o**, **Extended Data Fig. 4l**). Importantly, implanted animals displayed normal mechanical sensitivity for months after surgery (**Fig. 1p**).

#### Post-mortem tissue analysis

Post-mortem examination of the lumbar enlargement did not reveal significant histopathology (**Fig. 1q, Extended Data 3a-b**) unless signs of fibrosis were present. Between 2-5 months after implantation, we observed fibrosis development in 5 of 20 preparations. These animals with fibrosis showed significant spinal cord pathology (**Extended Data 3c**) accompanied by various behavioral signs (paralysis, altered gait, skin lesions). We only conducted longitudinal imaging on animals without signs of significant fibrosis.

### Motion correction of the spinal cord in awake animals

#### A pipeline for spinal cord motion correction

As noted above, awake state spinal cord recordings have significant challenges compared to recordings in the brain: locomotion-driven spinal cord movement leading to large rostro-caudal shifts (>650 microns in certain animals within and across sessions), rapid non-rigid deformations of the field of view, and presence of obstacles—such as neovascularizations (**Extended Data Fig. 8a**), bubbles, etc.—that move differently from the primary tissue of interest (**Fig. 2a**). Here, we developed a multi-step, hierarchical workflow that allows users to determine which algorithm is best suited to handle the imaging data (**Fig. 2b**). The workflow consists of a large-displacement motion correction method (**LD-MCM**), a non-rigid motion correction method (**NR-MCM**), and a semi- or fully-automated cross-session motion correction method (**CS-MCM**). We briefly discuss each technique below; see **Methods** for details.

#### Deep learning feature identification and control point registration to correct large spinal cord motion

To automate the correction of large rostrocaudal shifts, LD-MCM uses deep learning (e.g. DeepLabCut [DLC](Mathis et al., 2020)) which tracks features followed by control point and rigid registration (**Fig. 2c**). Although image alignment can be done using feature detectors—such as MSER, FAST, SIFT, and others (Işık, 2015; Liu et al., 2021)—spinal cord imaging movies contain a variety of features across multiple layers that cause these methods to identify non-relevant features. Here, we tracked consistent vasculature, as a proxy for spinal cord motion, which improved registration (**Extended Data Fig. 5a**). To achieve this, we used deep learning methods in which features are manually annotated on a subset of frames and a model is trained to identify them in novel video frames (**Extended Data Fig. 5b-f)**. We found that LD-MCM improved motion correction in spinal cord imaging movies compared to widely used TurboReg or NoRMCorre (Thevenaz et al., 1998; Pnevmatikakis and Giovannucci, 2017) methods (**Fig. 2d-f, Supplementary Video 8**). With minimal training (20 frames from one session), we consistently identified vasculature features across months of imaging, from which reliable motion correction is possible with minimal user input **(Extended Data Fig. 5g)**.

**Fig. 5.**
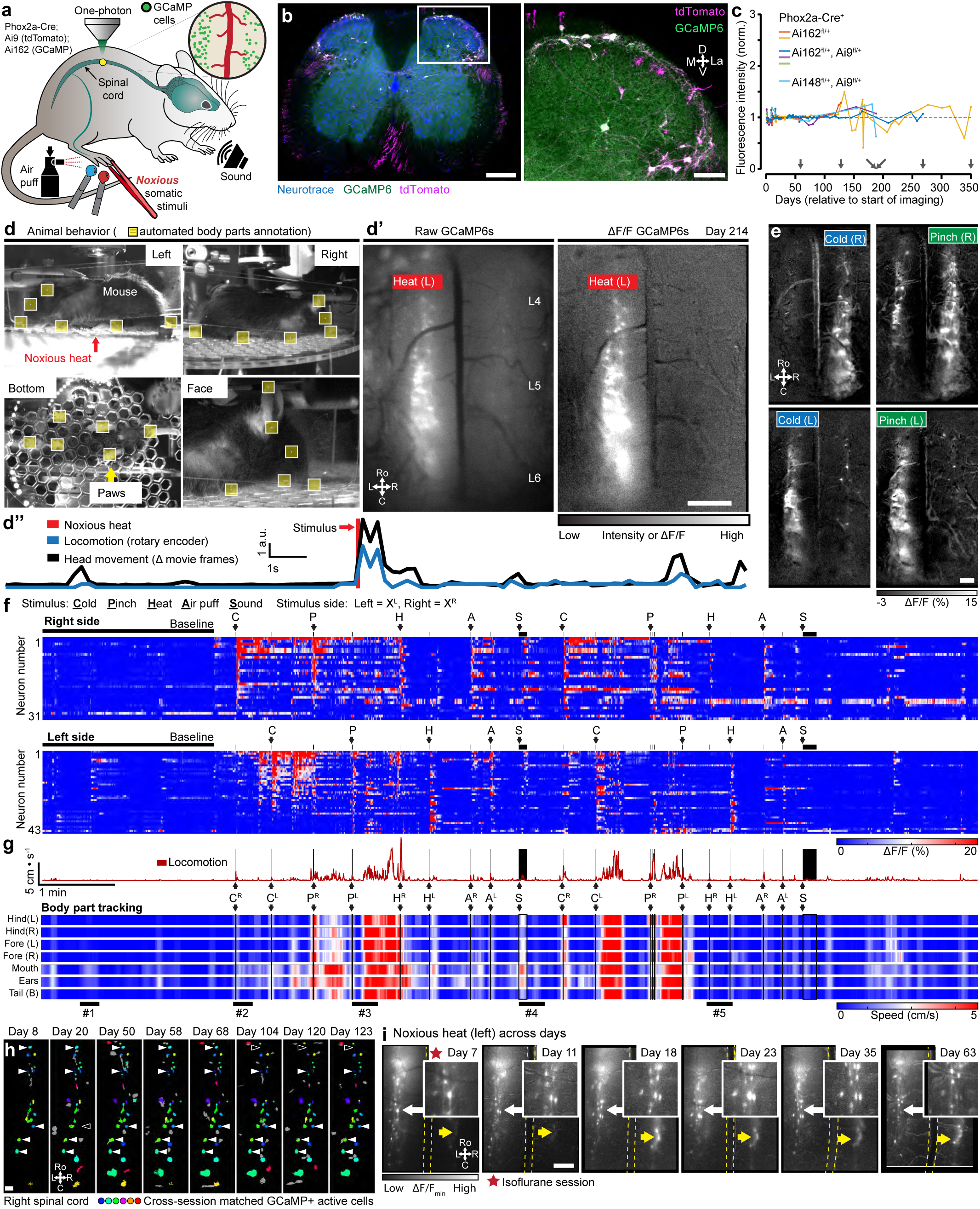
Long-term bilateral imaging of spinal cord dorsal horn neural activity in awake mice. **a.** Genetic approach to image activity of individual spinal cord projection neurons (SCPNs) that express GCaMP6s and/or tdTomato in Phox2a-Cre; Ai162/Ai148; Ai9 mice. Hindpaws received noxious thermal (hot and cold water) or mechanical (forceps pinch or air puff) stimuli; a startle-inducing stimulus (sound) is presented near the head. **b.** Confocal fluorescence image of 50-µm-thick coronal tissue sections showing location of dorsal horn SCPNs. Inset (white box) magnified to the right. Scale bars, 300 and 100 (inset) µm. **c.** Fluorescence intensity from individual one-photon imaging sessions spanning months from SCPN-GCaMP6s mice (n = 6), normalized as in Fig. 3f. **d.** Simultaneous monitoring of L4-L6 spinal cord neuronal activity and behavior on 214 days after the first imaging session. Simultaneous acquisition of multi-angle behavioral data from multiple cameras processed using DLC to track movement of different body parts (yellow boxes). **d’**, frames from a single time point showing neural activity (GCaMP6s) in individual SCPNs in response to noxious heat from the raw and Δ*F/F* movies. **d”**, trace of locomotion (measured from rotary encoder) and head movement (measured as a delta of frames from head-focused camera) before and after application of noxious stimuli (red vertical line). Scale bar in **d’**, 500 µm. **e.** Recording as in **d** shows ipsilateral, and weaker contralateral, responses of SCPNs to noxious cold and mechanical stimuli. Scale bar, 300 µm. **f.** Activity of individual SCPNs as in **d**, on the left or right spinal cord. Stimuli preceded by ∼2-min baseline recording. Letters above each black arrow indicate the stimulus presented (C: cold; P: pinch; H; heat; A: air puff; S: sound); black bar denotes duration of the sound stimuli. **g.** Locomotion aligned to neural activity and stimuli as in **f**. Lower heat map displays movement (i.e. speed) of specific body parts, as in **d**. Numbers below heat map highlight when: #**1**, spontaneous head movement is correlated with SCPN activity; #**2**, stimulus-induced locomotion and ipsilateral-only SCPN activity; **#3**, post-stimulus pause in behavior activity after ipsilateral SCPN activity followed by robust movement that correlates with onset of secondary burst of activity; #**4**, sound stimulus induces locomotor behavior with minimal SCPN activation; and #**5**, ipsilateral only SCPN activity in response to a noxious heat stimulus applied to each hindpaw. L and R superscripts for each stimulus denote left or right hindpaw application. **h.** SCPNs extracted from individual awake animal imaging sessions (except day 8) and aligned across days. Color indicates the same cell aligned across days; filled and open arrows indicate when that particular cell is or is not identified after cell extraction across imaging sessions, respectively. Scale bar, 100 µm. **i.** SCPN activity (mean projection of Δ*F*/*F*_min_ post-stimulus) after noxious heat applied to the left hindpaw across multiple imaging sessions. Yellow dotted lines indicate the dorsal vein. Yellow arrows show consistent activity in the dorsal horn contralateral to the stimulated hindpaw. Insets, white arrows indicate enlarged areas showing consistent response of the same neurons across multiple imaging sessions. Scale bar, 300 µm.

#### Diffeomorphic transformations to handle non-rigid spinal cord motion

To handle non-rigid motion, we adapted displacement field-based registration based on Maxwell’s Demons (Thirion, 1998; Vercauteren et al., 2009). These techniques can lead to registration of images containing a mix of deformations and spatial shifts of features in the imaging field of view (**Fig. 2g-h**) and have been used previously for cross-session and other motion correction in brain Ca^2+^ imaging (Reggiani et al., 2023). NR-MCM implemented on GCaMP spinal cord imaging movies in awake animals improved registration compared to TurboReg and NoRMCorre (**Fig. 2i-k, Extended Data Fig. 6, Supplementary Video 9**).

**Fig. 6.**
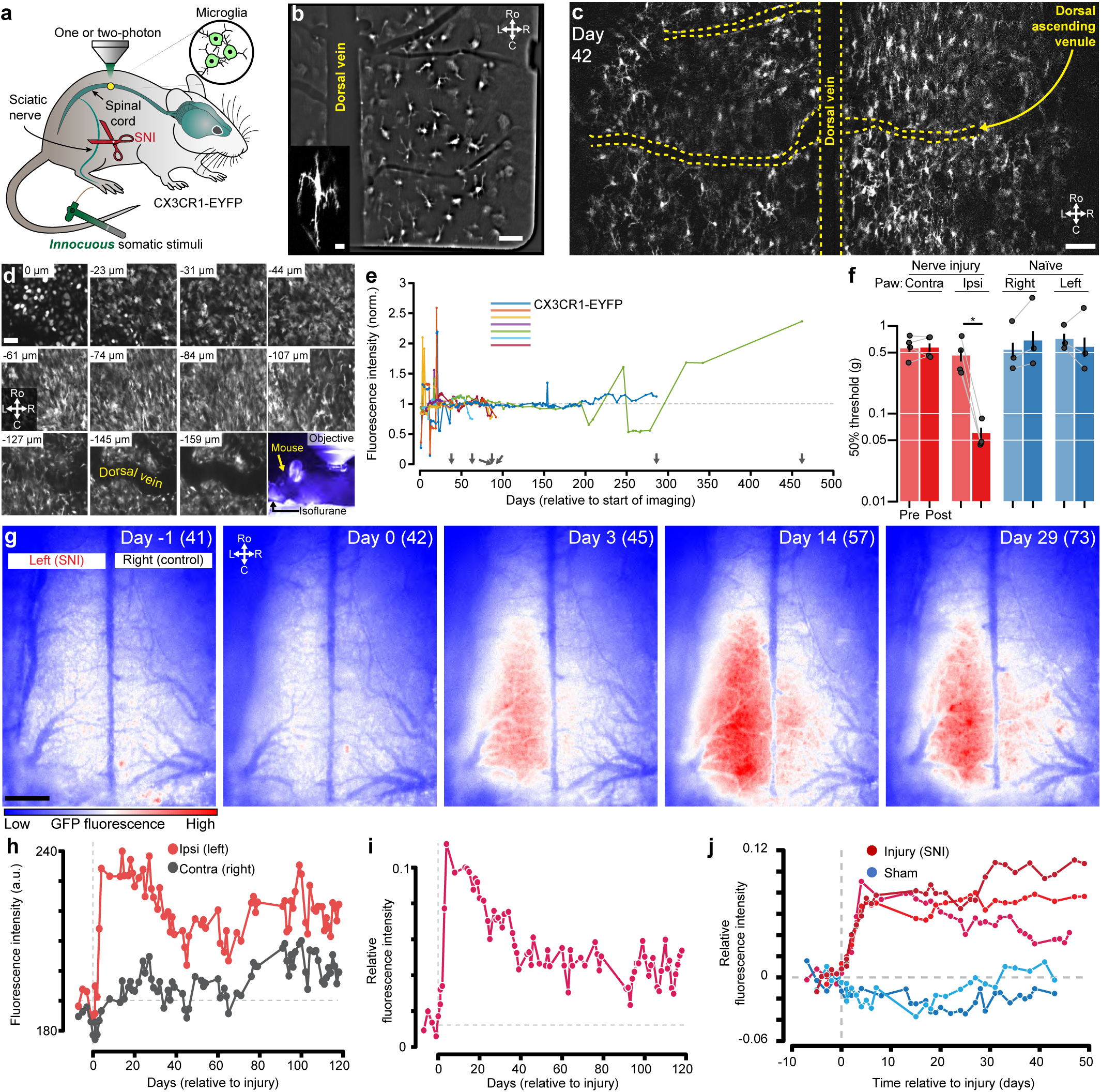
Long-term imaging of spinal cord microglia before and after injury in awake mice. **a.** Genetic strategy for longitudinal one- and two-photon imaging of microglia (CX3CR1-EYFP, as in Fig. 1q) before and after inducing a neuropathic pain model (spared nerve injury (SNI); (Corder et al., 2019)). **b.** One-photon mean projection image (bandpass filtered) of EYFP+ spinal cord microglia in an acute spinal preparation. Inset, higher magnification two-photon image shows the cell body and processes of a single microglial cell from the same session. Scale bar, 50 and 20 (inset) µm. **c.** Two-photon fluorescence imaging (920 nm, green channel) of EYFP+ microglia in the spinal cord of an awake mouse. Shown is a single montage plane from an imaging session 42 days post-window placement. Scale bar, 50 µm. **d.** Multi-plane, two-photon imaging of microglia (spinal cord) and monocytes (meninges) in an anesthetized mouse (2% isoflurane) spanning the meninges (top-left) into the spinal cord (bottom-right). Each image is the mean image of 10 time points. Depth is relative to meninges. Scale bar, 50 µm. **e.** Fluorescence intensity of the spinal cord parenchyma across individual one-photon imaging sessions spanning months from CX3CR1-EYFP mice (n = 7) normalized as in Fig. 3f. **f.** Mechanical sensitivity (50% withdrawal threshold, von Frey method) of naïve (n = 3 mice) or injured (n = 4) mice in **g-j**. Each dot is the mean of each animal’s thresholds before (Pre) or after (Post) surgery (SNI) or an anesthesia-only procedure (naïve). Bar plot and error are mean ± SD. Von Frey testing confirms that the SNI mice are hypersensitive on the injured side. **g.** One-photon fluorescence imaging of spinal cord microglia GFP intensity before (day -1) and after (days 0, 3, 14, and 29) left-side SNI model shows increased ipsilateral fluorescence. Numbers in parentheses are the imaging session days relative to the start of imaging of this mouse. Note bilateral fluorescence increases at later time points post-SNI. Scale bar, 300 µm. **h.** Absolute fluorescence intensity (a.u., imaging acquisition parameters held constant), of each side of the spinal cord, before and after injury in a CX3CR1-EYFP mouse as in **g**. **i.** Relative GFP fluorescence intensity (same mouse as in **g**-**h**) is defined as (I-C)/(I+C) where I and C are the mean ipsi- and contralateral side (relative to injury) fluorescence intensity of the spinal cord on that day. **j.** Relative intensity, same as in **i**, for the larger cohort of injured (n = 3) and sham (n = 2) mice.

#### Cross-session motion correction across a variety of experimental conditions

To compare expression across imaging sessions of spinal cord axons (Thy1-GFP mice, **Fig. 3a-h**), dorsal horn neurons and primary sensory axons (AAV-mediated GFP and tdTomato expression, **Fig. 3i-m**), and microglia (CX3CR1-EYFP mice, **Fig. 6**), we developed a multi-step motion correction protocol. This protocol involved manual correction for large shifts, rotations, and other FOV changes followed by multiple rounds of rigid registration, which led to significantly improved alignment (**Fig. 3e**). We also employed a modified LD-MCM protocol to track individual features, including vasculature and axons, across months to over a year (**Supplementary Video 10**).

### Long-term optical access in behaving animals

#### Spinal imaging setup for long-term imaging in behaving animals

Animal attachment to the imaging set-up typically takes less than a minute, aided by the use of clamps (**Fig. 4c**) and the fact that mice do not need to be anesthetized. To reduce the strain of lateral movements by the animal on the implant, while maintaining camera visibility of mice, we placed the mice in an imaging apparatus with infrared acrylic blinders (**Extended Data Fig. 9a**). To maximize the signal-to-noise (SNR) while maintaining an exposure of ≤10 ms, which minimizes motion blur (**Extended Data Fig. 7b**) and thus improves motion correction, we optimized several aspects of the preparation. We found that sCMOS cameras with >95% quantum efficiency (λ=500-600) and read noise <1.4 e^-^ (e.g. Photometrics Kinetix) along with low magnification/higher NA objectives (e.g. Fluar 5x/0.25NA) helped achieve this goal. Multiple cameras positioned around the running wheel provided a concurrent recording of behavior (**Fig. 5d**). Notably, our preparation is also compatible with spinal cord imaging in freely moving animals using miniature microscopes (**Extended Data Fig. 9e-j**). Further details are provided in the **Methods** and certain aspects (e.g. imaging and behavior camera synchronization to stimulus delivery) are comparable to our previously described awake animal imaging apparatus (Ahanonu and Corder, 2022).

#### Long-term Thy1-GFP axonal imaging demonstrates the persistence of spinal cord clarity

Absent injury, long ascending and descending axons should be intact. Therefore, to demonstrate long-term optical clarity, we recorded axonal GFP expression in Thy1-GFP mice for months to over a year (**Fig. 3a-c, Supplementary Video 10**). During these long-term recordings, in some mice the spinal cord would shift tens to hundreds of microns and stay stable around the new position, regardless of animal motion on a given day (**Fig. 3d**). To correct for these shifts we used CS-MCM and confirmed stable image quality and fluorescence intensity with one or two-photon imaging up to 1.5 years (557 days) of imaging (**Fig. 3e-f, Extended Data Fig. 7c**). Repeated awake imaging of the same animal is stable across dozens to hundreds of recording sessions (>260 sessions, **Extended Data Fig 7a**).

#### Longitudinal observation of virally-mediated gene expression

Long-term awake imaging of the spinal cord opens up the possibility of monitoring protein expression over time— including changes after injury and during disease progression—without the confounds of anesthesia and with detailed spatiotemporal information. Here, we monitored green (nuclearly-localized GFP) and red (tdTomato) fluorescent proteins after retro-orbital injection of AAV-PHP.eB-CAG-nls-GFP and AAV-PHP.S-CAG-tdTomato (**Fig. 3i-m**), which preferentially targets central and peripheral nervous systems, respectively (Chan et al., 2017). We found that GFP expression was upregulated in the dorsal horn, with less expression in the dorsal columns, consistent with dorsal horn-derived, rather than primary sensory neuronal expression. Two-photon imaging verified nuclear expression of the construct (**Fig. 3k**). TdTomato expression also increased over time, including in presumed primary sensory axons in the dorsal columns (**Fig. 3l, Extended Data Fig. 7d-e, Supplementary Video 11**).

#### Anesthesia-induced changes in vasculature

Anesthesia alters vascular dynamics in addition to neural activity. For example, isoflurane causes vasodilation (Schwinn et al., 1990; Matta et al., 1999; Constantinides and Murphy, 2016). Dilation of the midline spinal cord dorsal vein and dAVs could alter optical access to underlying grey and white matter. To compare the influence of anesthesia on dorsal vein diameter, we imaged the spinal cord under both anesthesia and awake, behaving conditions (**Fig. 3n**). The dorsal vein diameter was larger under isoflurane anesthesia (**Fig. 3o**, **Extended Data Fig. 8b**). Further, the detection of axonal GFP fluorescence was consistently reduced under isoflurane compared to the awake state sessions. When imaging the anesthetized and awake states in the same session, we observed that isoflurane induced a rapid and smooth reduction in fluorescence intensity, which correlated with the increased vascular diameter (R^2^ = 0.98 during induction of general anesthesia, **Fig. 3p**, **Extended Data Fig. 8c-e**, **Supplementary Video 12**). Allowing the animal to wake up led to a gradual return of the fluorescence towards baseline. Interestingly, two-photon imaging in the awake state revealed additional Thy1-GFP axons, not detected during anesthesia (**Fig. 3q**).

#### Spinal cord somatotopic maps revealed using large area neuronal Ca^2+^ imaging

To directly assess the topographic organization of inputs into the lumbar enlargement (Swett and Woolf, 1985; Takahashi et al., 2002, 2003; Odagaki et al., 2019), we took advantage of the large FOV provided by the spinal window (**Fig. 4d**). We assessed somatotopy using bulk Ca^2+^ imaging by labeling neurons throughout the lumbosacral spinal cord dorsal horn in transgenic animals after intraspinal injection of a retrograde virus (AAVretro-hSyn-Cre), which drove Cre-dependent tdTomato (Ai9) and GCaMP6s (Ai162) expression (**Fig. 4a-b**). Mice showed consistent behavioral responses to tail stimulation but at times we observed significant Ca^2+^ transients in the absence of stimulus-induced locomotion (**Fig. 4e-g**). We observed a somatotopic organization of the Ca^2+^ imaging responses to tail and hindlimb stimulation, with a rostrocaudal and mediolateral orientation that is consistent with termination zones of primary mechanosensitive afferents (**Fig. 4h-k, Supplementary Video 13**). This approach should prove of great interest for future studies of the reorganization of afferent inputs post injury (Merrill and Wall, 1972; Basbaum and Wall, 1976; Li and Zhuo, 1998).

#### Dorsal horn neural activity in awake, behaving animals

Spinal imaging in awake mice will provide insight to many unanswered questions about the code that the spinal cord projection neurons (SCPNs) transmit to the brain. We gained access to SCPNs using a transgenic approach combining the SCPN-biased Cre-driver line (Phox2a-Cre, (Roome et al., 2020)) and a Cre-dependent GCaMP6s line (Ai162, (Daigle et al., 2018)) (**Fig. 5a**). This led to expression within SCPNs throughout Lamina I of the dorsal horn (**Fig. 5b,** **Extended Data Fig. 3d**). Using one-photon microscopy, we consistently imaged these superficially-located neurons for months (n = 6 mice, **Fig. 5c**). By applying different noxious stimuli we could address the longstanding question about the polymodality of lamina I SCPNs. We concurrently monitored animal behavior and then identified individual body parts using DLC (**Extended Data Fig. 9b-d**) and processed GCaMP movies using CIAtah (Ahanonu and Corder, 2022). With this approach, we observed increased neuronal activity that coincided with escape-related behavior and/or head movements (**Fig. 5d, Supplementary Video 14**). As expected, the side of the spinal cord ipsilateral to the stimulated hindpaw showed increased activity relative to the contralateral side (**Fig. 5d-e**) and responses were consistent across repeated applications of the same noxious stimulus (**Extended Data Fig. 10a-c**). On multiple occasions, however, we observed SCPN activity contralateral to the stimulated hindpaw (**Fig. 5f**, **Extended Data Fig. 10d-f**). Interestingly, we also consistently found SCPNs activated during animal motion, whether or not we delivered a stimulus, an observation unexpected as lamina I SCPNs do not directly receive light touch or proprioceptive input in uninjured animals (**Fig. 5f, Extended Data Fig. 10d-f**). Occasionally, the mice exhibited minimal escape-like nocifensive behavior, yet we still observed robust SCPN activity. Tracking of individual animal body parts helped identify instances in which head movement without locomotion correlated with SCPN activity (**Fig. 5g**, bar #1). Importantly, our preparation allows monitoring of individual neurons over time (**Fig. 5h**) and we find that a subset of SCPNs consistently respond to the same stimulus across days (**Fig. 5i, Supplementary Video 15**). In contrast to consistent contralateral activity in the awake preparation, we observed only ipsilateral-side SCPN activity in anesthetized preparations. Further, we did not observe spontaneous activity in the anesthetized preparation, underscoring the importance of recordings in awake preparations (**Extended Data Fig. 10g-l**).

#### Long-term imaging of microglia before and after nerve injury

In the context of pain, spinal cord microglia undergo a number of transcriptomic and morphological changes in addition to proliferation (Guan et al., 2016; Ji et al., 2019; Donnelly et al., 2020). With our new preparation, we conducted long-term imaging of microglia using CX3CR1-EYFP mice (Parkhurst et al., 2013; Tan et al., 2022) and monitored changes before and after inducing neuropathic pain using the partial sciatic nerve injury (SNI) model (Shields et al., 2003; Corder et al., 2019) (**Fig. 6a**). The CX3CR1-EYFP line expresses in all monocytes, including microglia, in the meninges and spinal cord parenchyma. One- and two-photon *in vivo* imaging verified the morphological features of the different monocyte populations (**Fig. 6b-d, Supplementary Video 16**). Using one-photon imaging, we are able to track microglia expression across months (**Fig. 6e**). After inducing SNI, we observed persistent mechanical hypersensitivity in the injured cohort (**Fig. 6f**) and imaged experimental (n = 4) and control (n = 3) mice before and after the injury. In all cases, we show results relative to the ipsilateral and contralateral spinal cord. Consistent with cross sectional analyses in histochemical preparations after SNI, CX3CR1-EYFP fluorescence significantly increased ipsilateral to the injury **(Fig. 6g-h)**. We observed a persistent relative difference in EYFP intensity between sides over 4 months, with a clear increase ipsilateral to the nerve injury by 3 days post-SNI (**Fig. 6i-j, Supplementary Video 17**). Interestingly, over time fluorescence was also detected contralateral to the nerve injury. These results indicate that microglia can be studied before, during, and after chronic neuropathic pain develops, and with our preparation we can now study the utility of, for example, pharmacological interventions during the transition from acute to chronic pain.

## DISCUSSION

We have demonstrated surgical, experimental, and computational methods that overcome major limitations to long-term spinal cord recording in the awake, behaving mouse. By incorporating fluoropolymer membranes that provide long-term fibrosis regrowth inhibition, we reliably maintain clear optical access to the spinal cord, for many months. We predict that the use of these fluoropolymers (PRECLUDE and Teflon AF 2400) extends beyond the spinal cord, and can improve longevity and health of standard and whole cortex cranial windows. Our improved structural design provides an expansive view of the spinal cord, allowing concurrent monitoring of both sides within multiple segments of the lumbar enlargement. Furthermore, to handle the complex spinal cord motion that occurs during awake spinal cord imaging, we developed an image registration pipeline that combines deep-learning control point feature tracking, deformation-based non-rigid, and rigid methods. These algorithms are available within CIAtah, our existing Ca^2+^ imaging analysis pipeline (Corder et al., 2019).

Our design enables recording from the superficial dorsal horn, continuously, in the awake, behaving mouse, without repreparation (e.g. (Wu et al., 2022; Shekhtmeyster et al., 2023)). Further, while we focus on spinally-anchored animal recordings, our spinal chamber design is amenable for use with miniature microscope designs with longer working distances and larger fields of view (Scherrer et al., 2023). The latter designs will enable analyses in freely moving animals, broadening the repertoire of behaviors that we can correlate with spinal cord neural or glial activity. By monitoring the activity of the same population of neurons as well as non-neural cells before, during, and after injuries, we are now able to address longstanding questions about the transition from acute to chronic pain in the awake, behaving mouse. Previous studies were limited to cross-sectional analyses, in different mice at different times after injury. Of particular interest will be tests of the effects of novel analgesics (e.g., ɑ2a adrenergic receptor (Fink et al., 2022) and other agonists (Yekkirala et al., 2017)) across different stages of disease progression.

As expected, we found that isoflurane anesthesia reduces the excitability of dorsal horn neurons. Particularly notable was the appearance of spontaneous activity in the awake preparation and a greatly reduced response of neurons in the anesthetized state. Interestingly, neural activity contralateral to the side of stimulation in the awake state was readily apparent and clearly contrasted the minimal or non-existence of stimulus-evoked contralateral activity under anesthesia. How the emergence of these commissural neural dynamics in the awake state regulates somatosensation and whether these circuits are dysregulated in the chronic pain condition are unknown. Additionally, studies in the same mouse, with and without anesthesia, are now possible and will greatly expand our understanding of the basic physiology of spinal cord nociceptive processing, which previously, with very few exceptions, were performed in anesthetized animals.

Our focus on lamina I dorsal horn projection neurons is particularly significant as these neurons transmit information used by the brain to generate pain, itch, and other percepts. In awake mice, we recorded the properties of identified projection neurons within the Phox2A lineage and correlated their activity with evoked behavioral responses. Importantly, the Phox2A-expressing neurons constitute 50% of all dorsal horn projection neurons and display substantial overlap of Gpr83, Tacr1, and Tac1-expressing subpopulations (Alsulaiman et al., 2021). Of particular interest is whether so-called nociceptive specific neurons identified in anesthetized animals are, in fact, responsive to both innocuous and noxious stimuli in the awake state. Our initial findings are also relevant to the labeled line vs. patterning (population coding) question, namely, the extent to which projection neurons are polymodal or only respond to one input modality. In fact, we identified a heterogeneous population of lamina I projection neurons, including many responding to cold, heat, and noxious mechanical stimulation. The results presented here provide compelling *in vivo* evidence that supports previous recordings from *ex vivo* spinal preparations (Hachisuka et al., 2016; Warwick et al., 2022), intravital anesthetized imaging (Chisholm et al., 2021), and ribosomal profiling analysis (Wercberger et al., 2021). These studies collectively suggest that a significant population of dorsal horn projection neurons in superficial laminae are polymodal, responding not only to different modalities of pain-inducing stimuli but also to pruritogens that induce itch. To what extent the polymodality of the Phox2A-expressing population extends to the projection neurons that do not express this gene, remains to be determined. Lastly, advancements in optical techniques, combined with the reliability of our approach and flexibility of machine learning, also promise to allow reliable imaging and analyzing of cells in deeper layers of the spinal cord, such as by the use of three-photon imaging (Cheng et al., 2019), adaptive optics (Rodríguez et al., 2021), targeted illumination (Xiao et al., 2021), and their combination (Streich et al., 2021).

To address the complex spinal cord motion in the awake mouse, we developed a deep learning-based method that enables motion correction of large rostrocaudal shifts, even in the presence of overlapping objects, aided by the tracking of biological-relevant features in the imaging data. Feature detection was robust with minimal training data and could define fine branching of the dorsal horn vasculature, while ignoring out-of-focused structures on unwanted focal planes. Advances in zero-shot machine vision methods, such as foundational models similar to Segment Anything Model (Bommasani et al., 2021; Kirillov et al., 2023), offer the promise of even faster annotation with minimal user input. This can be combined with simultaneous multi-color imaging of fluorescent fiducial markers placed on the spinal cord during surgery and imaging at longer wavelengths, which have less occlusions, to conduct LD-MCM correction of GCaMP imaging videos. This will then allow improved cross-session identification of cells, e.g. by using iterative multi-day cell extraction and alignment approaches to identify additional cells or those with weak activity on a given day (Tasci, 2020). For non-rigid motion correction, future improvements can be introduced, including deformation-based methods that are more computationally efficient and also mass-preserving (Emond et al., 2020). Such modifications will be critical to maintain numerical accuracy across frames and allow us to reduce the need to restrict motion correction to rostrocaudal oriented displacement fields.

In summary, our methodological achievements showcase spinal projection neuron activity dynamics in tandem with complex behavioral measures and illustrate the post-injury activation of non-neuronal cells, namely microglia, which together contribute to chronic pain initiation and maintenance. These technological advances will illuminate the tissue and nerve injury-induced changes during the transition from acute to chronic pain. Notably, although applying a noxious stimulus normally evokes significant behavioral responses in the mouse and significant activity of the projection neurons, in several instances, we observed that the stimulus-evoked activity did not occur concomitantly with behavior. We presume that this unexpected dissociation of activity and behavior reflects a mix of the brain’s complex descending controls on the activity of dorsal horn neurons along with gating of nociceptive information within various nodes of the pain neuroaxis. Clearly, these findings would never be detected in anesthetized mice. To better understand such sensorimotor transformations across the nervous system, we anticipate studies combining analyses of spinal cord dorsal horn activity in behaving mice with what, to date, are more common brain imaging protocols.

## ACKNOWLEDGMENTS

We would like to thank the following colleagues for materials and assistance. Axel Nimmerjahn and Daniela Duarte for demonstrating their spinal cord setup. Amos Gottlieb and Random Technologies for generously gifting Teflon AF material. Dan Bernards and Tejal Desai for helping with the plasma treatment of Teflon AF. Youngho Seo and Ryan Tang for help conducting microCT experiments in the MicroPET/CT, MicroSPECT/CT, MicroCT, and Optical Imaging center. MicroCT experiments reported in this publication were supported in part by the Office of the Director, NIH under grant S10OD012301. DeLaine Larson for help optimizing one- and two-photon imaging. Eric Lam for help with, and advice on, machining and 3D printing. We thank the following people for reagents and mice. Donald McDonald and Peter Baluk, who provided low magnification objectives. Santos Puente and Isaac Delgado of VICI Metronics for merchandising Teflon AF. Wendy Xin and Jonah Chan for Thy1-YFP-H mice. Hua Su and Rich Liang for Thy1-GFP-M mice. Brian Roome for sending the Phox2a-Cre mouse line to the Basbaum lab. This work was supported by NIH NSR35097306 (A.B.), Open Philanthropy (A.B.), DARPA 9691 (A.B.), HHMI Hanna H. Gray Fellowship (B.A.), NIH R35 NINDS Supplement Funding (B.A.), NIH F32 5F32DE029384 (A.C.), Canadian Institutes of Health Research (PJT-162225, MOP-77556, PJT-153053, and PJT-159839) (A.K.), and NSF Graduate Research Fellowship 2034836 (M.R.C.).

## AUTHOR CONTRIBUTIONS

B.A., A.C., and A.B. designed the project and wrote the manuscript; A.C. and B.A built the instrumentation for surgery and imaging and developed the surgical and imaging protocols. A.C. and B.A. performed surgeries, histology, imaging, image processing, and data analysis; B.A. developed and tested the motion correction algorithms and performed animal behavior. M.RC. assisted with experiments and manuscript preparation, and created the supplementary surgery videos. A.K. provided the Phox2a-Cre mouse line.

## EXTENDED DATA FIGURES

**Extended Data Fig. 1.**
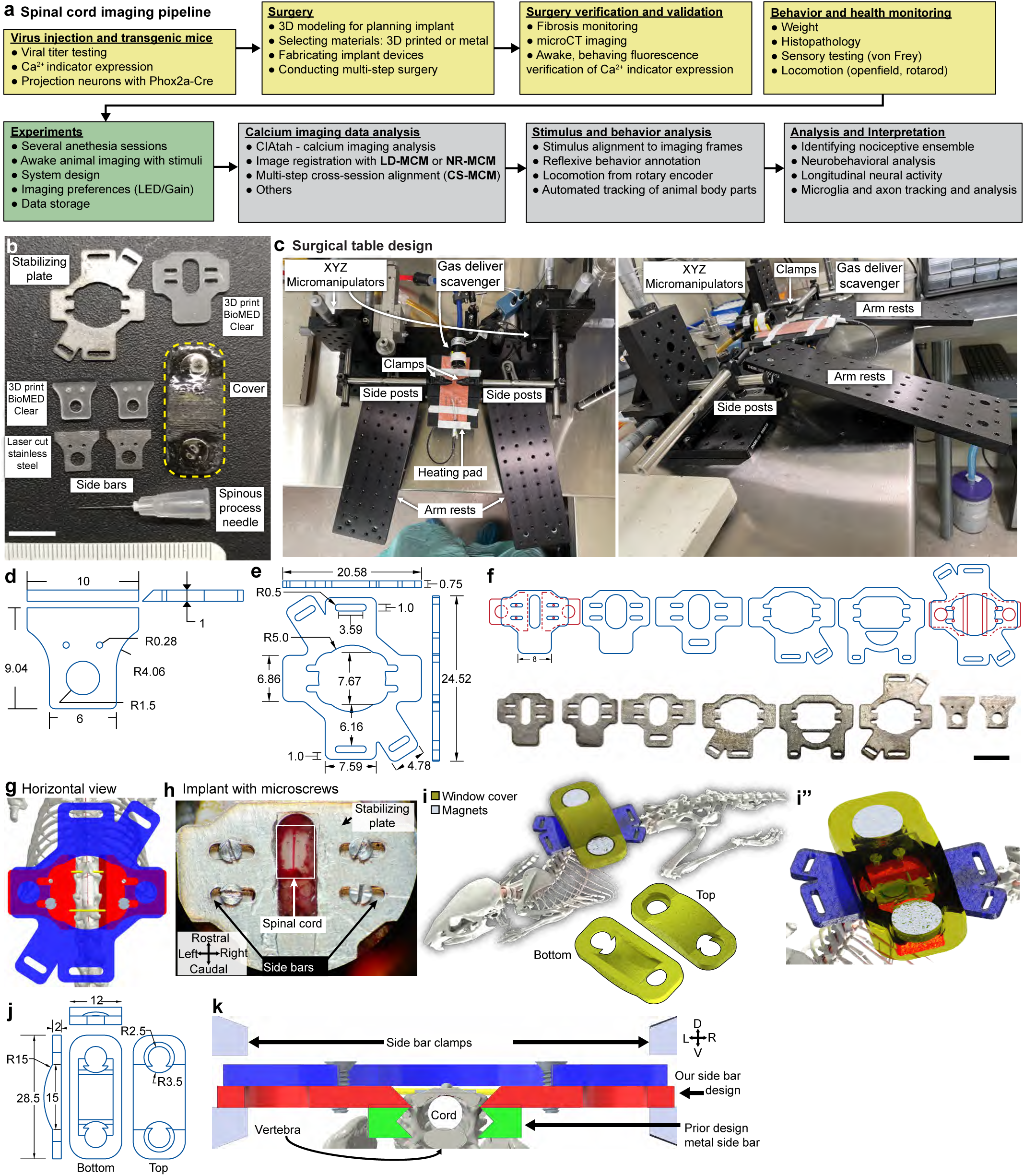
Spinal chamber experimental overview, fabrication, and designs. **a.** Spinal cord imaging workflow, including steps for surgery preparation, spinal chamber implantation, *in vivo* imaging, and data analysis. Several steps, such as microCT validation, are optional. **b.** Spinal implant chamber components. A set of both 3D printed and laser cut stainless steel side bars and stabilizing plates are shown. Each 33G needle is clipped during chamber implant surgery after its placement through the spinous process. Scale bar, 1 cm. **c.** Spinal cord surgery setup made from commercially available components and 3D printed parts, see **Supplementary Table 1** for a parts list. **d.** Side bars technical diagram with dimensions before and after (example taper angle) grinding during fabrication; units in mm. Scale bar, 1 cm. **e.** Stabilizing plate technical diagram with dimensions; units in mm. **f.** Several designs (top row, CAD; bottom row, real image) of the stabilizing plate that allows for different levels of cord exposure as well as locations where clamp can be applied or where the animal can be handled. Side bars included for size comparison. Scale bar, 1 cm. **g.** A 3D model showing a horizontal view of the spinal cord implant chamber and optional screws. **h.** Example of alternative design of spinal implant chamber (see **f**) with miniature screws. **i.** A 3D render of spinal cord chamber window cover for protection of implant. **j.** Technical diagram of side bar cover; units in mm. **k.** Coronal view of the present implant with dorsal oriented attachment to the T12-L1 vertebrae, compared to prior strategies. Note that our design allows standard clamps to manipulate the chamber during surgery and imaging. Colors for items: same as in Fig. 1a.

**Extended Data Fig. 2.**
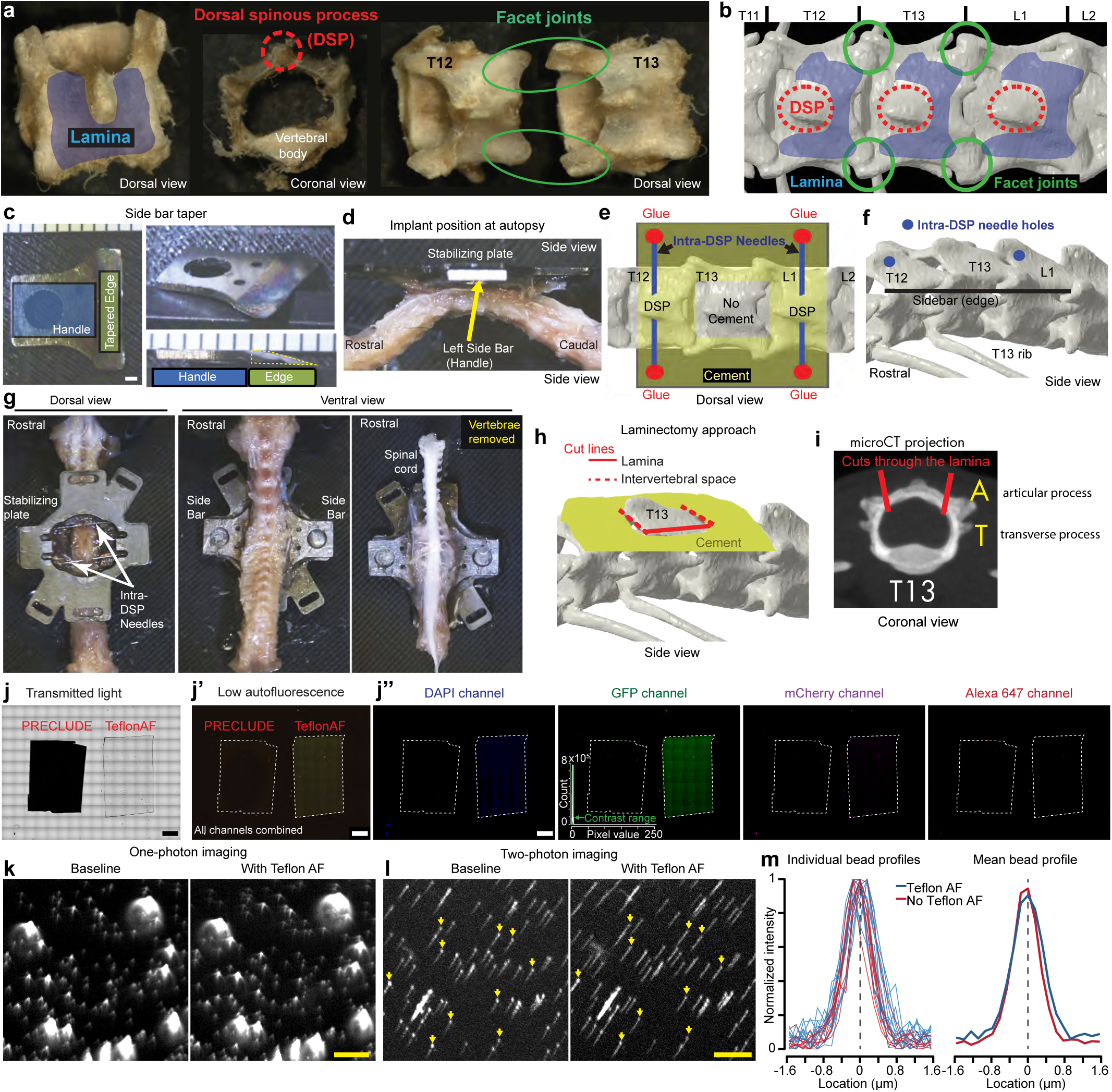
Blueprint of chamber implantation and fluoropolymer characterization. **a.** Highlighted vertebral anatomy critical to chamber implantation, including lamina (blue shading), dorsal spinous process (DSP, red circle), and facet joint (green circle). **b.** 3D rendering of the T12-L1 vertebrae of the spinal column visualized from above. Different orientations of the laminae, DSPs, and facet joints are highlighted. Note, when side bars are placed in the correct position, only the two circled facet joints are surgically exposed. They rest above the side bars. **c.** After laser-cutting of the side bar, and prior to implantation, the side bar edges are manually tapered by a grinding wheel. Scale bar, 1 mm. **d.** The taper allows room for the spinal process needles to be inserted and the side bar to be fixated above the T13 transverse process (the image shown is perpendicular to the spinal column). **e.** After the spinal process needles are superglued above the tapered side bars (red dots), the laminae, and DSPs of T12 and L1 serve as binding substrates for the dental cement. To allow for future laminectomy, the T13 lamina is kept free of cement. **f.** A view from the side shows where the spinal process needles bore through the DSP of T12 and L1 (solid blue circles) and bind to dental cement above the side bars. **g.** Spinal column dissection of a chamber-implanted animal. From a dorsal view, the stabilizing plate, the spinal process needles, and the T13 lamina are visible. When viewed from below, the bottom surface of the side bars is visible. After removing the vertebral bodies, the location of the spinal cord in relation to the placement of the side bars is confirmed to be centered on T13. **h.** Laminectomy removes the T13 lamina. As a >300 micron distance from the midline is necessary for the dorsal horn to be visible for imaging, lateral offset of the lamina cut is important. **i.** A microCT-generated image shows a cross-section of the T13 vertebra. Red lines indicate the lateral extent to which the T13 lamina is transected during laminectomy to access the spinal cord residing under the lamina in the spinal canal. **j.** Confocal microscope images of PRECLUDE and Teflon AF collected using transmitted light or lasers (405,488,561,640 nm) demonstrate transparency and minimal autofluorescence of Teflon AF. Brightness and contrast matched across the rightmost four images. Scale bar, 2 mm. **k.** Mean projection image from one-photon imaging of 1-µm yellow-green microspheres before and after placing Teflon AF on top of the microspheres; brightness and contrast matched. Scale bar, 20 µm. **l.** As for **k,** except two-photon imaging of the same microsphere slide. Arrows indicate beads used for measurements in **m**. Scale bar, 20 µm. **m.** Profile through 10 beads matched in two-photon imaging (as in **l**) with and without Teflon AF.

**Extended Data Fig. 3.**
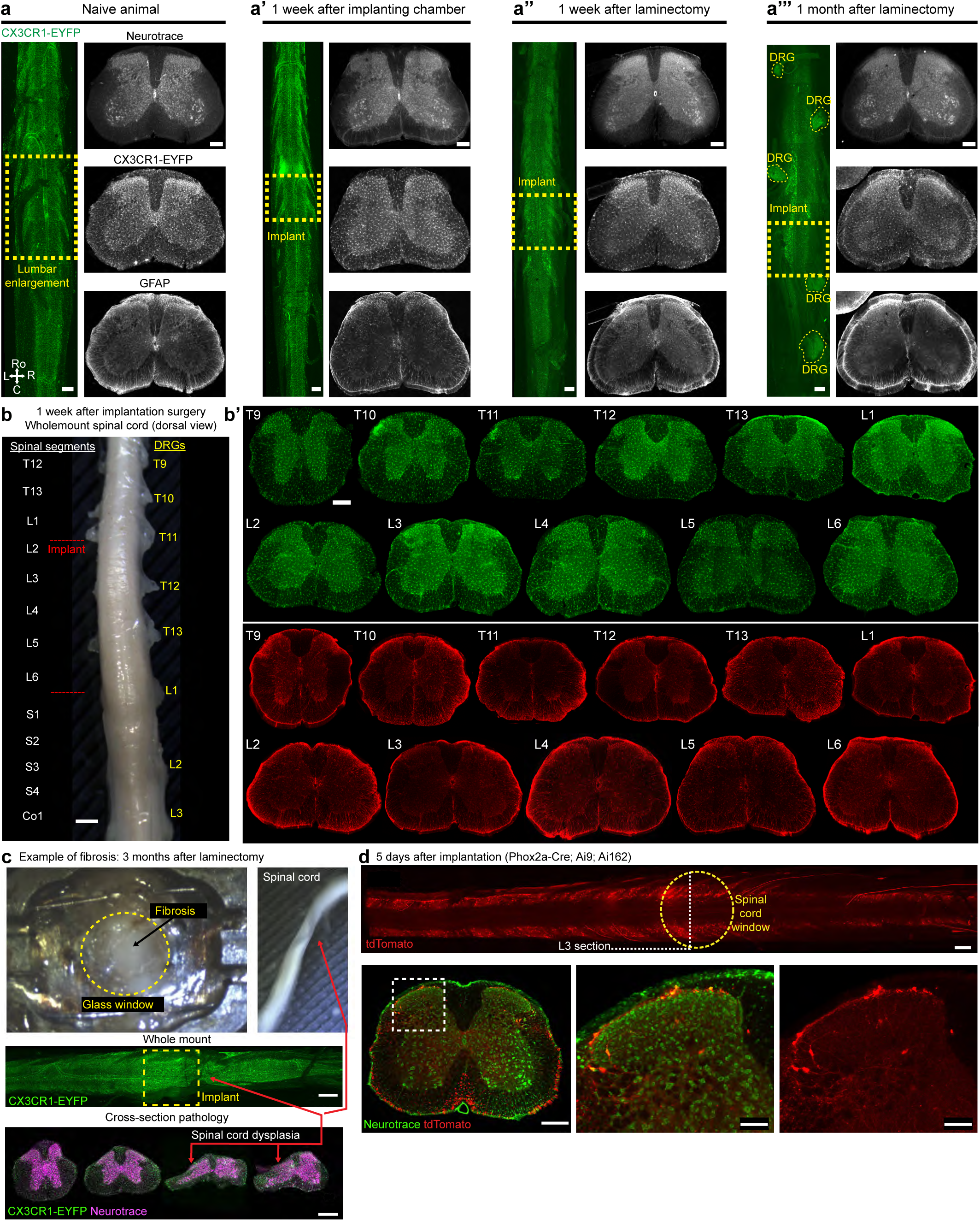
Histological analysis post chamber implant and laminectomy. **a.** Examples of naive and post-surgical CX3CR1-EYFP mice showing both EYFP in wholemount of the spinal cord and cross-section immunohistochemistry with Neurotrace and GFAP. Scale bars, 500 µm (wholemount) and 200 µm (coronal slices). **b.** Spinal cord dissection and histology of a CX3CR1-EYFP mouse 1 week after chamber implantation. The wholemount image (left) shows dorsal root ganglia in relation to the implant and the associated spinal segments. (b’) Cross-sectional views of EYFP (green) and GFAP staining (red) show a lack of gliosis near the implant. Scale bars, 1 mm (wholemount) and 200 µm (coronal slices). **c.** Post-mortem examination of a mouse that developed fibrosis after 3 months of imaging. A whitish spongy tissue filled the space beneath the coverslip, which displaced the spinal cord. Wholemount and cross-sectional histopathology show a large compression of the spinal cord at caudal levels of the implant. Scale bars, 1 mm (wholemount) and 500 µm (coronal slices). **d.** Spinal cord histology of a Phox2a-Cre; Ai162; Ai9 (LSL-tdTomato) mouse 5 days after chamber implant. Superficial lamina I projection neurons (tdTomato, red) are visualized by wholemount and a cross-section of L3 with a Neurotrace (green) counter-stain. tdTomato+ neurons are found in their appropriate laminae 1 and 5 locations. The dashed yellow circle shows the target location for the spinal cord window. Scale bars, 500 (wholemount), 300 (coronal slice), and 100 µm (zoomed coronal slices).

**Extended Data Fig. 4.**
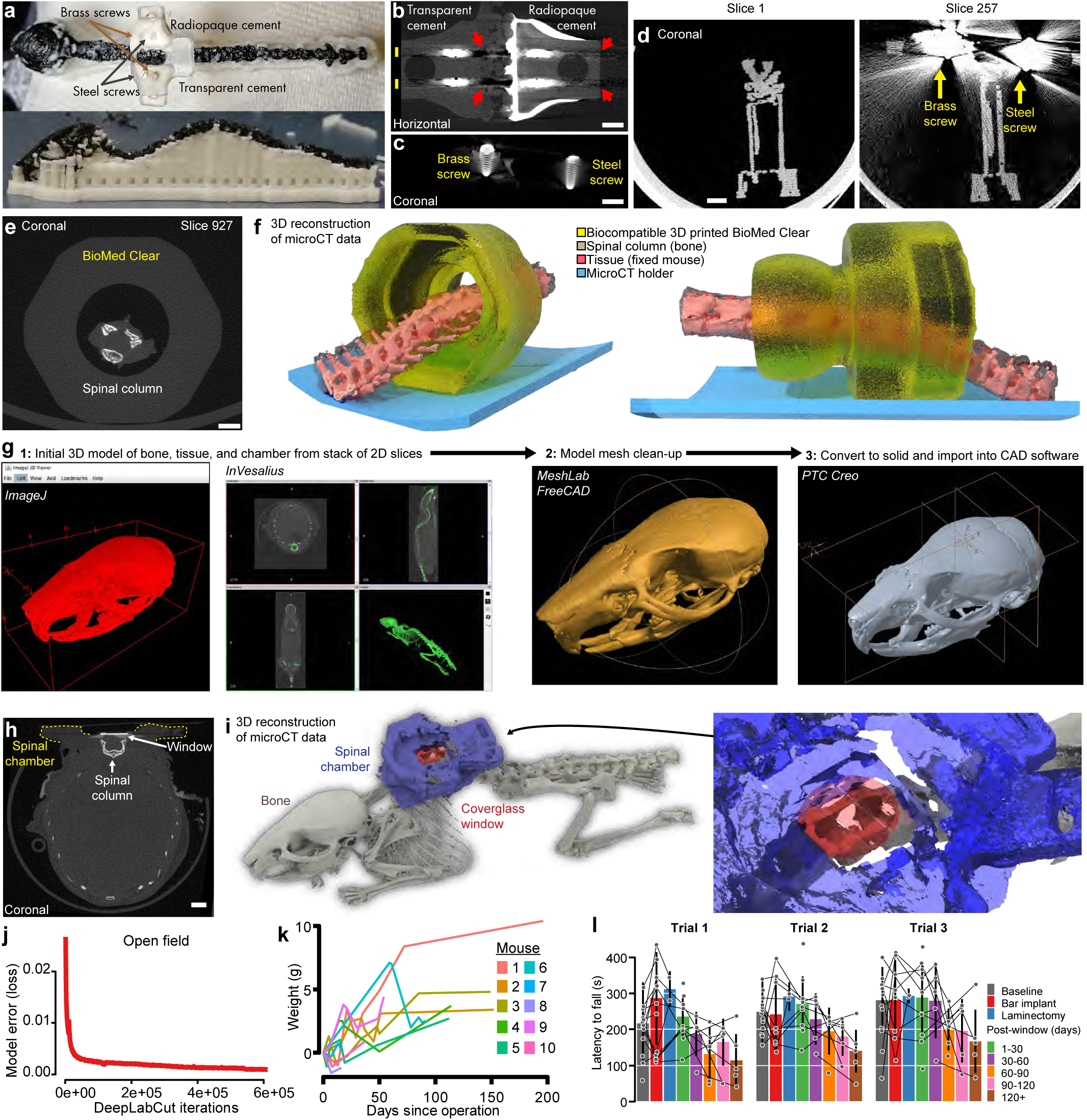
Validation of spinal implant with microCT, animal health, and behavior. **a.** 3D printed phantom, used for microCT validation studies, representing skull and spinal column. A 3D printed spinal chamber (Surgical Guide) is implanted with radio-transparent and -opaque dental cement along with miniature brass and steel screws (to evaluate impact on microCT scans). **b.** Reconstructed horizontal view from microCT scan of phantom in **a** of 3D printed side bars showing radiotransparent properties of the materials. Yellow bars indicate acquisition planes with reconstruction artifacts; red arrows highlight reduced reconstruction of spinal chamber and column. Scale bars, 2 mm. **c.** Coronal view of scan as in **b** shows details of metal screws along with artifact scan lines aligned with areas of higher and lower density of the metal screws. Scale bars, 2 mm. **d.** Coronal view through sections of the phantom without (left) and with (right) metal screws in the acquisition plane shows the artifacts introduced by miniature steel screws. Scale bars, 2 mm. **e.** Coronal section from microCT scan (20 μm resolution) of a dissected mouse spinal column, with tissue and muscle attached, placed inside a 3D printed test piece, using the same material (BioMED Clear) as for the 3D printed spinal chamber. Note the ability to reconstruct details and internal geometry of the spinal column, along with surrounding soft tissue. Scale bars, 2 mm. **f.** Off-axis and sagittal views of 3D reconstructed microCT scan as in **e**. Note the detailed reconstruction of the spinal column, soft tissue, and geometry of the test piece, confirming that BioMed Clear is microCT compatible. **g.** Pipeline for 3D reconstruction of microCT scans; see **Methods** for details. **h.** Coronal view of mouse with 3D printed spinal chamber (see Fig. 1g**-i**) showing an acquisition plane at the T13 laminectomy location. Scale bars, 2 mm. **i.** 3D reconstruction of the mouse in Figs. 1g-i and h with bone (gray), spinal chamber (blue), and circular glass coverslip window (red). Inset: zoomed in view highlights the T13 laminectomy and placement of the circular coverglass. **j.** Model error (sum of score map cross-entropy and body part location L1-distance losses) as a function of DeepLabCut iterations for model trained using data from 3 mice. Model is trained for 600,000 iterations until convergence. **k.** Weights of individual animals after bar implant. **l.** Mean (per animal) latency to fall in all three trials on an accelerating rotarod, comparing naïve (n = 14) and post-surgery mice, at different stages (n = 12, 2, 10, 5, 5, 5, 5, respectively).

**Extended Data Fig. 5.**
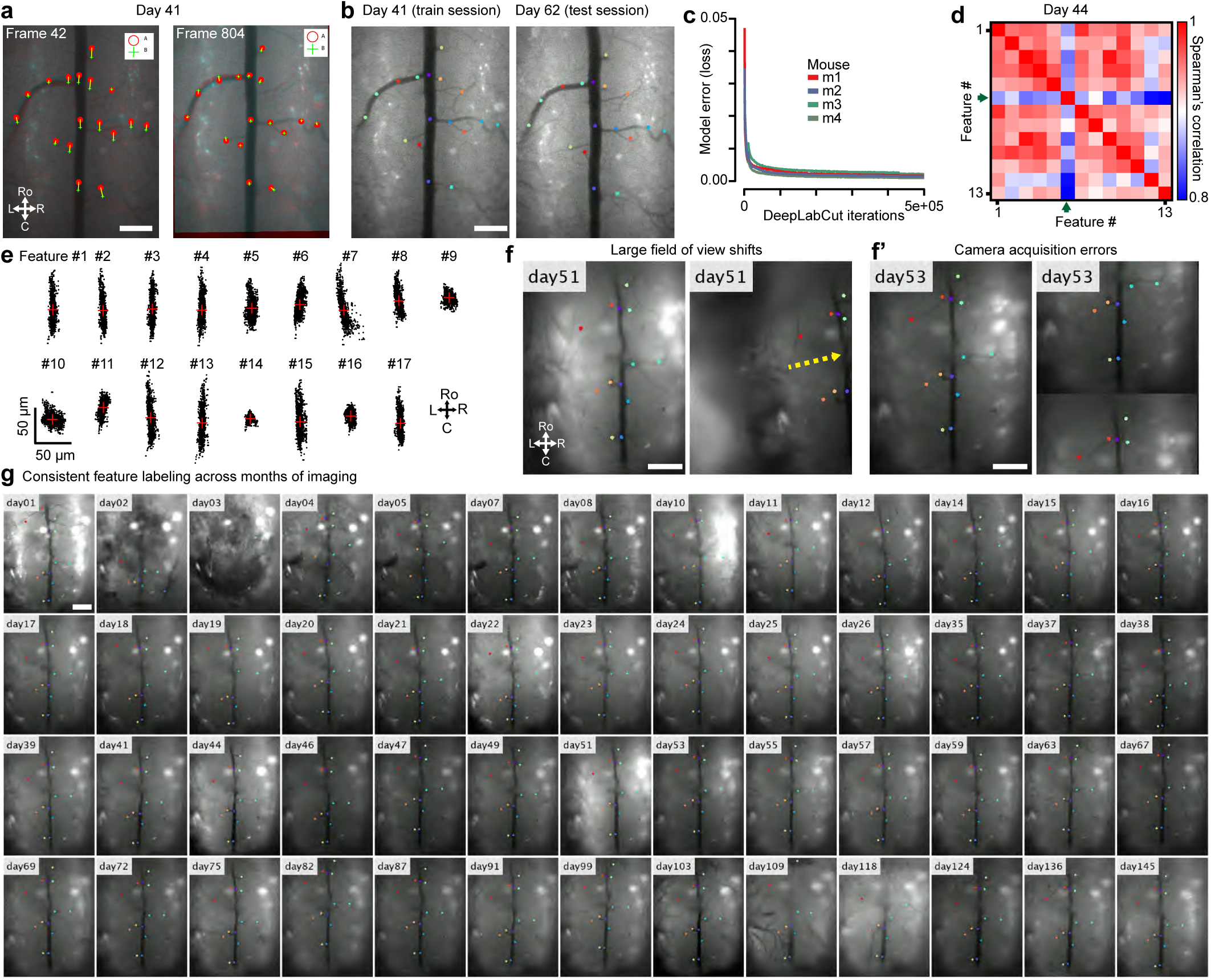
Deep-learning feature detection and control point motion correction. **a.** Comparison of reference frame 42 to movement frame 804 before and after control point motion correction. Scale bar, 300 µm. **b.** Example from a Phox2a-Cre; Ai162 mouse of DLC-identified vascular features used for cross-session registration with day 41 used for training the DLC model. Scale bar, 300 µm. **c.** Model error (sum of score map cross-entropy and body part location L1-distance losses) as a function of DeepLabCut iterations shows that performance approaches convergence by 500,000 iterations. **d.** Spearman’s correlation of each feature to other features in a movie from a Phox2a-Cre; Ai162 mouse. Green arrow indicates a feature that has reduced correlation with all other features and can thus be thrown out, improving overall motion correction. **e.** Point clouds for rostrocaudal and mediolateral movement of several features (from mouse in **a**) before motion correction. Each dot (2001 frames) represents the location of that feature on an individual frame during an imaging session (∼6 min, 13.9 Hz). **f.** DLC identification is consistent across both (i) large mediolateral shifts in the field of view not present in the training set and (ii) camera errors that result in a split of the field of view. These shifts can be used to rapidly exclude frames or movies in downstream analysis. Scale bar, 300 µm. **g.** Consistent DeepLapCut labeling of vascular features in a Phox2a-Cre; Ai162 (GCaMP6s) mouse across 52 neural activity imaging sessions, spanning nearly 5 months. DeepLabCut was trained using manual annotation from 20 frames from a single imaging session (day 75) demonstrating the robustness of annotation and consistency of imaging. Scale bar, 300 µm.

**Extended Data Fig. 6.**
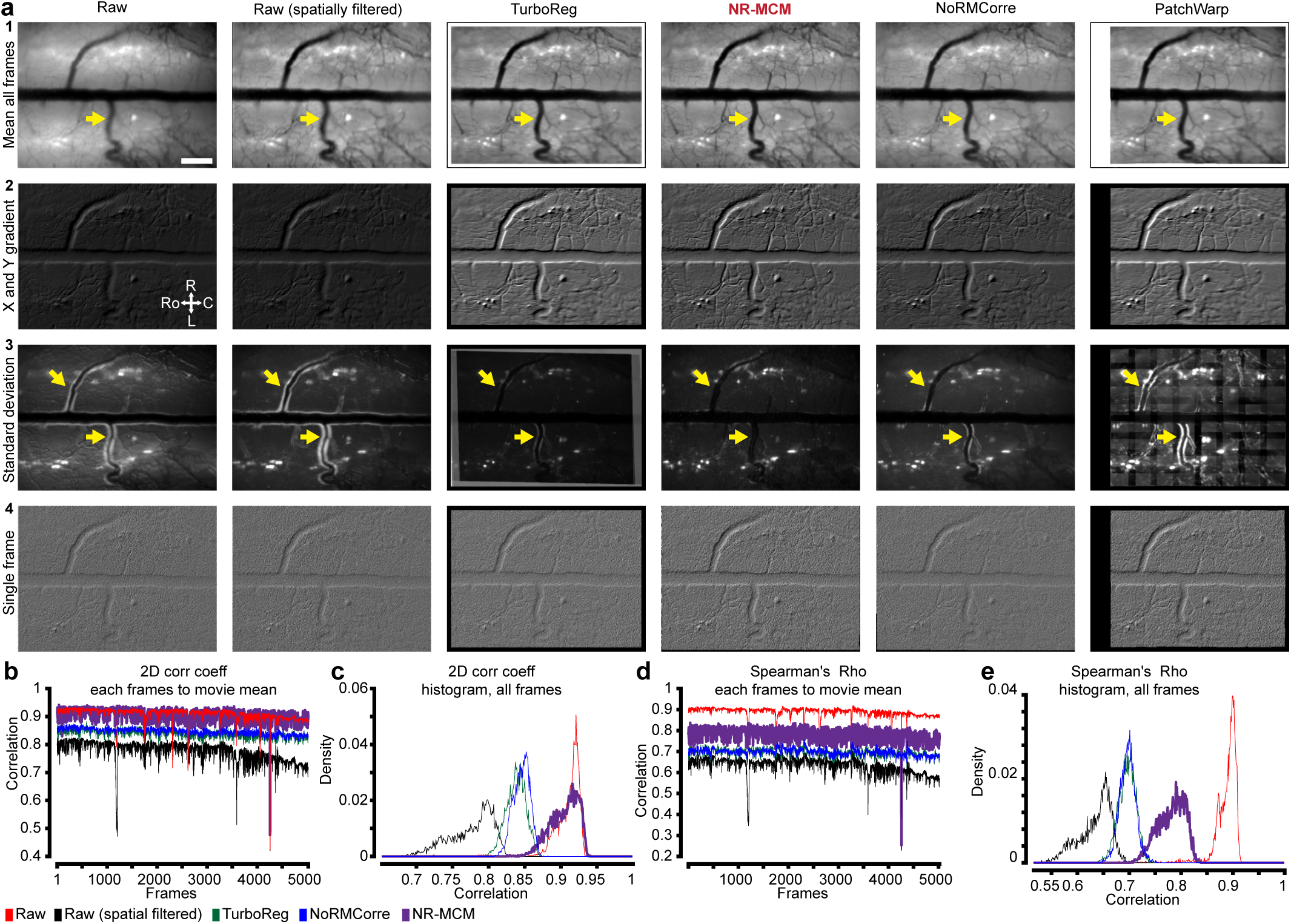
Deformation-based motion correction using displacement fields. **a.** Each motion correction method run on the movie (5,000 frames, 13.9 Hz) from a Phox2a-Cre; Ai162 (GCaMP6s) displays 1: the mean of all movie frames, 2: combined numerical gradient in both lateral directions on the mean frame, 3: the standard deviation over all movie frames (hence visibility of neurons on left and right side of the spinal cord), and 4: Δ*F*/*F* frames. The standard deviation and mean frames show the improved performance of NR-MCM compared to TurboReg or NoRMCorre. Arrows indicate areas of interest where differences between methods are most evident. **b.** 2D correlation coefficient of all frames to the mean frame of the movie (as in **a**) for NR-MCM compared to raw, TurboReg, and NoRMCorre. All movies (except raw) were spatially filtered to remove large magnitude, low-frequency changes in fluorescence, which artificially enhances correlations. **c.** Histogram of 2D correlation coefficients over all frames from **b**. **d.** Spearman’s rho of all frames to the mean frame of the movie (as in **a**) for NR-MCM compared to raw, TurboReg, and NoRMCorre. All movies (except raw) were spatially filtered to remove large magnitude, low-frequency changes in fluorescence, which artificially enhances correlations. **e.** Histogram of Spearman’s rho values over all frames from **d**.

**Extended Data Fig. 7.**
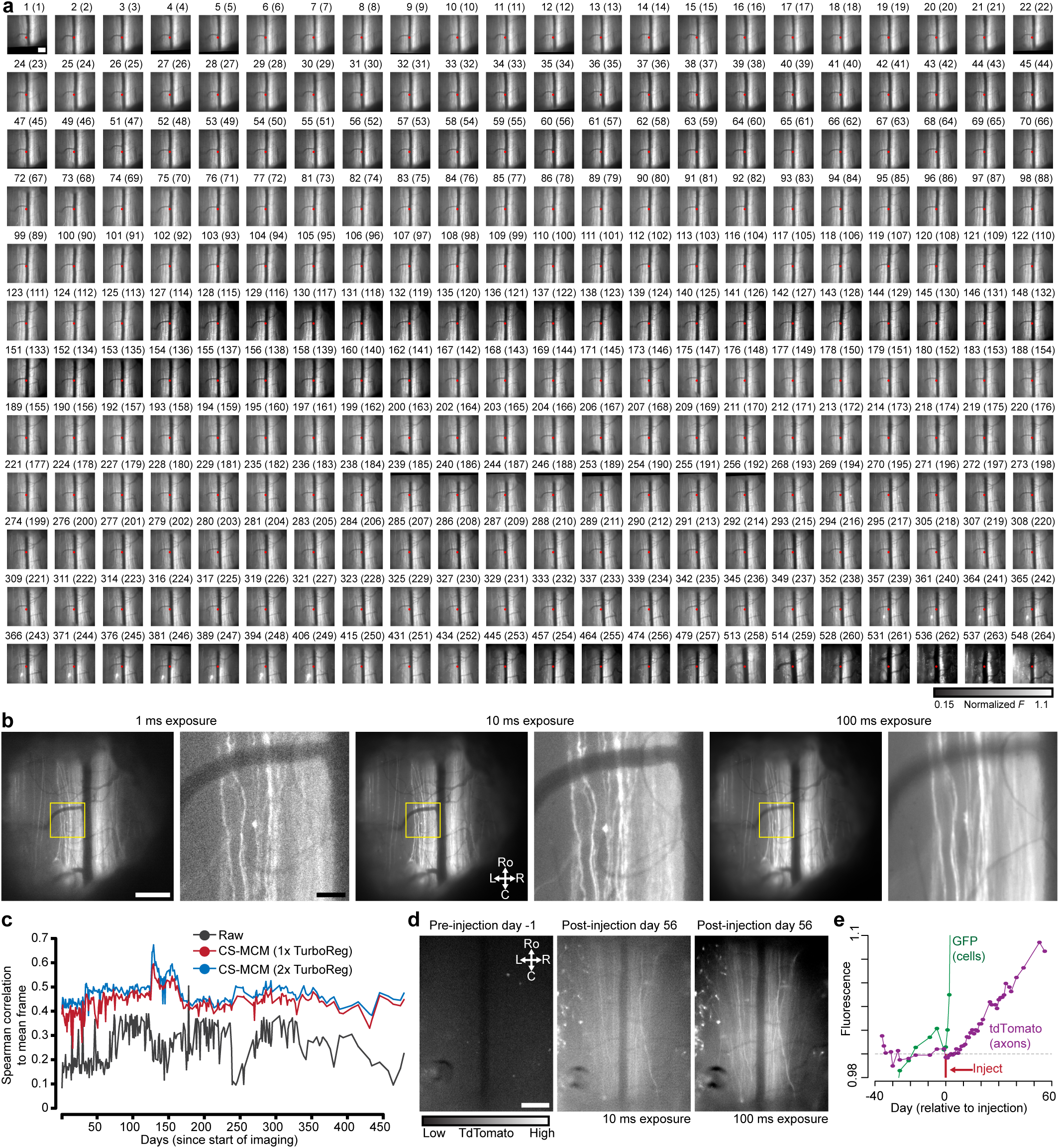
Long-term imaging of cell bodies and axons in the spinal cord of awake mice. **a.** Individual frames from 264 imaging sessions from the spinal cord of a Thy1-GFP mouse imaged for 548 days after cross-session alignment with CS-MCM. Red dot indicates the center of the field of view, near the intersection of the dorsal vein and a dorsal ascending venule. For display purposes, due to different cameras used during certain imaging sessions, here each frame is displayed between the 0.1th to 99.5th percentile of movie pixel values. Scale bar, 200 µm. **b.** Reduction in clarity of GFP+ axons (Thy1-GFP mouse) with increasing sCMOS exposure times (LED power held constant). As a trade-off between SNR and clarity, we used 10-20 ms exposure times. Scale bars, 300 and 50 µm. **c.** Spearman correlation to the mean frame of a raw movie from a Thy1-GFP mouse (as in **a**) showing improved cross-session alignment after CS-MCM. **d.** Increase in tdTomato expression in the dorsal columns after retro-orbital injection of AAV-PHP.S-tdTomato (as in Fig. 3i). Day 56, shows 10- and 100-ms exposure. Scale bar, 300 µm. **e.** Near daily imaging of GFP and tdTomato fluorescence normalized to baseline (pre retro-orbital injection). Zoomed in view of Fig. 3m highlights tdTomato signal increase from baseline.

**Extended Data Fig. 8.**
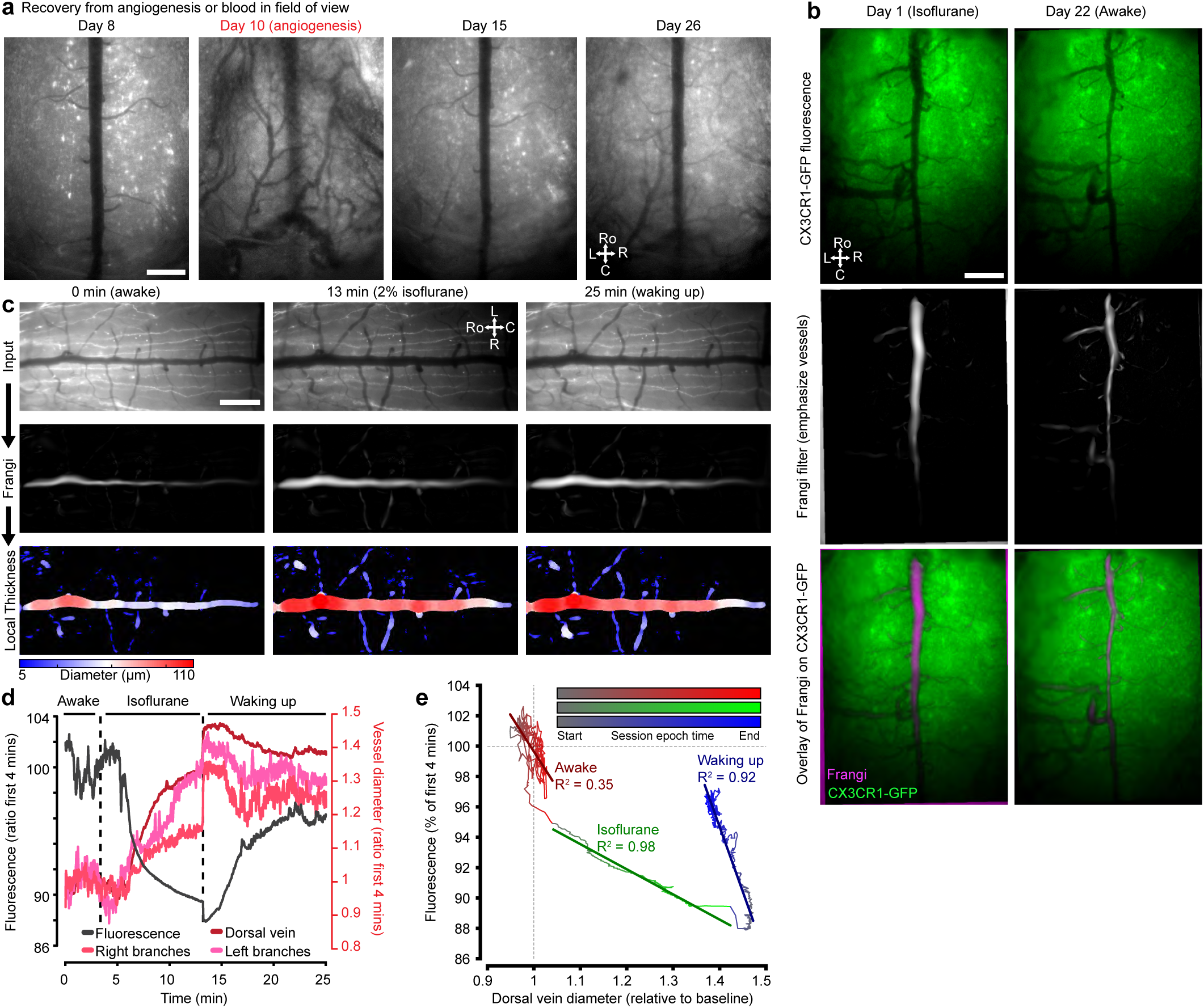
Transient angiogenesis and vascular dynamics in awake and anesthetized states. **a.** Individual frames across imaging sessions show onset and reversal of angiogenesis in the spinal cord of a CX3CR1-EYFP mouse. Scale bar, 300 µm. **b.** Change in spinal cord vessel diameter between general anesthesia and awake states in a CX3CR1-EYFP mouse. Middle row illustrates the same frames after application of a Hessian-based Frangi vesselness filter that highlights the dorsal vein and a subset of dorsal ascending venules. These filtered images are used to calculate changes in vessel diameter. Scale bar, 300 µm. **c.** Procedure for determining diameter of dorsal vein and ascending venules: a Frangi filter was applied to highlight vessels and their local thickness was then calculated to determine vessel diameter. Example frames are illustrated across three major behavioral states of a Thy1-GFP mouse during a 25 minute imaging session. Scale bar, 300 µm. **d.** Temporal change of vessel diameter and whole-frame fluorescence (normalized to 4-min awake baseline) within a single imaging session in a Thy1-GFP mouse before and after induction of general anesthesia (2% isoflurane). Same as Fig. 3p, but here additional right and left dorsal ascending venules are shown. **e.** Correlation of dorsal vein diameter and fluorescence during a 25 minute imaging session across several behavioral states: awake (red), induction and maintenance of general anesthesia (green, isoflurane 2%), and waking up (emergence) from general anesthesia (blue). First order polynomial best-fit lines and R^2^ indicated by darker colored lines and associated text, respectively.

**Extended Data Fig. 9.**
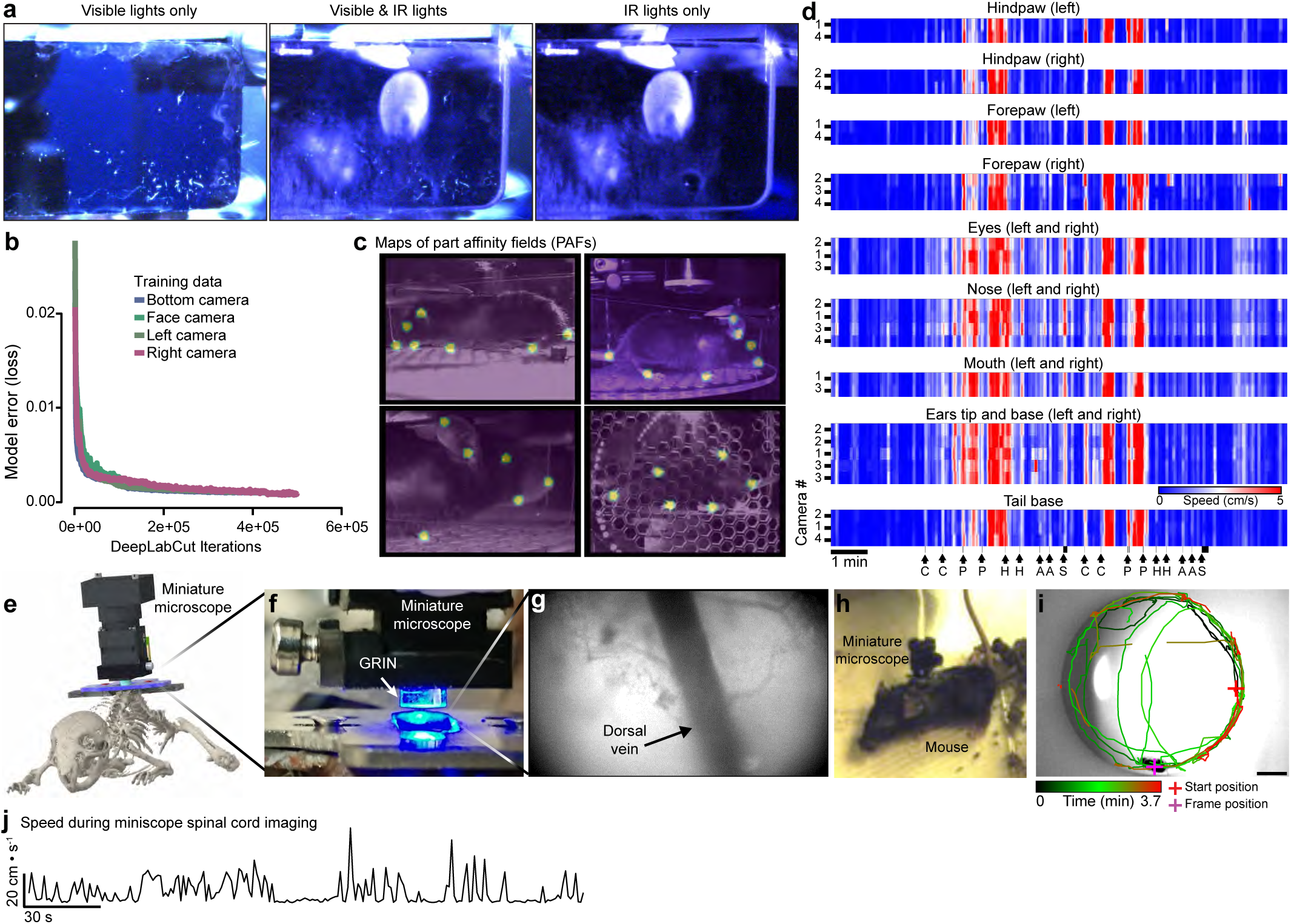
Behavior tracking and freely moving spinal cord imaging with a miniature microscope. **a.** Visibly opaque (black) infrared acrylic allows imaging of animal behavior using near-IR light sources and cameras, while blocking animal observation of experimenters (e.g. during stimulus delivery). **b.** Model error (sum of score map cross-entropy and body part location L1-distance losses) as a function of DeepLabCut iterations for model trained using data from one mouse. Model training is terminated after 500,000 iterations, when the loss asymptotes. **c.** Part affinity fields for DeepLabCut networks across multiple cameras. **d.** Speed of individual body parts across shows correlation of body part movement across cameras (#1-4). The mean speed across all cameras for each body part is used for display in Fig. 5g. Camera locations correspond to 1, left side of the body; 2, right side of the body; 3, right face; and 4, below the animal. Letters below each black arrow indicate the stimulus presented (C: cold; P: pinch; H; heat; A: air puff; S: sound); black bar denotes duration of the sound stimuli. **e.** 3D CAD of miniature microscope positioning above spinal implant chamber. **f.** Image of miniature microscope mounting. **g.** View of dorsal vein after procedure in **f**. **h.** Ambulating mouse after mounting procedure. **i.** General locomotion of a mouse in an open field during freely moving spinal cord imaging. Scale bar, 10 cm. **j.** Locomotor trace during the open field session in **i** (3.68 min, 10 Hz).

**Extended Data Fig. 10.**
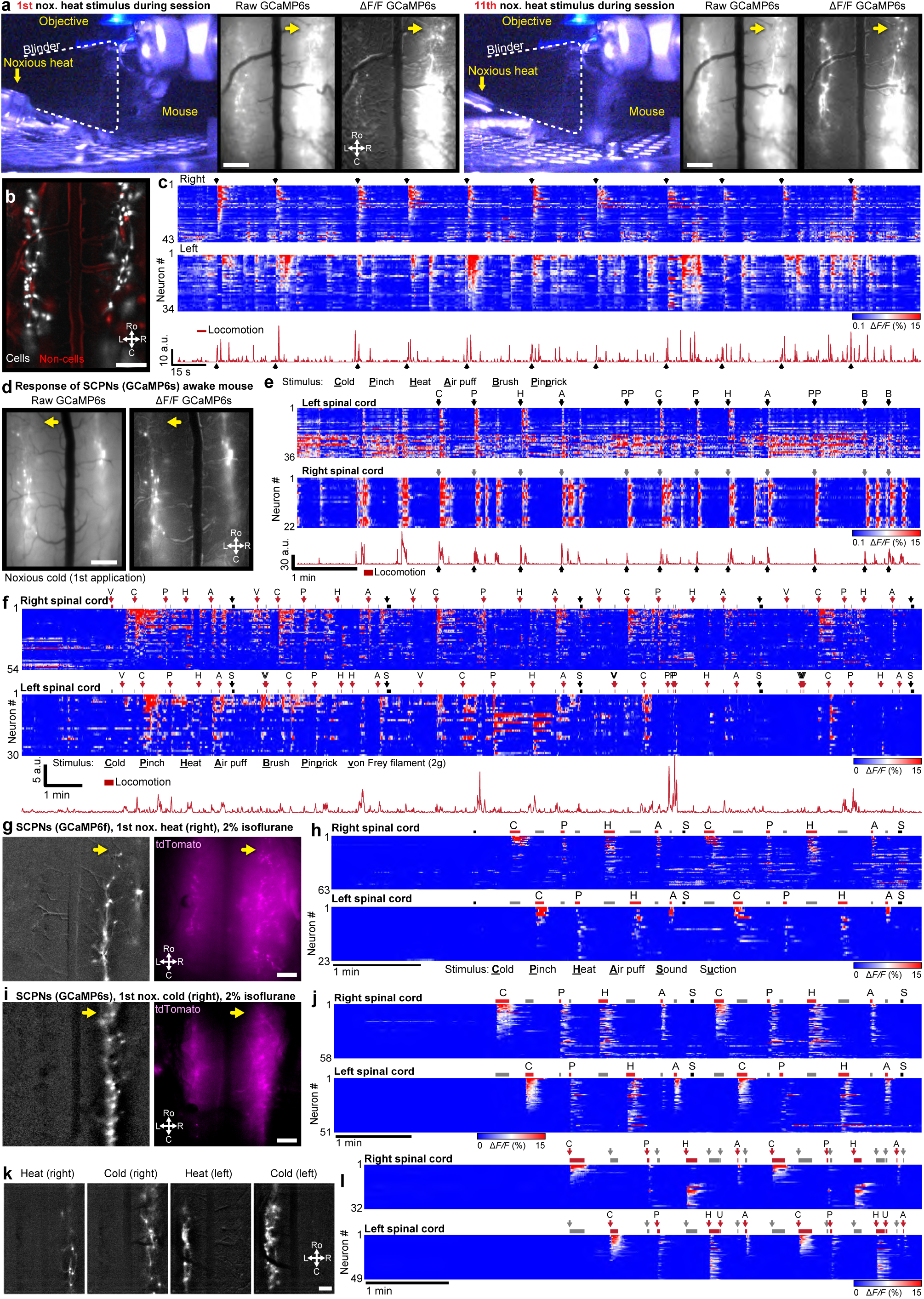
Imaging of spinal cord neuronal activity in awake and anesthetized animals. **a.** Consistency of noxious stimulus-evoked SCPN responses in Phox2a-Cre; Ai162 (GCaMP6s) after application of the 1st and 11th noxious heat stimulus, in the same imaging session. Shown are the behavior and associated raw and Δ*F/F* frames. Scale bar, 300 µm. **b.** Cell extraction outputs (CIAtah processing followed by PCA-ICA) shows cell (white, after manual sorting) compared to non-cell (red) outputs; the latter are excluded from further analysis. Scale bar, 300 µm. **c.** Activity of individual SCPNs (GCaMP Δ*F*/*F*), as in **a-b**, on the left or right spinal cord during a single imaging session (5.61 min, 13.9 Hz). Black arrows point to noxious heat applied to the right hindpaw. **d.** Raw and Δ*F/F* processed GCaMP frames from Phox2a-Cre; Ai162 (GCaMP6s) after hindpaw noxious stimulus. Scale bar, 300 µm. **e.** Activity of individual SCPNs (GCaMP Δ*F*/*F*), as in **d**, on the left or right spinal cord during a single imaging session (10 min, 9.75 Hz). Letters above each arrow denote the stimulus presented to the right hindpaw (C: cold; P: pinch; H; heat; A: air puff; PP: pin prick; B: brush). **f.** Extended recording session (25.47 min, 20 Hz) for the same mouse as in Fig. 5**d-g** shows SCPN stimulus-evoked activity in response to 5 blocks of stimulus applications. “V” is a 2-g von Frey filament. **g.** Δ*F/F* processed GCaMP and raw tdTomato frames from Phox2a-Cre; Ai148 (GCaMP6f); Ai9 (tdTomato) mouse under general anesthesia (2% isoflurane) shows overlap in expression. Scale bar, 300 µm. **h.** Activity of individual SCPNs (GCaMP Δ*F*/*F*), as in **g**, on the left and right spinal cord during a single imaging session (5.15 min, 20 Hz) during application of various noxious and non-noxious stimuli. There is a ∼2 min baseline period at the start of the session, prior to stimulus presentation. **i.** Same as **g,** except from a Phox2a-Cre; Ai162 (GCaMP6s); Ai9 (tdTomato). Scale bar, 300 µm. **j.** Same as **h,** but for the animal in **i,** during a single imaging session (7.74 min, 20 Hz). **k.** Same as **g,** except from a Phox2a-Cre; Ai162 (GCaMP6s) as in Fig. 5d**-g**. Scale bar, 300 µm. **l.** Same as **h,** but for the animal in **k,** during a single imaging session (6.72 min, 13.9 Hz).

## SUPPLEMENTARY VIDEOS

**Supplementary Video 1: Normal behavior of mice with implanted spinal cord chamber (Fig. 1j)**

Mice in homecage 40 days (green tape) and 50 days (yellow tape) after chamber implant are very active.

**Supplementary Video 2: Chamber implant (Fig. 1d)**

Step-by-step guide for attachment of the spinal chamber to the vertebral column; playback speed for each surgical step indicated on top left.

**Supplementary Video 3: Laminectomy (Fig. 1d)**

Step-by-step guide for laminectomy and placement of PRECLUDE to inhibit fibrosis; playback speed for each surgical step indicated on top left.

**Supplementary Video 4: Window placement (Fig. 1d)**

Step-by-step guide for PRECLUDE removal, Teflon AF application, and placement of circular glass coverslip window; playback speed for each surgical step indicated on top left.

**Supplementary Video 5: microCT sagittal slice (Fig. 1g-i)**

Sagittal slice through mouse with BioMed Clear spinal chamber implanted at T12-L1 and following laminectomy.

**Supplementary Video 6: microCT 3D reconstruction (Fig. 1i)**

3D rendering of microCT, post-laminectomy (T13), showing sequential removal of soft tissue (brown), spinal implant chamber (white), and glass window (red), leaving just the bone (gray).

**Supplementary Video 7: Open field tracking of mice with spinal implant (Fig. 1l-n)**

Open field tracking of behavior post-laminectomy for DeepLabCut network trained (top row) and network naive (bottom row) movies.

**Supplementary Video 8: Large shift motion correction using deep learning, control point registration, and rigid correction (Fig. 2c-f)**

Large rostrocaudal shift motion correction using LD-MCM compared to TurboReg and NorMCorre during spinal cord projection neuron recording in a Phox2a-Cre; Ai162 (GCaMP6s); Ai9 (tdTomato) mouse. Dots indicate features tracked using DeepLabCut in each movie; black bars in movies indicate sections of no usable data.

**Supplementary Video 9: Deformation correction using displacement fields (Fig. 2h-k)**

Non-rigid motion correction using NR-MCM compared to NoRMCorre and TurboReg, during spinal cord projection neuron recording in a Phox2a-Cre; Ai162 (GCaMP6s); Ai9 (tdTomato) mouse.

**Supplementary Video 10: Long-term Thy1-GFP spinal cord imaging, for over a year (Fig. 3c)**

Long-term imaging of a Thy1-GFP mouse after CS-MCM motion correction and with DeepLabCut tracking of vasculature features (colored dots).

**Supplementary Video 11: Time-lapse spinal cord expression of GFP and tdTomato after retro-orbital injection (Fig. 3i-m)**

Time-lapse imaging of increasing GFP and tdTomato expression before and after retro-orbital injection of AAV-PHP.eB-GFP and AAV-PHP.S-tdTomato.

**Supplementary Video 12: Rapid changes in fluorescence and vasculature upon isoflurane induction (Fig. 3p)**

Recording of change in vasculature diameter and Thy1-GFP fluorescence during a single session as the mouse enters and exits general anesthesia (2% isoflurane). Time on the bottom left is hours:minutes:seconds.

**Supplementary Video 13: Somatotopic mapping of spinal cord activity in response to caudal body stimulation (Fig. 4i-k)**

Bulk imaging of GCaMP6s activity after stimulation of indicated body parts from Ai162 mouse injected with AAV2retro-hSyn-Cre into the spinal cord.

**Supplementary Video 14: Imaging spinal cord projection neurons in response to various stimuli in the awake, behaving mouse (Fig. 5d-g)**

Awake spinal cord imaging session in response to noxious thermal and mechanical stimuli. Video shows spinal cord projection neuron activity (Phox2a-Cre; Ai162), behavior from multiple camera angles (colored squares indicate DLC tracking points), locomotion (white, rotary encoder), head movement (cyan), and stimulus application (red).

**Supplementary Video 15: Cross-day imaging of the same area of the dorsal horn in anesthetized and awake states, over a month (Fig. 5i)**

Recording of spinal cord projection neuron activity across 20 days, before and after applying noxious heat to the right hindpaw (red bar on left side of movie). Days 7-8 are under isoflurane anesthesia.

**Supplementary Video 16: Multiplane imaging of monocytes in a CX3CR1-EYFP mouse (Fig. 6d)**

Two-photon imaging of monocytes (CX3CR1-EYFP mouse under 2% isoflurane) showing multiple planes from the spinal cord parenchyma to the meninges during a ∼15-min session.

**Supplementary Video 17: Cross-day timelapse shows increased nerve injury-induced CX3CR1-mediated GFP microglia expression (Fig. 6g)**

Time-lapse imaging for months of microglia dynamics before and after nerve injury (left hindpaw, SNI model). Red regions indicate areas of greater fluorescence.

## METHODS

### Animals

We conducted all animal experiments in accordance with protocols approved by the University of California, San Francisco Institutional Animal Care and Use Committee (IACUC). We used the following mouse lines: Phox2a-Cre (Roome et al., 2020), provided by Artur Kania (Institut de Recherches Cliniques de Montréal); Ai162 floxed-GCAMP6s (#031562, JAX); C57BL6 (#000664, JAX); Ai148 floxed-GCaMP6f (#030328, JAX); CX3CR1-CreER-IRES-EYFP mice (Parkhurst et al., 2013; Tan et al., 2022), provided by Zhonghui Guan (UCSF); Thy1-YFP-H (Feng et al., 2000), provided by Jonah Chan (UCSF); and Thy1-GFP-M (Feng et al., 2000), provided by Rich Liang (UCSF). We studied both male and female mice.

### Viruses

The nomenclature for all AAVs used in this study refers to the AAV capsid. All AAVs contained the AAV2 ITR unless otherwise specified, e.g. AAV9 indicates AAV2/9 for AAV2 inverted terminal repeat (ITR) sequence and AAV9 capsid.

### Custom fabrication of spinal implant chamber components

To reduce cost and increase accessibility, we optimized the design (PTC Creo 6.0–9.0) and fabrication of the spinal implant chamber and used standard components and widely accessible fabrication technologies. We laser cut the side bars from multipurpose 304 stainless steel, purchased as 0.048’’/1.2192 mm (McMaster-Carr, Elmhurst, IL, 8983K114), 0.036”/0.9144 (McMaster, 8983K113), 0.024’’/0.6096 (McMaster, 8983K111), or 0.01’’/0.254 (McMaster, 3254K322) thick sheets. We predominantly used 0.036’’ (0.9144-mm) thick side bars. We outsourced custom metal laser cutting to Laser Alliance LLC (Milpitas, CA), using a laser beam kerf of 0.008" and the design files provided in the GitHub repository. Cost can come out to ∼$10 per side bar or stabilizing plate. Using the provided design (**Extended Data Fig. 1d-e**) and files, it is possible to use any laser cutting contractor after adjusting the dimensions to take into account the beam diameter. We then manually tapered the edges of the side bars with a grinding wheel (WEN, 4276 2.1-Amp 6-Inch) to a shallow angle (Extended Data **Fig. 2c**). Metal was then cleaned with a wire brush and ddH_2_0 and autoclaved at 250 °C for 20 minutes before implant. The tapered edge fits under the articular processes of the T12-L1 vertebra and rests against the vertebral wall. Note, if the angle of the side bars’ taper is not steep enough, then additional V-groove cuts can be made near the locations where the spinous process needle will be placed. We laser cut the stabilizing plate from 0.75-mm (∼1/32’’) thick mild steel, outsourced to Laser Alliance LLC using the design files provided in the GitHub repository. Because of its ferromagnetic properties, we used mild steel, which made it possible to attach magnetic devices, such as the protective cover. We designed iterations of the metal stabilizing plate for custom handling and clamping during imaging and to maximize the working area around vertebrae during surgery (**Extended Data Fig. 1f**). The stabilizing plate is critical; without it the side bars have a high chance of snapping off, due to the lack of proper load distribution when the animal moves while fixed in the imaging setup.

To protect the glass coverslip window from scratching and damage, we 3D printed covers (Stratasys, uPrint ABS and MakerBot, Nylon 12 Carbon Fiber [375-0061A] or ABS-R [375-0071A]) to which we attached two neodymium magnets (McMaster, 5862K141) using cyanoacrylate glue (Henkel, Loctite 4311) or optical adhesive (Norland Products, NOA81) (**Extended Data Fig. 1i-j**). The cover snapped on top of the ferromagnetic stabilizing plate, which we removed during imaging. We found this to be a faster and more reliable cover method than attaching the cover with micro screws or using a sliding press fit around the side bars (due to the size, both methods increase the probability that the cover will snap or fall off). Note, to prevent accidental removal of the spinal chamber or animal distress due to immobility, it is critical when using the magnetic protective covers or mild steel stabilizing plate (as opposed to 316 stainless steel), to check that cages and all devices that animals interact with are non-magnetic.

To perform microCT imaging, we 3D printed side bars and stabilizing plate with biocompatible non-metallic materials (FormLabs, BioMed Clear Resin) (**Extended Data Fig. 1b**, **Extended Data Fig. 4a-i**) using Formlabs 3D printer located at the KAVLI-PBBR Fabrication and Design Center. Due to the reduced tensile strength of BioMed Clear compared to steel, we designed thicker side bars (1.0- and 2.0-mm thick) and stabilizing plates (1.5- and 2.0-mm thick). This design improved the strength and reduced the probability that the pieces would snap during imaging. Furthermore, the stabilizing plate design includes chamfers that make it easier to remove the pieces from the build plate with a chisel without damaging or snapping them; the side bars can be printed upside down and removed by chiseling under the tapered edge. We also tested and found that Surgical Guide (FormLabs) can substitute for BioMed Clear when 3D printing non-metallic pieces.

We also manufactured side bars by metal 3D printing directly, using the STL models provided on the GitHub repository. This option bypasses the need to manually taper the side bars. This allows for more precise control over the final angle, the addition of V-grooves to accommodate spinous process needles, and other features customized for experimental conditions. We tested products from Protolabs, using direct metal stainless steel 316L (CL 20ES) with 20 micron layers, and i.materialise (Materialise NV, Leuven, Belgium) using Titanium and High-Detail Stainless Steel. We found both work well and additional options are available from other 3D metal printing manufacturers (e.g. shapeways and xometry).

To create the side bars, we tested using CNC machining (Tormach, PCNC 1100). However, we found that this process introduced additional requirements, such as manufacturing additional jigs and devices to hold the side bars during tapering. Due to the small size of the side bars and the heat produced during the machining process, this often led to part failures. Further, this process is less accessible and more costly. It is also possible to have the pieces manufactured entirely using CNC machining, which improves accessibility but is more costly per piece and introduces design limitations. These limitations are not present when laser cutting or 3D printing; we found these to be less expensive on a per-piece cost.

### Spinal chamber and window surgical procedures

#### First procedure: Surgical Implant

We began surgical procedures by inducing and maintaining mice with 2% isoflurane and administered a non-steroidal anti-inflammatory drug (NSAID), carprofen (5.0 mg/kg, subcutaneous injection). We depilated and disinfected a 1-cm × 1.5-cm area around the hump of the back (Betadine and 70-75% ethanol). We then applied sterile ointment (Alcon, SYSTANE®, White Petrolatum-Mineral Oil) to the eyes. We then transferred mice to a custom surgical table featuring bilateral side posts that are micro-manipulatable in three axes and lined up to the lumbar enlargement. We incised the skin above the T12-L1 vertebrae using surgical scissors (Fine Science Tools [FST], #14060-11 or #14060-10), followed by intra-incisional 0.5% lidocaine. We used a gelatin sponge (Ethicon, Surgifoam®) soaked in sterile saline as needed for hemostasis. We only cut the paraspinal muscle and fascia overlying each lamina, starting at the midline, and resected using micro scissors (#15023-10, FST). To visualize and separate the tendinous attachments to T13, we temporarily laterally retracted the incised muscle (RS-6504, Roboz).

We then scraped the dorsal surfaces of the T12-L1 laminae (FST, 10075-16) and wiped (Beaver-Visitec International, Cellulose spears, Weck-Cel®) them free of remaining connective tissue. Next, we inserted dorsal spinous process needles (Accuderm Inc., 33G–1/2”) at the T12 and L1 vertebrae. We positioned the side bars under the needles, under the T12-T13 & T13-L1 facet joints, and against the vertebral wall. We masked the intervertebral spaces using small amounts of Kwik-Sil (KWIK-SIL, WPI), and then sutured (Patterson Dental, 6-0 PGA #090-1660) and sealed (Vetbond, 3M) the skin rostral and caudal to the side bars. We sequentially glued the needles, side bars, and the stabilizing plate to each other using super glue (Henkel Adhesives, Loctite®, Ethyl 2-cyanoacrylate). In most circumstances, we pretreated the laminar surfaces with a dentin activator (Parkell, FeCl, C&B Metabond kit).

To secure the implant in place, we then cemented (Parkell, PMMA, C&B Metabond kit) the entire working area, except the T13 lamina, and covered the open area above the T13 lamina bone with Kwik-sil. We removed the animal from anesthesia and monitored it with supplemental heat during recovery. We administered one dose of sustained-release buprenorphine, Ethiqa (3.25 mg/kg), by subcutaneous injection when the mice awoke. When fully ambulatory, we returned the mice to their home cage. The next day, we gave a second and third dose of carprofen (5.0 mg/kg) by subcutaneous injection. On day 3, we re-dose with the carprofen if necessary. We monitored the health of the mice twice daily for signs of lethargy, immobility, poor grooming, or weight loss. Mice with implants recovered from surgery for at least one week before proceeding to the laminectomy.

#### Second procedure: Laminectomy and PRECLUDE (opaque Teflon) application

Under 2% isoflurane anesthesia, we performed a T13 laminectomy using micro scissors (FST, #15010-09) or a crescent blade (FST, #10317-14). If required, we used a high-speed bone drill (Foredom, K.1070) to clear cement over the T13 laminectomy area. After laminectomy, we immediately covered the exposed spinal cord with a gelatin sponge (Ethicon, Surgifoam®) soaked in sterile saline. Using dual fine forceps, we pulled any remaining dura laterally to expose the leptomeninges. We inhibit fibrosis with GORE® PRECLUDE® Pericardial Membrane, which we purchased from a third-party medical vendor (dotmed.com). Next, we cut the GORE® PRECLUDE® Pericardial membrane to the size of the laminectomy opening using micro scissors and placed it directly over the exposed spinal cord. To limit ingrowth, we placed small pieces of Surgifoam around the membrane. We blotted these small pieces dry with an absorbent spear (Beaver-Visitec International, Weck-Cel®). Next, to protect and hold the PRECLUDE® membrane in place, we spread Kwik-Sil over the wider working area. We removed animals from anesthesia and monitored them closely post-op until they ambulated. We administered one dose of carprofen (5.0 mg/kg, sc) for postoperative analgesia. Mice with implants recovered from laminectomy for at least one week before proceeding to the window placement.

#### Third procedure: Teflon AF and window placement

Under general anesthesia (2% isoflurane), we removed the Kwik-sil and PRECLUDE® Pericardial membrane that was placed during the laminectomy procedure. Then, we immediately covered the spinal cord with a saline-soaked sponge (Ethicon, Surgifoam®). We cut a Teflon AF film (VICI Metronics, Poulsbo, WA; Teflon AF 2400, 50 µm thick) to the size of the exposed spinal cord and placed it directly over the leptomeninges. We placed Kwik-Sil above the Teflon AF then adhered a 3.0-mm glass coverslip (#0, CS-3R-0, Warner Instruments) on top, forming a multi-layered tier above the spinal cord, and we allowed it to harden for 10 minutes. We then added a supportive coat of bone cement or Norland UV-curable optical adhesive (NOA 81) from the base of the implant to the coverslip. We administered Carprofen (5.0 mg/kg, sc) for postoperative analgesia.

#### Teflon AF sourcing

The Teflon AF 2400 (33 or 50-micron thickness) used in this study was obtained from Amos Gottlieb at Random Technologies LLC. VICI Metronics commercially offers Teflon AF film of different thicknesses (https://www.vicimetronics.com/products/teflon-af-films-1) including 50-microns (SKU: AF-050-025-025).

### Additional surgical procedures

#### Retro-orbital injections

We delivered AAV-PHP.eB-CAG-nls-GFP (Addgene, 104061-PHPeB) and AAV-PHP.S-CAG-tdTomato (Addgene, 59462-PHP.S) via retro-orbital injections. We briefly anesthetized animals (2% isoflurane) and placed them on a covered heating pad (Kent Scientific, RT-JR-15). We pulled back the skin surrounding the to-be injected eye to force the eye to partially protrude. For analgesia, we applied a drop of 0.5% Proparacaine HCl (Medline Industries, 24208-730-06) onto the to-be injected eye, then dabbed away excess fluid using a gauze placed at the medial canthus. We then slowly guided the needle at ∼30° angle, with the bevel facing medially, until the needle contacted the underlying bone. We confirmed the absence of blood and then injected 100 µL of solution. We delivered 100 µL of the following viral titers: 2.3 × 10^11^ vg/mouse for AAV-PHP.S-CAG-tdTomato (2.3 × 10^13^ GC/mL) and 2.2 × 10^11^ vg/mouse for AAV-PHP.eB-CAG-NLS-GFP (2.2 × 10^13^ GC/mL). After the injection, we applied eye ointment (Alcon, 02444062 [Systane]) and transferred the animals to a new heated cage.

#### Intraspinal injections

We drove expression of Cre pan-neuronally in the spinal cord via intraspinal injection of AAV2retro-hSyn-Cre (Addgene, 105553-AAVrg) (**Fig. 4a**). Briefly, we pulled a glass micropipette (WPI, 1B100F-4, 1-mm borosilicate glass capillaries) using a PUL-100 puller (WPI) and cut it to a tip diameter of 50-100 µm. We attached the glass micropipette to a 10-µL Hamilton syringe (7653-01) using compression fittings (Hamilton) and filled it with mineral oil. We then aspirated the virus into the micropipette using an automated pump (Harvard Apparatus, 70-4507 [Pump 11 Elite]) attached to a stereotactic arm. We injected 50 nL at 10 nL/min into the dorsal horn ∼500 µm ventral to the surface of the meninges and held the micropipette in place for 3 minutes after injection. We made two injections on the same side of the spinal cord.

#### Neuropathic pain model

We performed spared nerve injury (SNI) as previously described (Shields et al., 2003; Corder et al., 2019). Briefly, in a subset of CX3CR1-EYFP mice, we anesthetized animals (2% isoflurane for both induction and maintenance) and transferred them to a stereotaxic surgical station (Kopf Instruments, Model 942). To remove hair from the hindlimb, we used a shaver (Wahl Professional, 8685) rather than hair removal cream, due to the risk of chemical burn and any of the substance getting into the open wound, which would alter the pain model. We then cleaned the surgical site using 70% ethanol followed by Betadine, and again 70% ethanol. We made a small incision in the skin along the mediolateral axis, then widened the hole by inserting and spreading scissors (FST, #14072-10) rather than performing additional cuts. This exposed the *biceps femoris* muscle that, along with the *artery genus descendes*, we use as a landmark to begin incision to expose the sciatic nerve. To avoid the possibility of accidentally cutting the nerve, we part the overlying muscle using forceps (FST, #11231-20 and #11223-20). After exposing the sciatic nerve, we identified the common peroneal (CP), tibial (T), and sural (S) branches. We performed a CP and T ligation near the CP-T-S branch point using 8-0 silk sutures (S&T, #03192) then transected (FST, #91500-09) the CP and T branches at two locations about 1.0-mm distal to the ligation site. We closed the muscle using 6-0 silk sutures (Henry Schein, # 101-2636) and sealed the skin using cyanoacrylic glue (3M, Vetbond [084-1469SB]).

To account for the fact that some animals had different pre-injury CX3CR1 induced GFP fluorescence (*F*) between left and right dorsal horns, we performed SNI surgeries on either the left (n = 3) or right (n = 1) sciatic nerves, after we measured the spinal cord microglia bulk fluorescence bilaterally and divided into groups of animals that either had *F*_ipsi_>*F*_contra_ or *F*_ipsi_<*F*_contra_.

### Optical imaging through the spinal window

#### Stereoscope validation of window optical clarity

Under anesthesia (2% isoflurane), we periodically checked the clarity of the window using a stereo microscope (Leica MZ12.5 stereoscope). We acquired images of the window using a large sensor camera (The Imaging Source, DFK 33UX183c) and vendor-provided image acquisition software (The Imaging Source, IC Capture 2.5). We strove to maintain optical imaging parameters (zoom, white balance, and other properties) across real-color image acquisition sessions.

### *In vivo* spinal cord imaging

#### One-photon imaging setup

Throughout the study, in both anesthetized and awake imaging, we acquired *in vivo* imaging data using a 3i VIVO Multiphoton Movable Objective Microscope (MOM) with Phasor equipped, which permitted switching between one- and two-photon imaging in the same animal and microscope, without touching the animal or altering the setup (**Fig. 3c,g**). We suppressed vibrations with an optical vibration control air table (TMC, 14-416-45). The one-photon light path consists of an LED light source (Excelitas, X-Cite 110 LED) directed into a vertical illuminator (Olympus) containing a green (Semrock, GFP-3035D-OMF-ZERO) or red (Semrock, TxRed-4040C-OMF-ZERO) filter set. Emitted fluorescent light reflected back through the illuminator through a tube lens (Olympus, U-TLU) onto a CCD or sCMOS camera.

We used several cameras for widefield fluorescence imaging: Photometrics CoolSNAP EZ, Photometrics Kinetix, Hamamatsu Fusion BT, Hamamatsu Prime BSI, PCO pco.edge 4.2 bi USB, and Zeiss AxioCam 712. We acquired videos for *in vivo* spinal cord imaging of awake animals at 10- or 20-ms exposure time across all cameras. Each camera had the following specifications and settings during use. The read noise range is given as most cameras had high sensitivity and dynamic range modes. We used a Photometrics CoolSNAP EZ with a QE of 60-65% QE and RMS read noise of 6-8 e^-^. We used Photometrics Kinetix cameras with a QE of ∼95% and RMS read noise of 0.7–1.3 e^-^ using sensitivity or dynamic range modes. For the imaging sessions using the Kinetix, to accommodate large differences in brightness between animal lines and fluorescent proteins, we used dynamic range mode for **Fig. 3** and **Fig. 6** animals, sensitivity mode for **Fig. 5** animals. We used a Hamamatsu Fusion BT with a QE of ∼95% and RMS read noise of 1.0–1.4 e^-^. We used a Photometrics Prime BSI with a QE of ∼95% and 1.1–1.8 e^-^ RMS read noise. We used a PCO pco.edge 4.2 bi USB with a ∼95% QE and 1.1–1.8 e^-^ RMS read noise. We used a Zeiss AxioCam 712 with a QE of ∼72% and RMS read noise of 1.29–2.2 e^-^.

We collected most data used in this study from Photometrics CoolSNAP EZ CCD and Photometrics Kinetix sCMOS cameras. To simultaneously image the left and right spinal cord, we used low magnification objectives: 4x/0.16NA (Olympus, UPLXAPO4X), 5x/0.16NA (Zeiss, 420630-9900), and 5x/0.25NA (Zeiss, 440125-0000-000). We conducted the majority of imaging using the 5x/0.25NA objective. Additionally, a 2x/0.08NA (Olympus, 1-U2B921) objective provided an overview for a very large field of view (FOV) at the cost of reduced signal. We used high magnification objectives, 20x/1.0NA (Zeiss, 421452-9600) and 20x/0.45NA (Olympus, LCPLN20XIR) to conduct high resolution imaging of cellular or axon morphology (**Fig. 6b-d**). To compare intensity across time, we maintained the LED power at a constant level on a per animal basis and periodically checked the power output using a light meter positioned at the focal point of the objective (see <u>LED power output for one-photon</u> <u>imaging</u>).

#### Two-photon imaging setup

The two-photon light path of the MOM scope consisted of an imaging laser (Coherent, Chameleon Discovery Ti:Saph) and a phasor laser (Spectral-Physics, FemtoTrain laser) that are directed to a 3-galvo Vector RS+ module, allowing for either dual-galvo (Cambridge Technology, 6215H) or resonant scanning. The latter enabled 30 fps full-frame scanning. After the galvos, laser light is passed through to the sample and emitted light is reflected to the PMTs through a long-pass dichroic (Chroma, 670 LP Dichroic) above the objective. We filtered IR light out with an IR blocking filter (Semrock, FF01-750/SP-25), split light to the PMTs with a red/green PMT Dichroic (Chroma, 565 dcxr), and placed green (Chroma, ET525/70m-2pm) and red (ET605/70m-2p, Chroma or FF02-641/75-25, Semrock) filters in front of each PMT (Hamamatsu, H11706P-40 GaAsP). We controlled focus and the axial position of the objective using a micromanipulator system (Sutter Instrument, MPC-200 and ROE-200). We set the laser to 920 nm when collecting images from animals expressing GCaMP, EYFP (CX3CR1-EYFP), or GFP (Thy1-GFP). For animals expressing tdTomato, we either imaged at 920 or 1050 nm. We controlled laser power with a Pockels cell set to 40% (4V, for a subset of experiments) and normally to 50–80% (5–8V) of maximum power. We measured laser power, using a near IR compatible light sensor and power meter (Thorlabs, S121C and PM100D), delivered at the focal plane to be 64.27 (40%) and for the majority of experiments at 88.73–135.3 (50–80%) mW. We set the gain of red and green PMTs to 65–80%, and held this gain constant for each animal for longitudinal cross-session imaging experiments.

#### Optimizing one-photon imaging for *in vivo* spinal cord imaging

To collect usable data for downstream analysis, we needed to record videos with <10 ms exposure times that minimize the blurring that occurs during rapid and large shifts of the spinal cord (**Extended Data Fig. 7b**). For example, longer exposures would hinder accurate motion correction both within and across sessions, and limit quantitative measurement of fluorescence signals, especially when conducting Ca^2+^ imaging. And temporal downsampling of the movies to boost signal-to-noise (SNR) is not always viable as there can be significant motion (e.g. larger than the size of a cell) between individual frames (e.g. at normal 13-20 Hz acquisition with 10-ms exposure). However, this fast exposure time leads to issues with older CCD cameras that have high read noise, as that can diminish the signal-to-noise ratio (SNR) and the ability to detect biologically relevant signals, such as transients during Ca^2+^ imaging. Thus, we optimized several aspects of the preparation to maximize SNR and minimize blur. Modern sCMOS cameras greatly aid in achieving this aim. We tested several systems (see <u>One-photon imaging setup</u>) and found that those with quantum efficiency >90-95% in the 500-600 nm wavelength range and read noise <1.4 e^-^ allowed for greatly improved imaging quality compared to low quantum efficiency CCDs or sCMOSes. For our initial studies, we collected data using a Photometrics CoolSNAP EZ CCD, then switched to the improved Photometrics Kinetix sCMOS. It is possible to use the effective global shutter in newer sCMOS cameras, such as the Kinetix, by only triggering the light source once all rows of the camera are acquiring data. This would further minimize the deformations seen due to fast rostrocaudal motion as the entire frame would be captured at once. This is in contrast to the majority of conventional fast-scanning (either with a galvo or resonant scanner) two-photon imaging systems, in which fast motion can often lead to deformations or missing data.

For bilateral spinal cord imaging of the entire or a large section of the FOV accessible post-laminectomy (**Fig. 4d**), we optimized imaging conditions by combining low-magnification objectives (4X/0.16NA, 5x/0.16NA, or 5x/0.25NA) with large-sensor sCMOS cameras (e.g. Photometrics Kinetix). Further, we wanted to minimize the effects of axial (dorsal-ventral) motion, which would cause cells to drop out or lead to significant changes in fluorescence intensity not related to Ca^2+^ activity. Another advantage of one-photon imaging with low-magnification objectives is the larger depth-of-field, which reduces sensitivity to dorsal-ventral (axial) motion. This feature reduces the need to compensate for axial motion with fast volumetric acquisition. Because the low-magnification objectives often have a lower NA, we tested a series of objectives and found that the Zeiss Fluar 5x/0.25NA objective, both theoretically and when measured, obtained the highest signal while having a tolerable field flatness that still allowed bilateral imaging of dorsal horn lamina I neurons and glia.

#### LED power output for one-photon imaging

To ensure the reliability of our fluorescence imaging measurements, we periodically tested the LED power output using a visible light sensor (Thorlabs, S120C) and power meter (Thorlabs, PM100D). For consistent power measurements across time, we centered the illumination profile on the center of the sensor and by directly focusing on the surface using the microscope’s software (SlideBook 6), we ensured that the sensor surface was in the focal plane of the microscope. We then used custom MATLAB scripts to create a calibration curve between the software’s arbitrary power scale and power (mW) or irradiance (mW/cm^2^). For all one-photon imaging experiments we acquired data with LED power set to 30-40 in SlideBook, which corresponded to total power (at given SlideBook power) of blue excitation light at the focal plane of 3.52 (30) and 4.28 (40) mW with the Zeiss Fluar 5x/0.25NA, 2.3 (30) and 2.78 (40) mW with the Zeiss 5x/0.16NA, 5.47 (30) and 6.61 (40) mW with the Zeiss 20X/1.0NA, and 3.71 (30) and 4.49 (40) mW with the Olympus 20X/0.45NA.

#### Microsphere measurements with and without Teflon AF

To confirm that Teflon AF did not affect optical imaging quality, we conducted microsphere measurements. We serially diluted and sonicated 1-µm yellow-green fluorescent microspheres (Invitrogen, F8765) to a final concentration of 1:10^6^ from stock in dH_2_O then placed the resulting solution on a standard glass slide and placed a #0 coverslip on top. We stored beads in the dark at 4 °C. We conducted one-photon imaging with a water-immersed Zeiss 20X/1.0NA at 5.47 mW (30 power in SlideBook) at 100 ms exposure followed by two-photon imaging at 920 nm with 7.43x zoom and power delivered ∼64 mW (PMT gain 90%) or ∼135 mW (PMT gain 65%). We placed the Teflon AF on top of the coverslip and repeated the imaging procedure on the same regions. To correct for any residual bidirectional scanning pixel shift errors, we used the *Correct X Shift* plugin within ImageJ. To plot the profile through individual beads, we selected individual beads, then used the *Plot Profile* built-in ImageJ tool followed by normalizing the profile intensity *I* at each location *x* by 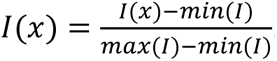 Results in **Extended Data Fig. 2k-m.**

#### Setup for spinal cord recording

To reduce the strain of the animal’s lateral movements on the implant, we placed the mice in an imaging apparatus with blinders (**Fig. 4c**, **Fig. 5d**, **Extended Data Fig. 9a**). These blinders underwent several iterations, starting with solid 3D printed materials (MakerBot, Nylon 12 Carbon Fiber [375-0061A] or ABS-R [375-0071A]). We settled on an infrared acrylic (ePlastics, ACRY31430.125PM11.555x11.850) that is optically transparent λ>700 nm, visibly black, faster to manufacture, and less likely to break than 3D printed materials. The blinder reduced the animals’ ability to see both the surroundings and the experimenters, while allowing us to use infrared 850 nm LED lights (Waveform Lighting, 7031.85; LIYUDL, B071KPSGCT; and Shenzhen Jing Cheng Digital Surveillance Co, IRINB04L) and cameras to monitor animal behavior. We prevented the mice from twisting and reaching for the objective by placing a top blinder that we could slide in and out as needed. We fixed mice in the apparatus using either one or two standard clamps (Thorlabs, PC2), either on the side bars or on the rostral or caudal wings of the stabilizing plate. By the final iteration we found that we only needed to use a single clamp— this simplified the design of the apparatus and allowed more flexibility, and made it possible to mount animals more rapidly. We used a goniometer (Thorlabs, GNL20) that gave us freedom to adjust the tip and tilt of the spinal window for each animal with the focal plane of the microscope. We first aligned the planes by eye, then conducted further adjustments while imaging until we determined we had in focus the maximum amount of the field of view. It should be noted that due to the large rostro-caudal and medio-lateral size of our window, the spinal cord in both orientations has a curvature that can at times prevent the entire FOV being in focus, requiring acquisition of 3D stack videos. We raised or lowered the animal with a jack (Thorlabs, L490) or mounting bracket (Thorlabs, C1515).

We used the same infrared acrylic to custom cut a 21.8-cm diameter circular gridded floor (see design files in GitHub repository). This design provided a visibly opaque surface for the animal to run on during imaging sessions, while we monitored limb movement with an infrared-capable camera and delivered peripheral stimuli (**Fig. 5d**, **Supplementary Video 14**). A prior version of the running wheel used an aluminum (to avoid rusting) grid (McMaster-Carr, 92725T51) with the same spacing and hexagonal pattern cut to fit into the circular running wheel. With either design, we attached them onto custom 3D printed parts that connected to the rotary encoder (Signswise, LN11-ERGA) which measured animal locomotion. We held the rotary encoder in place using a custom-designed 3D printed part that attaches onto standard ½’’ Thorlabs optical posts for integration into the setup.

We monitored animal behavior using 2-4 cameras in three configurations: 1) cameras monitoring the left and right side of the body, 2) cameras monitoring the face or rear of the animal, 3) cameras monitoring the left, right, and bottom of the animal, and 4) a camera zoomed in on the face. We synchronized these cameras with the microscope via TTLs delivered to each camera, via BNC and Hirose (The Imaging Source, CA-x2-HIR-OE/1.5) cables. Using a BNC splitter, we sent the same TTLs to a logic analyzer (Saleae, Logic 8) for synchronization with sensory stimuli, sound, and other experimental devices. To protect the bottom camera from animal waste, we laser cut a slot into a plastic cover (Corning, 07-200-600) that allowed it to slide into the space between the rotary encoder and 3D printed running wheel connector.

#### Awake imaging of the spinal cord

Before initiating awake recordings, we habituated mice to the imaging room and/or apparatus for 1-3 days. The room is 53.0–53.3 dBA with all instruments off and 59.7–60.4 dBA with all instruments on; measured with a sound meter (Tadeto, SL720) at the location where the animal would be during imaging. For experiments in which no stimulation is given, we imaged with all lights off, at an effective 0–1 lux. To facilitate visualization of hindpaw and accurate experimenter delivery of stimuli along with minimizing change in illumination for the mouse, we provided ∼4 lux red light using LEDs (ALITOVE, AL5RWPBK12V) controlled by a flicker-free LED dimmer (Waveform Lighting, 3081). At the beginning of each imaging session, we loaded animals onto the apparatus (while we lit the room with ∼280 lux white light) using the clamps and allowed them to habituate for several minutes with the lights off. We then collected the following one-photon widefield images or videos: 1) reference images in green and red channels at 1 × 1 and 2 × 2 binning; 2) a 3D stack in the green and, where applicable, red channel with 10 µm spacing over a range that started at the glass coverslip surface and extended into the spinal cord until we lost focus; 3) ∼13.9 or 20 Hz videos in the green, and when applicable red, channel at 1 × 1 and 2 × 2 binning (normally for 1,000 and 300 frames, respectively). For animals that underwent two-photon imaging, such as Thy1-GFP and CX3CR1-GFP animals, we collected: 1) maximum FOV (1x zoom) time series with 1 frame averaging, 2) a 3D stack (for CX3CR1-GFP animals) with 1 µm spacing, and 3) 2.0736x and 3.58318x zoomed in time series videos.

For GCaMP imaging, we used a modified protocol for the final widefield imaging step. We collected a ∼2-min baseline period in which we did not intentionally present any peripheral stimuli. We then delivered 2 to 5 blocks of stimuli: noxious cold (∼4 °C water), noxious mechanical (pinch with forceps), noxious heat (∼55 °C water), air puff (gas duster), or loud sound (93.4-94.1 dBA noise). For certain animals, we also delivered a noxious mechanical (pin prick, 25G needle), innocuous mechanical (2g von Frey hair), or brush (NicPro, MG015, Flat 1 and Round 1) to the hindpaw.

#### Sensory stimuli delivery during awake recordings

We delivered sensory stimuli to the left and right hindpaws or the face. As glass syringes have a better heat capacity than plastic, we used 2-mL glass syringes (Synthware Glass Syringes, S371202) to deliver noxious cold and heat stimuli. We pre-heated or -cooled the syringes in water baths at the set temperature, which reduces the drop in temperature of the applied liquid during the time it takes to transfer to the animal’s hindpaw. We cooled water to 4 °C (Yeosen, PH-F3) and heated water to ∼55 °C using a hot plate (Fisher Scientific, HP88854200) set to 170 °C. We concurrently monitored temperature. This approach made it possible, inexpensively and easily, to obtain consistent water temperature, compared to custom Peltier setups or water baths. For certain animals (**Fig. 4e-g**) we measured the force applied during the pinch, using a force sensitive resistor (Adafruit, 166) attached midway along the forceps (AVEN, 18434). For these studies we did not convert the force measurements from arbitrary to real units, as this approach was used solely to automatically synchronize stimuli with fluorescence imaging data and the behavior cameras’ videos. As we periodically observed bursts of SCPN activity without applying a stimulus, and to address the possibility that it was locomotion or other behaviors that drove SCPN activity, even during application of a noxious stimuli, we also delivered a loud sound directed at the animal’s face using a piezo buzzer (Intervox, BRP3018L-12-C). At the animal, the sound would be ∼93.4-94.1 dB with an intensity in the 2000 ± 500 Hz range. For certain animals, we used a small infrared LED placed in the FOV of one of the behavior cameras to indicate when we had activated the sound stimulus. This allowed further confirmation when synchronizing with video files. We sent a TTL to the logic analyzer, on a unique channel, whenever we triggered the sound. We used a 25G needle to deliver a noxious pin prick from below the animal through the floor grating. We considered this a “light” pin prick, as we tried to slowly bring the pin to the animal’s hindpaw and withdrew the pin on first contact, in contrast to a “heavy” pin prick in which a stabbing motion is made. The latter is more likely to puncture the animal’s skin and cause bleeding, which can alter interpretation of any subsequently delivered stimuli.

Using a custom-made clicker, we manually annotated each time we delivered a stimulus. The clicker synchronized this timing by sending a TTL into the same logic analyzer (Saleae, Logic 8) data stream as the microscope TTLs. To align the exact frame when each stimulus was delivered, we conducted post-hoc analysis by going through behavior videos for a given imaging session, frame-by-frame, near times when stimuli were indicated to have been delivered. We then annotated when the forceps, needle, von Frey fiber, or liquid first touched the animal’s hindpaw. We used these frame times in downstream analysis.

#### Recording during virally-mediated gene expression

Compared to expression of GFP, we observed dimmer tdTomato expression, which took longer to be detected, likely owing to the increased time to synthesize and transport the needed material from the sensory neurons’ somas and the more diffuse localization of the tdTomato compared to nuclearly localized GFP. Near daily imaging afforded by our preparation identified the expected logistic-like curve in expression of both proteins (**Fig. 3m**), which had a time course to maximal fluorescence that was similar to prior fiber photometry measurements in the brain (Fenno et al., 2020).

#### Anesthetized animal recordings

We anesthetized animals (2% isoflurane) inside an induction chamber and transferred them to the same setup as used for awake imaging, but with modifications for anesthetized recording. We provided heat (Stoelting, 50300) while we delivered 2% isoflurane via a nose cone (Kent Scientific, SOMNO-0801) and removed excess isoflurane using a vacuum line connected to the nose cone, or with a 3D printed scavenger placed around the nose cone. We monitored breathing throughout imaging. To prevent liquid spilling onto the heating pad and other devices, we placed a thermoplastic (HDPE) sheet below the animal’s hindpaws. We applied noxious thermal stimuli (∼4 °C for cold and ∼55 °C for heat) using a liquid drop, then rapidly removed it with a syringe attached to a vacuum line, which avoided tissue damage. We delivered a noxious mechanical pinch to the hindpaw or tail using forceps (AVEN, 18434) for ∼1–2 s. We delivered air puff for ∼1–5 s using a compressed gas duster (Dust-off 8541677532 and 8541677551); we only delivered with the canister facing in the upright orientation to avoid delivery of liquified gas, which can cause tissue damage (frostbite). For a subset of experiments, we delivered sound toward the face, as previously described. After experiments we returned the mice to the home cage and monitored them until they awoke.

#### Recording in anesthetized and awake states in the same session

For the session shown in **Fig. 3p** and **Extended Data Fig. 8c-e**, we mounted the animal as previously described. We then recorded the animal for ∼4 minutes before manually bringing a nose cone to the animal’s face while delivering 2% isoflurane at a higher than normal flow rate (∼4.0 L/min O_2_) until the animal became unresponsive and stopped moving. Then we reduced the flow rate to a normal maintenance rate (0.8 L/min O_2_). We continued imaging and finally removed the nose cone and allowed the animal to gradually wake up. For the experiments in **Fig. 3p**, there is a 05:36.9 min gap between the end of isoflurane and the return to the awake state.

#### Multi-plane recording of monocytes in CX3CR1-EYFP mice

We recorded multiple planes in CX3CR1-EYFP mice under general anesthesia (2% isoflurane) using the imaging setup previously described. To maintain optical clarity when using the 20x/1.0NA water objective during long-term recordings, we used ultrasound gel (Parker Labs, 638632490755), which has a similar refractive index as water, but is resistant to evaporation. We collected 9 or 11 planes from the spinal cord meninges to parenchyma consisting of slices spanning 212 and 159 µm, respectively. We manually set the spacing for the slices (**Fig. 6d**) based on planes with maximal difference in features (e.g. types of cells). We recorded at 920 nm for ∼21 and ∼15 mins delivering 61.05 ± 3.26 mW, Pockels cell set to 35% (3.5 V), and green PMT gain set to 67%.

#### MicroCT imaging and data processing

We conducted microCT imaging of naive and spinal chamber implant animals using standard imaging protocols approved by UCSF’s IACUC at UCSF’s China Basin. We induced and maintained animals under 2% isoflurane while scanning animals using the microCT component of the MILabs U-SPECT VECTor4/CT (MILabs B.V., Utrecht, The Netherlands) preclinical imaging system (MILabs Rec 12.00-st). To set the scanning bounds, we used built-in optical cameras followed by CT acquisition using X-ray tube parameters of 50/55 kV and 0.24/0.19 mA, 75 ms exposure per step, and 360° (0.375° step, 960 projections) scan acquired in step-and-shoot mode. We did not apply any binning during data acquisition (i.e. 1 × 1 binning). We created an isotropic reconstruction of microCT data using vendor-provided MILabs reconstruction software v12.00 at a voxel size of 0.02 mm × 0.02 mm × 0.02 mm with cone-beam filtered back-projection using the Feldkamp algorithm (Feldkamp et al., 1984).

We used a custom pipeline to process microCT data (**Extended Data Fig. 4g**), by first downsampling the raw files 2x in space, which facilitated faster processing. For data from whole body scans of animals, we manually went through each slice and removed scan artifacts where possible. For display purposes in **Fig. 1g** only—to allow easier reader visualization of bone, soft tissue, and spinal chamber implant as the quantitative difference in intensity is not critical for interpreting that data—we applied successive gamma correction (*V*_*out*_ = *V*_*in*_^*γ*^) with *γ* = 0.78 and *γ* = 0.70. By visually setting thresholds in ImageJ (version 1.53d) or Invaleus (version 3.1), we segmented bone, soft tissue, and (where applicable) 3D printed components and glass. In ImageJ, we then used the Volume Viewer method to construct a 3D mesh; with Invaleus we used built-in methods for setting masks and creating surfaces. We exported the resulting meshes and imported them into MeshLabs (version 2020.07). Within MeshLabs we manually removed vertices, faces, and edges that are due to scan and other artifacts. We then cleaned up the model, e.g. using Laplacian Smoothing (3 iterations), and ran Quadric Edge Collapse Decimation to reduce the number of vertices and faces to reduce computational load during 3D rendering.

For the model used to display the spinal cord implant chamber in **Fig. 1a**, we performed a virtual laminectomy by manually removing the overlying spinous process and lamina. Then to create a smooth surface mesh, we eliminated non-manifold edges and closed holes created by face removal. We exported the final mesh from MeshLabs as an STL file. We then imported files into FreeCAD for conversion from a mesh to a solid by converting the mesh to a shape and then converted the shape to a solid before exporting as a STEP file. To visualize 3D renders, we imported the resulting STL (MeshLabs) or STEP (FreeCAD) files into PTC Creo 6.x or 9.x. We converted spatial units to mm and scaled model 1000:1 to match the size and coordinate system commonly used in the rodent by checking that bregma to lambda distance was ∼4.2-mm, as expected. To display high-quality photorealistic renders, we assigned components realistic textures (e.g. metal, glass, etc.) along with using bump maps to simulate rough texture. These bump maps are not used when 3D printing or otherwise manufacturing the components and are purely for visual aid to the readers. We then used the Renderer module within PTC Creo with 64 direct bounces, 64 indirect bounces, quality level 5 shadows, and enabled global illumination, caustics, and interior lighting. For surgery videos, we used the Animation module within PTC Creo and a series of views and snapshots of the spinal implant components in various configurations. To minimize noise on each frame, we ran 2 min of rendering time per frame (∼4000–4500 samples) on a NVIDIA A5000 GPU. We saved the animation as an uncompressed AVI, which was then converted to an mp4 with ffmpeg before use in the surgery videos (**Supplementary Video 2-4**).

### Generation and sources for 3D models

We created 3D models using a mix of our own microCT generated data, our own models (e.g., for side bars and stabilizing plates), and publicly available designs. We downloaded the following models and used them in either **Fig. 1a** or Fig. 4c. We downloaded Thorlabs parts from their respective product web pages as STEP files. The 4x objective in Fig. 4c is https://www.thorlabs.com/thorproduct.cfm?partnumber=RMS4X-PF. McMaster-Carr provides design files for many of their components; we downloaded and used STEP files for screws (McMaster, 91781A350) and Neodymium magnets (McMaster, 5862K141). We downloaded the following GrabCad SOLIDWORKS files: https://grabcad.com/library/incremental-optical-rotary-encoder-400-pulse-1. For camera models, we used STEP files provided by The Imaging Source: https://www.theimagingsource.com/en-us/product/industrial/37u/dfk37bux252/. We converted all STEP and other formatted files into CREO PRT files to integrate into our assemblies.

### Histology

We perfused mice with phosphate-buffered saline (PBS), followed by 4% formaldehyde (4% FA, 10% formalin [Thermofisher Scientific, #119690010] in 1x PBS), dissected out the spinal column, and left the spinal column overnight in 4% FA. We manually dissected the spinal cord after the removal of overlying vertebrae and then switched samples to either further fix in 4% FA or PBS followed by cryoprotection with 30% (w/v) sucrose. We cut sections on a freezing microtome (Microm, HM 440E) at 100-μm thickness. We then washed samples three times with PBS in 0.3% or 0.8% Triton-X100 and with NGS blocking. We identified astrocytes by IHC with GFAP (#Z0334, Agilent Dako). We imaged microglia using the endogenous EYFP signal. We acquired Phox2a images by endogenous GFP or tdTomato. We stained cells or their nuclei using Neurotrace or DAPI, respectively. We acquired all histological images using an Olympus FV3000 equipped with 405, 488, 561, and 640 nm OBIS Coherent lasers.

### Behavioral testing

#### Assaying locomotion in the open field

The open field arena consisted of a custom-built 2-ft diameter white base (HDPE, TAP Plastics) and 15.5–16’’ tall white walls (Mr. Plastics, WHT POLYSTYRENE). We provided overhead lighting using either two LED lamps (Barrina, INWT504005650Fc and JOOFO, clipper0722-54) or LED light strips (Waveform Lighting, 3004.40 [5000K]) controlled by flicker-free LED dimmer (3081, Waveform Lighting). We acquired videos of animal locomotion using cameras (The Imaging Source, DMK 21AU04 and DFK 42BUC03) and image acquisition software (The Imaging Source, IC Capture 2.4 or 2.5 or Mathworks MATLAB Image Acquisition Toolbox). Before each experiment, we measured light power (JRLGD, LX1010B) and adjusted lighting to deliver ∼100 lux of light. We cleaned the arena with 70% ethanol before and after each animal’s session. We placed animals in the center of the open field arena and allowed them to freely locomote for the duration of recording (30 min, 15 Hz). We ran up to three animals simultaneously in three arenas in the same room.

#### Motor control using rotarod

We tested locomotor control using an accelerating rotarod assay. We used two different rotarod devices (Ugo Basile, 7650 and 47650) that consisted of a 30-mm diameter rod elevated above a floor that automatically detected when an animal fell. We habituated animals in the behavior room for 30 mins to 1 hr before testing. As the rotarod assay is a test of motor learning as well as motor coordination, we also evaluated and recorded animals for 3 trials during a single behavior session (one session per day). For each trial, we placed animals on the rod and positioned them facing in the direction opposing the rod’s motion to minimize the chance that they would immediately fall off when facing the direction of motion. The rod started at 4 rotations/min and increased at a consistent rate to 30 rotations/min for 5 mins, the final speed was maintained until the animal fell off. Animals had at least 3 mins between trials within a single day’s session; if we ran more than five animals during a single session, the time between sessions was at minimum the time taken to run all other cohorts of mice. Animals can “flatten” themselves onto the rotarod and thus hang onto the rod after they would have been evaluated as failing the task. This can lead to incorrect automatic measurement of the latency to fail the task and introduces different metrics between animals that fall and those that can wrap their bodies around the rotarod, which can also differ by age and body size. To circumvent this, we used cameras (The Imaging Source, DMK 21AU04 and DFK 42BUC03) to record all rotarod sessions and post-hoc determined the time when the animals either fell or hung into the rod and began their first circle around the rotarod.

#### Mechanical sensory thresholds

We measured mechanical sensory thresholds using von Frey filaments (Stoelting, 58011) and the SUDO up-down method (Bonin et al., 2014). We habituated mice for 30 mins - 1 hr on a custom von Frey rack (Ahanonu and Corder, 2022). We started with the 0.6 gram-force (gf) hair and then tested subsequent steps at lower or higher gf hair if the animal did or did not exhibit responses, respectively. We used the following series of von Frey hairs, numbered 1-9 for SUDO threshold calculations: 0.04, 0.07, 0.16, 0.4, 0.6, 1, 1.4, 2, 4 gf. We define responses as any of the following that occurred while the von Frey hair contacted the hindpaw: hindpaw lifting (withdrawal), shaking, licking, or guarding. We did not count toe spreading—splaying or other movements of the toes in response to contact with the von Frey hair but no withdrawal behaviors—as a response nor did we count responses occurring right after the offset of the von Frey hair, so as to avoid behaviors due to flicking of the hairs during offset or related artifacts. We pressed with von Frey hair until it bent to ensure consistent force application across trials and animals.

We calculated the SUDO PWT 50% mechanical threshold using *PWT* = 10^*x*∗*F*+*B*^, where x = 0.24, B = -1.54, and F is the final filament number, along with a +0.5 or -0.5 adjustment factor depending on whether the animal did not or did respond to the final filament, respectively. The estimated 50% threshold using the Chaplan Up-Down method is calculated as *T*_50%_ = 10^*log*_10_(*F*)+*k*∗*D*^, where F is the force (gf) of the final filament hair, *k* is the lookup value based on the sequence of responses, and *D* is the mean gf difference between adjacent hairs in the sequence. We used the lookup table as in the original Up-Down method (Chaplan et al., 1994). As we did not observe a difference between PWT and Chaplan Up-Down calculations, we present Chaplan Up-Down values. The von Frey hair estimated gf from the manufacturer can vary; thus, for each von Frey hair used we confirmed the gf using a balance (Ohaus, Adventurer SL).

### Data analysis

#### Open field analysis

To analyze open field behavior, we trained a DeepLabCut (version 2.2.3) model using 20 manually annotated frames from three mice with spinal chambers implanted. We selected frames based on visual appearance by using k-means clustering. We manually annotated the nose, torso (center, left, and right), and tail (base, mid, and tip). Training parameters: 600,000 iterations (**Extended Data Fig. 4j**); net type, resnet101; dataset augmentation, imgaug; global scale, 0.8; batch size, 1; fully connected parts; and GPU, NVIDIA A5000 (24 GB ram). Model training and test errors, with likelihood cutoff of 0.1, are 1.37 and 4.73 pixels. To confirm the accuracy of the model, we visually inspected by rapidly scrolling through dozens of open field movies, collected from the same arenas but using mice with and without spinal chamber implants. We imported body part locations from DeepLabCut CSV files into MATLAB using a custom CIAtah function. For a subset of videos we used an alternative ImageJ- and MATLAB-based algorithm for tracking mice that we previously developed and validated (Li et al., 2017).

To evaluate locomotor speed, we calculated the speed between (x,y) point pairs between successive frames using the following formula 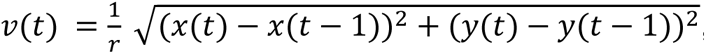, where t = frame, r = mean interframe interval time (s), and x,y are the animal’s position coordinates. We set the first frame’s speed to zero, to ease downstream computational analysis. We converted speeds to real units by calculating a pixel per cm conversion factor using a custom CIAtah GUI in which we selected a 60.96 cm distance for each video that corresponded to the diameter of the circular arena. To produce Fig. 1n, we took the mean speed over all the frames in a session.

#### Behavior video analysis

To track body parts in multi-camera behavior videos, as in Fig. 5d,**g**, we trained a separate DeepLabCut (version 2.2.3) model for each camera, rather than a generalized model across all cameras. We used 20 manually annotated frames from each movie for each model. We selected frames based on visual appearance by using k-means clustering to select frames from distinct clusters. Model training errors, with likelihood cutoff of 0.1, are 1.04, 1.19, 1.1, and 1.14 px for bottom, face, left, and right cameras, respectively. Model test errors, with likelihood cutoff of 0.1, are 4.06, 7.66, 7.36, and 5.78 px for bottom, face, left, and right cameras, respectively. Training parameters: 500,000 iterations (**Extended Data Fig. 9b**); net type, resnet101; dataset augmentation, *imgaug*; global scale, 0.8; batch size, 1; fully connected parts; and GPU, NVIDIA A5000 (24 GB ram). We visually inspected the tracking afterwards to confirm the general accuracy of the model, including when noxious stimuli were applied.

We chose to track several body parts based on their high probability of being in the camera FOV in almost all frames in an imaging session and likelihood of exhibiting significant motion during nocifensive behavior. We manually annotated the nose, eyes, tip and base of the ears, forepaws, hindpaws, and base of the tail. We separately annotated left and right hind and forepaws across the left and right facing cameras, respectively. In the bottom facing camera we annotated all of them. For the purposes of downstream analysis, owing to the rigid nature of the face, we did not distinguish between left or right sides of the eyes, ears, or mouth and combined the analysis of multiple views of each regardless of the side of the body. We combined the tracking of the ear tip and base to simplify analysis. For some videos, such as in Fig. 4g, we annotated a fixed bar to create a stationary control object to demonstrate stability of recording and tracking along with specificity of animal movement. It is possible to calibrate multiple cameras to enable 3D tracking of animal body parts. This enables more detailed analysis of body part movements, but increases the complexity of the setup and requires additional calibration steps.

We used the same method to calculate the speed as described in the Open field analysis section. To reduce the chance that we are using frames with sub-optimal tracking, we only used frames with likelihood >0.1. We used objects (e.g. objectives and blinders) with known size in the cameras’ FOVs to calculate the pixel to cm conversion factor, which we used to convert the *x* and *y* tracking from camera pixel to real units (cm). In certain cases, such as in Fig. 5g, to reduce the influence of small discrepancies in the frame-to-frame tracking, we smoothed the estimated body part speeds using a moving mean with a window of 5 s (100 frames).

#### Calculating locomotor speed using rotary encoders

The rotary encoder (Signswise, LN11-ERGA) outputs two channels, A and B, each time it is moved in a clockwise (*CW*) or counterclockwise (*CCW*) direction. We calculated both directions of motion independently using the equations *CW* = (*A* > *B*) > (*A*(*t* + 1) − *A*(*t*)) and *CCW* = −1 ∗ [(*B* > *A*) > (*B*(*t* + 1) − *B*(*t*))] where *A* and *B* are the vectors over all time points. We then combined them to obtain the total speed (i.e. events output by the encoder) with *v*(*t*) = |*CW*(*t*) + *CCW*(*t*)| over all frames. To calculate a conversion factor of 0.0691144 cm per pulse, we determined the circumference at the position on the running wheel where the animal would be and divided it by the 600 pulses per 360° rotation of the rotary encoder or as a formula 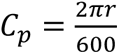 where r = ∼6.6 cm (radius from the center of running wheel to animal position). For display purposes, on some locomotor traces we downsampled the vector in time by dividing the speed vector *v* into evenly spaced groups of 4 frames and taking the mean within each group.

#### Ca^2+^ imaging data preprocessing

We processed Ca^2+^ imaging data using CIAtah, our Ca^2+^ imaging analysis software (Corder et al., 2019), and custom scripts in MATLAB (2022b). In general, we performed the following steps: spatially downsampling (for a subset of movies), detrending, calculating motion transformation coordinates, spatial bandpass filtering, registering each frame to reference frame, adding borders, fixing any problematic frames by setting their values to NaN, calculating relative fluorescence change, and temporally downsampling (for a subset of movies). We detail each step below.

To increase signal-to-noise and improve processing speed, we spatially downsampled each frame in the *x* and *y* lateral dimensions by conducting 2 × 2 or 4 × 4 bilinear interpolation. To account for photobleaching during imaging, we detrended the Ca^2+^ movies by calculating the mean for each frame, temporally ordered them from the first to last frame, and then fit a 1st- or 3rd-order polynomial curve to the fluorescent values. We then subtracted all pixels values in a given frame from the fitted values at each time point and added the mean, which detrended the movie while keeping the intensity values in a similar range as the raw movie and prevented the introduction of negative values that would complicate downstream analysis.

For movies that used rigid motion correction, we registered all frames to a reference frame using *TurboReg* (Thevenaz et al., 1998)). Here, we selected a subsection of the field of view with high-contrast features, such as dorsal veins and ascending venules, and minimal artifacts (such as bubbles or dust) that would affect registration. To improve motion correction, for each frame we subtracted the mean and then normalized the frame by subtracting it by a circular averaging filter (pillbox) of radius, one tenth the minimum row or column length (whichever was smaller). Subsequently, we cross-correlated the entire frame with a circular average filter of disk radius 3 pixels. We then performed an image complementation by subtracting each pixel from the maximum value in the frame; this inverted the image making blood vessels used for registration more prominent. We obtained 2-D spatial translation coordinated for each frame from *TurboReg* by comparing it to a reference frame kept constant for all frames in the movie. To improve cell extraction, we divided each frame of the raw movie by a bandpass-filtered version of that frame (cutoff frequency of 0–10 cycles) that suppressed background fluctuations, such as occurs with neuropil.To avoid issues with filtering frames containing NaN values, we performed this step before registration. We then registered each frame using the 2-D translation coordinates obtained for each frame. Due to differing amounts of motion correction across frames, which causes variable borders across frames, we added a fixed border for each frame in the movie, by calculating the maximum motion and using that or 14 pixels, whichever is smaller. We calculated relative fluorescence using the following formula: 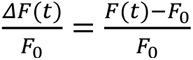 where F_0_ is either the mean image of the entire movie or the soft minimum calculated as the value equivalent to the bottom 0.1% of pixels. Finally, for a subset of movies we temporally smoothed the movie by downsampling the temporal dimension four-fold. For a *x* × *y* × *t* movie, this entailed bilinearly downsampling in *x* × *t* to minimize memory usage and improve processing times. This is equivalent to performing a 1D linear interpolation in time of the fluorescence intensity values of each pixel value. For a subset of movies that included areas outside the spinal cord in the imaging FOV, such as cement, we manually selected areas to keep and set all other areas to zero, which eliminated them from consideration during cell extraction, often improving the quality of extracted cells and reducing the number that need to be removed post-hoc.

For movies that used LD-MCM or NR-MCM, we performed similar pre-processing steps. Any deviations from the standard preprocessing are noted in their respective sections. To compare to LD-MCM and NR-MCM, we processed a subset of movies with NoRMCorre using our modified version of the repository (https://github.com/bahanonu/NoRMCorre) that only refactors the code into a namespace-safe package for integration into out codebase. We processed NoRMCorre motion corrected movies identically as described above and used default NoRMCorre parameters as defined in the CIAtah function *ciapkg.motion_correction.getNoRMCorreParams* (from commit a2e72a8) with the following modifications: d1 and d2 are the input movie tensor dimensions and grid_size of [64 64 1] for a subset of movies.

#### LD-MCM: deep learning-based identification of movie features

The basis for LD-MCM is identification of movie features followed by control point motion correction. The use of deep learning-identified control points addresses a challenging situation for prior motion correction methods, both for applications to spinal cord imaging and to other situations in which optical windows are used to gain access to the body. This includes contamination of the field of view by objects—such as vasculature due to angiogenesis (Extended Data Fig. 8a), bubbles in silicone adhesives (e.g. Kwik-Sil), cement or dust particles, and others—that directly overlay the FOV and move differently from the primary tissue of interest. For example, neovascularization or light fibrosis that overlaps with the FOV can impede motion correction as it can lead to motion correction algorithms that use those as fiducials instead of the spinal cord. Although this can be reduced by focusing down slightly more than desired to cause these elements to be further out-of-focus, this is not ideal and not possible in certain cases while still collecting usable data. LD-MCM helps mitigate the need to do this and provides superior performance. To consistently identify the same features, we took advantage of advances in deep learning that allow training of models with few examples to classify features. We use DeepLabCut (Mathis et al., 2020) in this paper but LD-MCM is agnostic to the method employed, just that it has the requisite accuracy to consistently identify features and generalize from the training set. While these deep learning algorithms are often used for tracking animal body parts or items in the environment, most of them do not contain priors that would preclude their use for tracking vasculature or other features. These algorithms are thus agnostic to the feature being identified as long as it has a consistent spatial structure across frames in a movie. Animals with prominent dorsal veins and dorsal ascending venules (dAV) made possible the most robust motion correction owing to the existence of distinct, stable landmarks.

The procedure for LD-MCM consists of the following procedures in MATLAB (using a mix of CIAtah and MATLAB built-in functions) and Python (for DeepLabCut analysis). To maintain consistency of pixel values and improve accuracy of feature tracking, we used custom MATLAB scripts to convert raw Ca^2+^ or Thy1-GFP movies from HDF5 (int16) to AVI (int8) format compatible with DeepLabCut. To avoid giving too much weight to outlier pixels, we calculated the soft maximum and minimum for each movie as the value equivalent to the 99.99th and 1th percentile of all pixel intensity values. We then converted using the following formulas: 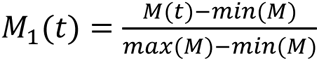 and *M*_*f*_(*t*) = (*M*_1_(*t*) ∗ 2 − 0.01) ∗ 255. This created a normalized movie then converted it to int8 units after shifting all values upward to avoid dark values that reduced feature identification accuracy. As we collected imaging data for some animals across multiple imaging cameras, which led to transformed FOVs, for certain movies we performed a 90° rotation and flipped the field of view to align the FOV with the imaging session used for training data. To increase signal-to-noise, reduce training and analysis processing times, and reduce file sizes, for a subset of animals we spatially downsampled their movies by the *x* and *y* lateral dimensions by conducting 2 × 2 or 4 × 4 bilinear interpolation. We found that running on downsampled movies and then upsampling the feature tracking *x* and *y* lateral coordinates was faster and often worked better than running the downsampled movie-trained model on raw movies, likely owing to the decreased SNR of the raw movies. We then exported these movies as AVI files for use with DeepLabCut.

To consistently identify vascular features across frames and imaging sessions, we selected frames from a single imaging session for mice with large rostrocaudal motion based on visual appearance. Here we used k-means clustering to select frames from distinct clusters along with manual selection to include frames when the maximal motion occurs in the movie. The latter is due to their sometimes infrequent occurrence and we wanted to include those frames in the training set. We then manually annotated key, stable vasculature within ∼20 frames and used these to train a DeepLabCut (version 2.2.3) model. For mice used in Fig. 2, **Extended Data Fig. 5**, and **Extended Data Fig. 10f**, we used the following DeepLabCut training parameters: 500,000 iterations (**Extended Data Fig. 5c**); net type, resnet50 or resnet101; dataset augmentation, imgaug; global scale, 0.8; batch size, 1; fully connected parts; and GPU, NVIDIA A5000 (24 GB ram). Model training errors with likelihood cutoff of 0.1 are 0.96, 0.87, 0.96, 1.36, and 0.81 px across each of five mice. Model test errors with likelihood cutoff of 0.1 are 2.45, 2.34, 3.97, 19.66, and 2.87 px across each of five mice. We used the resulting model to annotate the same vasculature in model-naïve frames and movies (Fig. 3c, **Extended Data Fig. 5a-b,f-g**), then conducted visual inspection of tracking-annotated movies to additionally verify model accuracy across a variety of conditions, including those outside the training data (**Extended Data Fig. 5f**).

We imported feature locations from DeepLabCut CSV files into MATLAB using a custom CIAtah function. To compensate for rotated or flipped FOVs across imaging sessions, in certain animals due to the use of multiple cameras, for a subset of animal’s videos we rotated and flipped the movie tensor as needed to match the orientation used for DeepLabCut tracking. We treated the resulting vascular tracking as control points. To reduce the influence of low-quality tracking on resulting translation matrix estimation, for each frame we only accepted features with tracking likelihood>0.99. We corrected for large shifts using point feature matching algorithms that estimates a 2-D transformation matrix (*D*, using *estgeotform2d* in MATLAB). While there are often non-rigid motion on top of the large rostrocaudal shifts, we used a rigid transformation—in contrast to similarity, affine, or projective—as that allowed us to correct for the large shifts and then handle residual motion with other methods, while preserving the mass of the image. Using the other transformation types often produced warping (shear or tilting) and scaling of the motion corrected frame, which are undesirable. We used 1,000 trials and used inliner pairs by only including those reference and motion frame point pairs that are within 20 pixels after applying the transformation. This eliminates outliers that can otherwise reduce the accuracy of the registration. Before registration and to avoid issues with NaN value pixels, we correct for photobleaching and background fluorescence in Ca^2+^ movies by detrending and spatially filtering movies as described in prior sections. We then register frames (*imwarp* in MATLAB) using *D* with linear interpolation. We checked the stability of the resulting registered movie and then fixed residual motion owing to slight variability frame-to-frame in feature localization. To do this we calculated the rigid transformation matrix (*TurboReg*) on a subsection of the FOV that was most stable and minimized overlapping features. We used *TurboReg* to estimate an affine transformation with no rotation or skew along with each frame being normalized as described previously (mean subtraction, image inversion, and subtracting movie by 2-D low-pass version of itself). We then registered the non-normalized frames to the reference frame using the 2D transformation matrix (imwarp, to avoid issues with NaNs in existing output). To improve registration in a subset of movies, we separately conducted LD-MCM motion correction of the left and right spinal cord and then combined the results before further downstream analysis.

There are some features that on particular sessions have deviated enough from the training dataset that resulted in inconsistent labeling. This can be due to a change in the vasculature size, e.g. if a vessel diameter becomes smaller during different arousal states or when the vessel disappears during the course of long-term imaging. These features can hamper motion correction and thus should be removed. As large movement of the FOV often is correlated across all feature points, it is possible to calculate the per-frame displacement correlation coefficient between all features and eliminate those that fall outside a predefined range (**Extended Data Fig. 5d**). We did this via manual inspection, but clustering or using a cut-off criterion of the mean correlation of each feature to all other features can also be used.

#### NR-MCM: Deformation correction using diffeomorphic registration with displacement fields

We load movie tensors (MATLAB) and detrend as previously described. To reduce the influence of background fluctuations, we then normalized each frame by subtracting the frame by a version of the image cross-correlated with a circular averaging filter (pillbox) of radius, one tenth the minimum row or column length (whichever was smaller). The resulting modified frame matrix was then cross-correlated by a circular average filter of radius 3 pixels. We calculated the *x* and *y* lateral displacement fields (*imregdemons* in MATLAB) using existing demons algorithm-based methods to estimate the displacement fields (Vercauteren et al., 2009). Parameters for *imregdemons* are 3 pyramidal levels; 2000, 400, and 100 iterations for each level; and accumulated field smoothing of 1.5. We then discarded the modified movie tensor. To improve cell extraction, we then divided each frame of the raw movie by a bandpass-filtered version of that frame (cutoff frequency of 0–10 cycles) that suppressed background fluctuations, such as neuropil. We performed this step before registration to avoid issues with filtering frames containing NaN values. Visual inspection and quantitative feature tracking with LD-MCM indicated that in our movies the majority of spinal cord motion is rostrocaudal (Fig 2c**,e** and **Extended Data Fig. 5e**). Thus, we set the mediolateral displacement fields to zero before image registration. This reduced the introduction of improper mediolateral shifts, especially during times of high neural activity. We applied the modified displacement field to the processed movie using linear interpolation (*imwarp* in MATLAB), resulting in a movie that closely matched the template (Fig. 2g). We visually checked the resulting movies and then reduced residual motion in the movies by applying a rigid motion correction as described in the LD-MCM section.

Beyond comparing NR-MCM to *NoRMCorre* and *TurboReg*, we further tested using another recent patch-based non-rigid registration method *PatchWarp (Hattori and Komiyama, 2022)*. We used our slightly modified implementation that reduced memory overhead and improved parallelization performance (see https://github.com/bahanonu/PatchWarp). However, we found that PatchWarp introduced blurring and the patches were visible after taking the standard deviation of the temporal dimension of each movie (**Extended Data Fig. 6a**). As the blurring would function as a low-pass filter and artificially raise the 2D correlation coefficient, we omitted PatchWarp from correlation-based analysis. We used the default PatchWarp parameters in *patchwarp_demo* (commit a4a69ed) with the following changes: rigid_template_tiffstack_num = 1, rigid_template_block_num = 1, warp_template_tiffstack_num = 1, and network_temp_copy = 0. NR-MCM can be applied after LD-MCM for cases in which, after large shifts are corrected, there is still residual non-rigid motion that needs to be corrected. However, care must be taken as the prior issue of obstructive overlying layers can lead to suboptimal NR-MCM results.

#### Extraction of neuronal shapes, locations, and activity traces from calcium imaging data

After processing each session’s Ca^2+^ imaging videos, we extracted the neuronal shapes and activity traces using existing cell extraction algorithms. We first attempted to use the widely used PCA-ICA algorithm (Mukamel et al., 2009) using μ = 0.1, termination tolerance of 5e-6, and max iterations of 1,000. We found that when using PCA-ICA many of the ICA spatial filters included spatial information from multiple cells in the movie. This is likely due to the highly correlated activity of SCPNs recorded during stimulus application, especially under anesthesia where the baseline activity is reduced, and neuron cell bodies are close to one another and more difficult to distinguish with one-photon imaging. To minimize the potential confounds that this causes, we also tested two other cell extraction methods: EXTRACT based on robust statistics (Inan et al., 2021) and CELLMax based on maximum likelihood (Kitch, 2015; Ahanonu, 2018). These methods reduce cross-talk and produce spatial filters that closely match the shape of the recorded cells. In the figures herein, we display CELLMax spatial filters and neural activity traces. We used the following CELLMax parameters (listing major ones changed from default): percent frames per iteration, 0.5–0.7; gridSpacing, 18; gridWidth, 10; movieImageCorrThreshold, 0.2; and downsampleFactorTime, 10–40. To transform CELLMax scaled probability outputs to estimated Δ*F*/*F* activity traces (or more generally movie units), we multiplied each scaled probability output by the maximum pixel intensity value of its corresponding spatial filter.

Cell extraction algorithms produce false positives—such as other sources of signal within the movie (neuropil, blood vessel artifacts, etc.) or noise—that need to be eliminated (**Extended Data Fig. 10b**). While it is possible to calculate certain parameters (signal-to-noise ratio of activity trace, rise and decay times of Ca^2+^ transients, etc.) and apply heuristics to eliminate cells, this can lead to many false negatives for cells that don’t meet prior assumptions. We are interested in capturing as many true cells as possible; thus, we manually classified every output cell throughout the paper, unless otherwise specified, using a custom GUI in CIAtah. We used criteria such as algorithm source image shape (e.g. solid circular 2-D Gaussian blob for one-photon imaging); existence of identifiable events in the Ca^2+^ imaging movie that matched transients within the algorithm’s output cell activity trace, and other features.

#### Calculating blood vessel diameter and fluorescence in anesthesia and awake imaging experiments

To calculate the blood vessel diameter during the transition from awake to anesthesia (Fig. 3p, **Extended Data Fig. 8c-e**), we first bandpass filtered the data to remove fluctuations in background intensity during imaging. We normalized each frame between zero and one using the formula 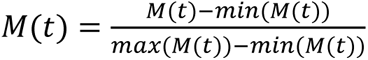. To improve consistency of vessel estimates across frames, we matched the histograms between each frame and a reference image and then multiplied all values by 150. To enhance the dorsal vein and ascending venules, we used Hessian-based Frangi vesselness filtering (Kroon, 2010) with *FrangiFilter2D* in MATLAB. This highlights the blood vessels in the movies and suppresses other non-vasculature signals (Frangi et al., 1998; Longo et al., 2020). Parameters used: sigma range of 1–20, step side between sigmas of 2, Frangi β_1_ correction constant of 0.5, Frangi β_2_ correction constant of 15, and detect blade ridges. We exported movies for processing in ImageJ.

For a subset of the movies we applied a 2 × 2 median filter. We then thresholded each frame (*setThreshold* [0.03, 1×10^30^] and “Convert to Mask”) and then selected a subsection of the FOV around the dorsal vein to process. We passed the resulting movie to the Fiji plugin *Local Thickness*, see https://imagej.net/imagej-wiki-static/Local_Thickness and (Dougherty and Kunzelmann, 2007), with a threshold of 40 or 80, depending on the movie. To avoid the algorithm introducing temporal correlations into the local thickness calculations, we calculated local thickness on each frame independently. Otherwise the method will interpret the third-order spatiotemporal tensor with dimensions *X* × *Y* × *T* as a third-order spatial tensor with dimensions *X* × *Y* × *Z*. We then manually selected a region of the dorsal vein or ascending venules and calculated the mean thickness. To display diameter relative to the start of the imaging session for the animal in Fig. 3p and **Extended Data Fig. 8d-e**, we normalized both fluorescence and blood vessel diameter to the first 4 minutes of the imaging session during which time the animals were awake.

To calculate the cross-session registration correlation, we used Frangi vesselness filtering previously described on the Thy1-GFP-M and CX3CR1-EYFP animals. This allowed us to obtain a representation of each session’s frames that are less influenced by changes in baseline or other (e.g. neural activity) fluorescence.

#### Analysis of spinal cord somatotopy

To create the mean contour map and to reduce noise, easing visualization, we median filtered the mean projection image. To calculate contour lines, which indicate the relative location of fluorescence, we used two ImageJ plugins, https://imagej.nih.gov/ij/plugins/contour-plotter.html and https://imagej.net/imagej-wiki-static/Contour_Lines, after thresholding images. We only included contour lines that mapped onto the outermost edge of the bulk GCaMP activity and used GIMP (2.10.22), so as to create transparent overlays.

#### CS-MCM: Cross-session registration

CS-MCM involved a single manual and multiple automated steps. To correct for differences in frame size across cameras, which can introduce computational difficulties, we padded the *x* and *y* lateral dimensions of each frame with zero value pixels. Thus, all frames across all imaging sessions matched the frame with the maximum *x* and *y* dimensions. To manually correct for large shifts that occurred, sometimes due to shifting of the spinal cord across sessions, which resulted in a permanent change in the field of view (FOV) through the laminectomy area, we developed a custom manual motion correction GUI within CIAtah. This approach displayed a mean frame for a reference and the current movie and allowed for translations (rigid using *imwarp* in MATLAB on each click by users) along with the ability to rotate and flip the FOV to accommodate changes in the FOV due to acquiring movies with multiple cameras for certain animals. We performed this step primarily to reduce the search space and complexity of the motion correction for the automated portions. However, to automate the manual step, we also demonstrated that we can use LD-MCM for feature identification and initial motion correction across sessions (Fig. 3d, **Extended Data Fig. 5g**, **Supplementary Video 10**). We then performed rigid motion correct within each session using *TurboReg* on a subsection of the FOV, leading to a stable movie for each imaging session. Next, to speed up motion correction, we conducted cross-session motion correction between frames consisting of the mean calculated across all temporal pixels for the current session and a constant reference session (Fig. 3d). To further improve registration, we performed affine-based registration without rotation or skew followed by an additional affine-based registration with rotation and no skew. We used the following TurboReg parameters: 30 pixel x and y smoothing, 6 pyramid levels, normalization with mean subtraction followed by pillbox disks of radii 20 (for subtraction) and 10 (on post-subtraction frame), and used the *transfturboreg* function (instead of *imwarp*) for image transformation. Lastly, to further refine the alignment, we concatenated the *X* × *Y* × *T* movie tensors from all *N* imaging sessions into a *X* × *Y* × (*T*N)* movie tensor, then conducted *TurboReg* motion correction on the combined movie using the same reference frame for all frames (Fig. 3e). These movies were then used to conduct downstream analysis, such as calculating changes in CX3CR1-EYFP fluorescence over time (Fig. 6h**-j**).

#### Cross-session cell identification

To align cells across imaging sessions in the most computationally efficient manner, we performed motion correction and cross-session alignment, after performing cell extraction and manual curation. We have previously shown that this is a fast and reliable method of cross-session alignment and reduces the computational complexity and memory requirements that occur with directly registering the movie tensors (Corder et al., 2019; Ahanonu and Corder, 2022). We used *computeMatchObjBtwnTrials* within CIAtah to align cells across sessions with the following parameters: *TurboReg* motion correction with affine transformations (rotation enabled and skew disabled) or or displacement field (see NR-MCM section), one or five rounds of motion correction consisting of sequential centroid- and cell shape-based registration, centroids greater than 5 or 15 pixels distance are not matched, and cell shapes must have an image correlation >0.6 to be matched. The spinal cord can grow and deform during long-term imaging, leading to a difference in the distance between the strip of cells on the left and right side of the spinal cord. This can reduce the accuracy of cross-session registration when registering cells from both sides simultaneously. To get around this issue we registered left and right sides of the spinal cord separately by removing cells on the contralateral spinal cord using a custom GUI added to *computeMatchObjBtwnTrials*. To allow visualization of cross-session matched cells, we then colored coded matched cells based on their global cell identification number across imaging sessions; cells without cross-session matched data are colored gray.

#### Cross-session fluorescence intensity

To demonstrate the consistency of imaging over time, we used fluorescence intensity as this will change as optical clarity is reduced, e.g. fibrosis is anticipated to and often caused a drop in fluorescence intensity. For each animal, we took 6 frames from the 2 × 2 binned widefield imaging movie and then calculated the mean intensity over all pixels in all frames. For certain sessions, where no movie is available, we used the signal widefield reference image collected during each day of imaging. To compensate for changes in fluorescence intensity values across multiple CCD and sCMOS cameras used in the study during one-photon imaging, we attempted two methods. For the first, we imaged GFP+ cells in the same area of the basolateral amygdala, from a slide containing 100-µm thick sections from a Thy1-GFP-M mouse. This was repeated with each camera using the same objectives used for each animal (e.g. Fluar 5x/0.25NA). To only include GFP+ signal in the calculation, we identified cells and extracted their mean signal intensity using a custom MATLAB function in which we identified cells by applying the following operations: spatial bandpass filtering (keep spatial frequencies 5-40 cycles), detected edges using Sobel method, dilated the edges and filled the holes to identify cell shapes, removed any cells at the border of the image, eroded the binary image to further separate cells and eliminate noise, and then segmented cells (using *regionprops* in MATLAB). We calculated the mean signal based on all pixel values inside each segmented cell’s binary mask. To produce the cross-session fluorescence intensity curves, we then constructed a correction factor for each camera relative to the mean signal from the CoolSNAP EZ camera and divided the total whole-frame fluorescence intensity values by this correction factor.

This procedure corrected for the large disparity in intensity values, but still produced residual differences, likely owing to differences between GCaMP and GFP expression across animals and to slight changes in the imaging setup when cameras were swapped or during routine maintenance. We thus used a second method in which we normalized each curve by calculating the mean fluorescence intensity for each animal for each camera (and the camera mode used for that animal, e.g. sensitivity vs. dynamic range) and divided all intensity values associated with each camera to its own mean intensity to produce the final displayed curves (Fig. 3f, Fig. 5c, and Fig. 6e).

#### Microglial analysis after nerve injury

To calculate relative changes in microglia activity before and after nerve injury, we performed CS-MCM on each animal, as described in prior sections, using 5 frames taken from each imaging session’s 2 × 2 binning acquired movies. We then manually selected rectangular regions on the left and right side of the spinal cord, excluding areas that were outside the FOV or were substantially blocked by cement or other overlying features. We cropped the movie to these left and right side regions, then calculated the mean of all pixels on each *X* × *Y* frame, which created a *1* × *T*N* vector, where *N* is the number of imaging sessions. Next, we calculated the mean on each *1* × *T* vector for each movie. For injured animals, we calculated the ratio of the ipsilateral and contralateral side, using the formula 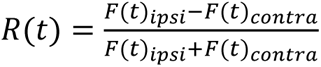 for each frame (Fig. 6g**-h**). For naive animals, we arbitrarily chose a side as “ipsi’’ and used the same calculation. To compare across animals that had different baseline ratios between the fluorescence on the left and right spinal cord, we normalized curves in Fig. 6g**-h** by subtracting the mean value of all baseline sessions.

### Statistics and reproducibility

We conducted statistical analysis in RStudio (1.4.1106) using R (4.1.0). We created Fig. 1n**-p**, Fig. 6f, and **Extended Data Fig. 4l** using the R library *ggplot2*. We indicate significance in figures using the following nomenclature: *** is p < 0.001, ** is p < 0.01, and * is p < 0.05. To assess impact on general locomotion (Fig. 1n), we performed a one-way ANOVA (*aov* in R) on the mean locomotor speed in the open field, with groups consisting of pre- and post-surgery animals. We then performed post-hoc analysis with a Dunnett’s test comparing each post-surgery condition to the pre-surgery baseline. To assess impact on coordinated locomotor behavior (Fig. 1o and **Extended Data Fig. 4l**), we performed a two-way ANOVA, which evaluated the effect of group and trial on latency to fall. We found an effect only on “trials” but not on group or interaction between surgery and group. We then performed a one-way ANOVA within each trial so as to only compare the groups, followed by a post-hoc Dunnett’s test comparing each group to pre-surgery baseline. We excluded animals that exhibited signs of leg paralysis and were thus unable to perform the task. To determine whether spinal chamber implantation impacted mechanical sensitivity (Fig. 1p), we performed a two-way ANOVA comparing interaction between hindpaw and surgical state. We then performed a one-way ANOVA followed by a post-hoc Dunnett’s test comparing each group to pre-surgery baseline. We excluded data points for CX3CR1-GFP animals after they had nerve injury, as that would confound interpretation due to hypersensitivity. To compare LD-MCM rostrocaudal displacement to other methods (Fig. 2f), instead of having to perform a statistical test of difference in variance between samples (e.g. Levene’s test), we transformed the data by computing the absolute difference in distance for each feature and time point from its mean location for that animal’s imaging session. We then computed the mean over all frames for a given session and performed statistics on these values. Next, we performed a one-way repeated measures ANOVA followed by a post-hoc Dunnett’s test comparing each method to LD-MCM. To compare NR-MCM mean frame correlation coefficient, used as a measure of the reduction in spinal cord non-rigid motion (Fig. 2k), we took the mean of all frames for each imaging session and then conducted a one-way ANOVA by method followed by a post-hoc Dunnett’s test comparing each method to NR-MCM. To directly assess von Frey mechanical sensitivity (Fig. 6f), we conducted a paired T-test comparing before and after surgery across each hindpaw and surgery group. Multiple comparisons correction with Holm– Bonferroni method.

We ran or compiled code in MATLAB (2022b and occasionally 2021b), Python (3.9.12) using Anaconda (4.12.0) environment, RStudio (1.4.1106), R (4.1.0 and 4.0.2), ImageJ (1.53d), Fiji (1.53q), Saleae Logic (various iterations of version 2, e.g. 2.4.6), and the Arduino IDE (1.8.13 and 2.0). We converted between file containers (e.g. AVI to MP4) using ffmpeg (version 2020-10-21-git-289e964873). We ran image acquisition and conducted data analysis on computers running Windows 10.

### Data availability

Data generated in this study are available from the corresponding authors by request.

### Design availability

3D STL and STEP files of the side bars and stabilizing plate along with a 3D model and TIFF stack of the entire mouse body from one of our microCT scans (Fig. 1a) can be found online: https://github.com/basbaumlab/spinal_cord_imaging. Any future updates to the design or additional files will be published on that repository.

### Code availability

Code for processing Ca^2+^ imaging data is available as part of the CIAtah software package under an MIT license (see LICENSE file): https://github.com/bahanonu/ciatah. Code for LD-MCM (feature identification followed by control point motion correction) and NR-MCM (deformation correction using displacement fields) will be integrated into CIAtah and any future updates will be published on that repository.

## SUPPLEMENTARY TEXT

### Surgical considerations for chamber implant and assembly

The side bar position will vary slightly for each mouse, so the freedom of movement of the side bars greatly facilitates the positioning of the chamber during surgery. To adjust the width of the metal chamber, instead of using small metal screws that are difficult to handle, we use Super Glue to attach the metal parts. Metal surfaces covered with Super Glue are then clamped together setting a strong bond, which obviates the need for set screws. Surgifoam® is used in the first procedure for hemostasis and protection of the intervertebral tissue, before Kwik-Sil. To keep the tissue hydrated, saline-soaked Surgifoam pieces are placed in intervertebral areas and in rostral and caudal areas of open tissue, until the bars are in place. After the bars are in position, the Surgifoam is removed and the space is briefly dried with a sponge. Small drops of Kwik-sil are then applied to cover the intervertebral space; these sit at room temperature until cured. The Kwik-sil mask is maintained throughout cementing steps. Critically, the mask protects intervertebral space from the dentin activator and liquid cement, which are applied in bulk to free-flow around the mask. An unprotected intervertebral space is vulnerable to chemical and mechanical injury, which will not only create a lot of bleeding but also lead to permanent damage to the spinal meninges and spinal cord.

The needles that are inserted perpendicular to the spinal column, through the dorsal spinous processes (DSP), are a key step and allow the components to be constructed around the vertebrae. We use a 33G needle (Accuderm, NP335) with a beveled edge and a bore size diameter of 0.2 mm. Place the needle on a 1mL syringe and rotate the cutting edge to facilitate driving it through the DSP bone. When the needle is through, snip the needle at the base of the shaft to have equal lengths of the needle on each side.

**Extended Data Fig. 2a-i** illustrates the cross-sectional anatomy of a vertebra and the location of the dorsal spinous process, facet joints, and lamina, and how the side bars are positioned relative to these vertebral landmarks. Note that only the facet joints surrounding the middle vertebrae (T13) are exposed during surgery. Before cement application, it is important to debride any tissue above the vertebral bone as this tissue could come into contact with the cement. To further improve the bone cement (PMMA) connection to the bone, a pretreatment with a dentin activator (included in Metabond kit) is recommended. Of course, any cement placed on soft tissue or ill-prepared bone will not hold, as can happen when blood debris sticks to the bone. Before installing the stabilizing plate, gently press down the middle vertebrae and carefully lower both side bars, so that the alignment between vertebrae is as flat as possible before fixation.

### Post-operative recovery

#### Robust recovery

The animals are fully mobile after surgery. At 24 hrs, the animals are somewhat limited in their speed of movement but are alert, ambulate, and groom. By 48 hrs, the animals spontaneously rear and show normal movements.

#### Implant location considerations

Placing side bars under the facet joints surrounding the middle vertebra of interest (along with the dorsal spinous process needles on the flanking vertebrae) can theoretically be attempted at any triplet of vertebrae along the spine. For instance, we were able to successfully image the L6/S1 spinal cord by a T13-L2 side bar placement and L1 laminectomy (Fig. 4i**-k**). However, L1 animals commonly presented with severe urinary retention and succumbed 48-72 hours after surgery. We have not explored whether other rostrocaudal positions are possible. Post-op excretion of urine through manual compression at 24 hours, under anesthesia, was necessary for survival in cases associated with L1 surgery. In some cases, there were post-op complications with a slow-release form of Buprenorphine, namely, Ethiqa.

#### Cutaneous self-injury

Occasionally chemical or mechanical intervertebral injury during surgery can produce a focal, but treatable, problem. The animals will show focal skin lesions due to self-inflicted biting. For the T13-centered implant, which overlies the L4/L5 spinal cord segments, biting occurred in the L4/L5 dermatome, including the hindpaw. Attention to the intraoperative, intervertebral mask using Kwik-sil, will prevent the potential toxicity of liquid dentin activator (ferric chloride) or uncured cement (PMMA).

### Mechanism of fibrosis inhibition with fluoropolymers

In previous spinal cord optical imaging methods (Farrar et al., 2012), post-laminectomy fibrosis was abated by room-temperature vulcanizing (RTV) silicone adhesive, namely, Kwik-sil, applied directly over the meninges. However, we found that the dura becomes re-established under a bare Kwik-Sil or a precured Polydimethylsiloxane (PDMS) layer. Also, we commonly observed that, within a month, fibrosis that blocks the field of view emerges under the silicone layer. Therefore, we adopted two fluoropolymer materials, GORE® PRECLUDE® Pericardial Membrane, and Teflon™ AF. To skip the intermediate PRECLUDE® membrane step entirely, we attempted untreated Teflon AF placement immediately after laminectomy, but this approach failed. PRECLUDE®’s greater effectiveness at inhibiting fibrosis after laminectomy may be due to its microporous texture, which may accelerate the mechanical adhesion of meningeal cells. Teflon AF has a remarkably low surface energy and thus a molecular coating of hydrophilic units may be needed for adhesion, similar to another fluoropolymer, CYTOP, and its successful interface with the brain meninges after PEO-coating (Takahashi et al., 2020). Indeed, increasing the hydrophilicity of Teflon AF, by molecular etching, specifically an oxygen plasma coating (Harrick Expanded plasma cleaner PDC-001, 3 minutes at 55W), demonstrated improved compatibility over untreated Teflon AF in the post-laminectomy period. However, to establish a strategy that does not require sophisticated material pretreatment and is likely more accessible to the field, we applied the microporous PRECLUDE® membrane immediately after laminectomy and then swapped for the Teflon AF membrane in a second operation.

The mechanism of the fibrosis-inhibiting properties of ePTFE, in the form of PRECLUDE®, or Teflon AF is unresolved. In some cases, we observed dramatic evidence of transient meningeal neo-vascularization after placing Teflon AF (Extended Data Fig. 8). What is clear is that for months to years, surgically implanted PRECLUDE® Pericardial membranes develop few to no adhesions, allowing the material’s quick removal and an overall decrease in time of cardiovascular surgery at reoperation (Loebe, 1993; Zehr, 1993). There are also reports of ePTFE’s effectiveness in preventing postoperative peridural fibrosis (Cemil, 2009), which has major implications in combating Failed-back syndrome. Again, the outstanding hydrophobic, low surface energy and no nonspecific absorption properties of Teflon contribute to its chronic biocompatibility without integrating physically with the tissue. The overarching theory is that Teflon is an inert physical material that can prevent mechanical contact between meningeal layers and connective tissue, for an extended time, and thus would allow independent healing of the meningeal surface (Ahmad, 2020). In this mechanism, ePTFE is an effective barrier agent above the spinal cord. Thus, dural regeneration and fibrosis are marginalized to locations that are not problematic for imaging. This mechanism has also been suggested to enable cranial imaging studies (Shtoyerman et al., 2000; Chen et al. 2002). Other materials, namely polyethylene terephthalate (PET) film, used in brain imaging protocols may play a similar role as Teflon AF, though remain untested as agents that could preserve spinal visibility (Ghanbari et al., 2019). In future studies, it is possible that reducing the thickness of the Teflon AF to a nanosheet would adhere more tightly to the spinal meninges, than did the thicker membranes used in this study. Though nanosheet handling is difficult, fluoropolymer nanosheets of 130 nm thickness, namely CYTOP, have proven sufficient for long-term cranial imaging (Takahashi et al., 2020). Interestingly, in future approaches, as nanosheets are penetrable, it may be possible to permit electrode manipulation, optical recording, and chronic micro-injections in a stable state of fibrosis inhibition.

### Equipment and materials

#### Customized surgical table

To construct the spinal cord surgical station, two Thor labs 3-axis manipulators equipped with clamps were installed on an MB618 plate. In the middle of the clamps, two MB412 plates created slanted armrests for a customizable, 3D-printed surgical bed, following previous designs (Farrar et al., 2012). Additional components of gas delivery (Posi-vac, Patterson Scientific), active waste gas scavenging (EVAC 2, Patterson Scientific), vacuum aspirator (in-house vacuum flask), and isothermic bed (FHC) were integrated into the custom surgical set-up. For details see: (**Extended Data Fig. 1c**) and the GitHub containing CAD files of the individual 3D printed components.

### Step-by-step surgical procedure

The following step-by-step surgical procedures are complemented by diagrams in Supplemental Fig. 1-3 and **Supplementary Videos 2-4**.

Chamber implant (see Supplemental Fig. 1 and Video 2)

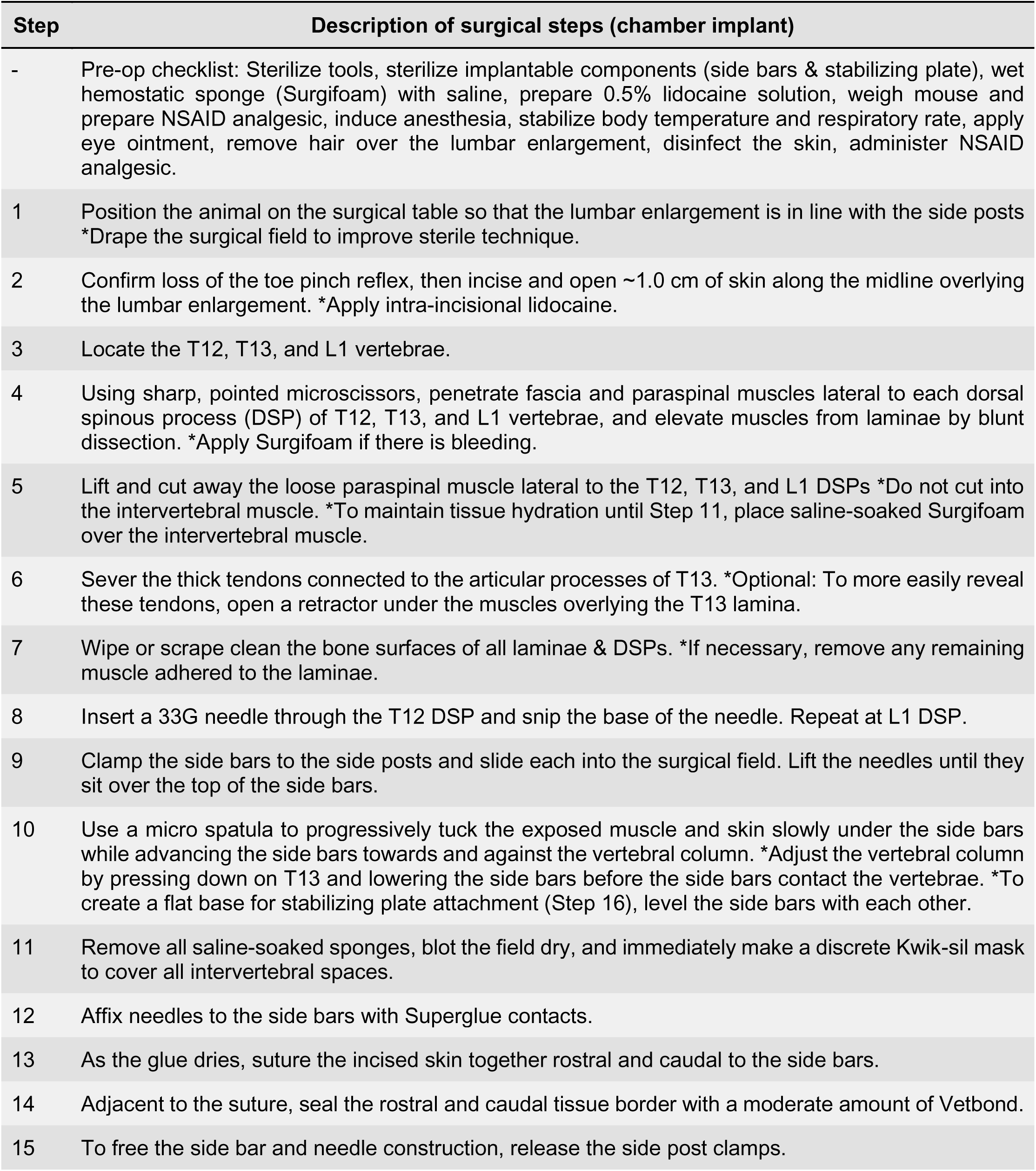
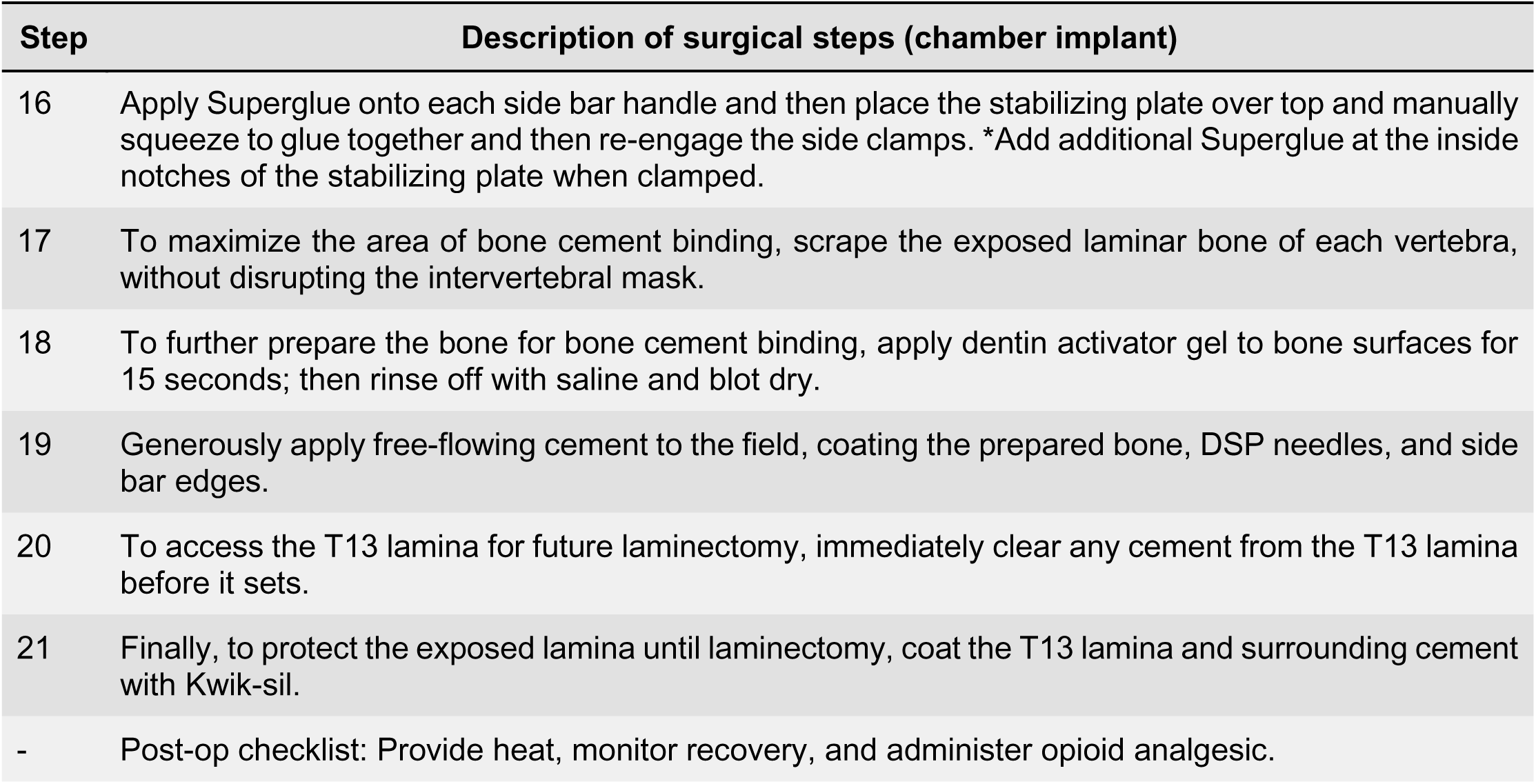

Laminectomy (see Supplemental Fig. 2 and Video 3)

| Step | Description of surgical steps (laminectomy) |
| --- | --- |
| - | Pre-op checklist: Weigh the mouse and prepare NSAID analgesic, induce anesthesia, stabilize body temperature and respiratory rate, and apply eye ointment. |
| 1 | Remove the Kwik-sil coat over the T13 lamina. *If necessary, reduce cement at the edges of the T13 lamina with a microdrill burr. |
| 2 | Cut through the lamina as laterally as possible, on both sides. |
| 3 | Transect the intervertebral ligaments to free the lamina and then remove the lamina *Take care to separate any attachments to the lamina before pulling away. |
| 4 | Immediately add saline-soaked hemostatic sponge over the spinal cord to maintain hydration and stop any bleeding. |
| 5 | Separate the dural layer and retract laterally to expose the leptomeninges (arachnoid and pia mater) and replace the hemostatic sponge. *Widen the laminectomy, if necessary, to expose the characteristic grayish hue of the dorsal horn. |
| 6 | Shape a single piece of the PRECLUDE membrane to the size of the exposed spinal cord. |
| 7 | Remove Surgifoam and place the PRECLUDE membrane directly over the spinal cord in contact with the leptomeninges. |
| 8 | Place a small amount of saline-soaked Surgifoam around the border of the Preclude membrane (this helps limit ingrowth). |
| 9 | Blot dry the Surgifoam and apply Kwik-sil to close off the entire field and hold the Preclude membrane in place until window placement. |
| - | Post-op checklist: Provide heat, monitor recovery, and administer NSAID analgesics. |

Window placement (see Supplemental Fig. 3 and Video 4)

| Step | Description of surgical steps (window placement) |
| --- | --- |
| - | Pre-op checklist: Weigh the mouse and prepare NSAID analgesic, induce anesthesia, stabilize body temperature and respiratory rate, and apply eye ointment. |
| 1 | Remove the Kwik-sil coat and PRECLUDE membrane to expose the spinal cord. |
| 2 | Immediately add saline-soaked hemostatic sponge over the spinal cord to maintain hydration and stop any bleeding. |
| 3 | Shape a single piece of the Teflon AF membrane to the size of the exposed spinal cord and place it directly over the spinal leptomeninges. |
| 4 | Add a small amount of Kwik-sil to flow over the Teflon AF & laminectomy area. |
| 5 | Immediately set a 3.0 mm circular coverslip *Press down on the coverslip to squeeze out the excess Kwik-Sil adhesive. |
| 6 | Allow Kwik-sil adhesive to harden for 10 min. |
| 7 | Permanently set the coverslip with bone cement or Norland optical adhesive *Completely cover the edges of the Kwik-sil layer to seal the preparation. |
| - | Post-op checklist: Provide heat, monitor recovery, and administer NSAID analgesics. |

## SUPPLEMENTARY FIGURES

**Supplementary Fig. 1.**
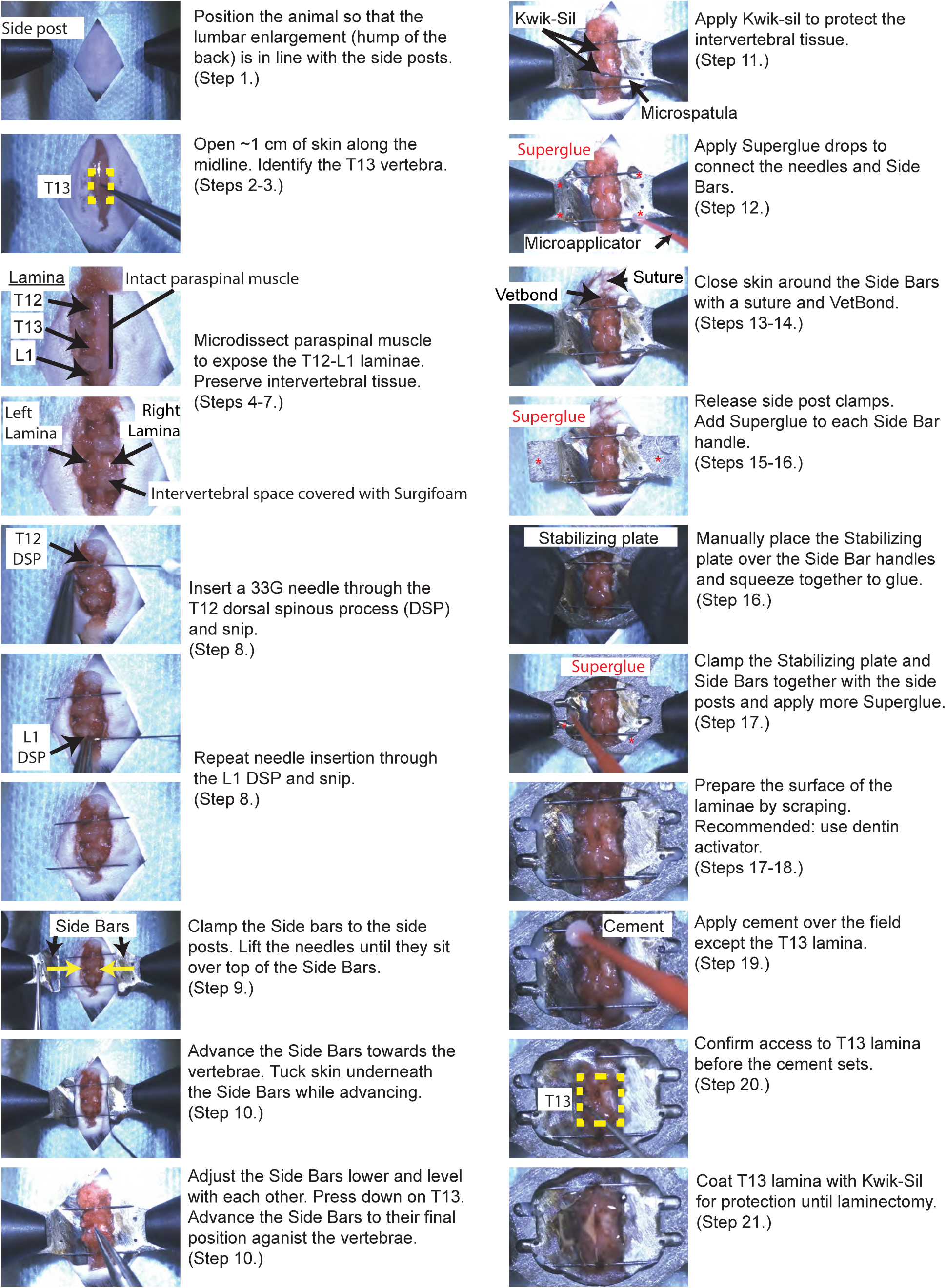
Key frames of the first procedure - Chamber implantation. Three metal pieces are implanted paraspinally, in 21 steps.

**Supplementary Fig. 2.**
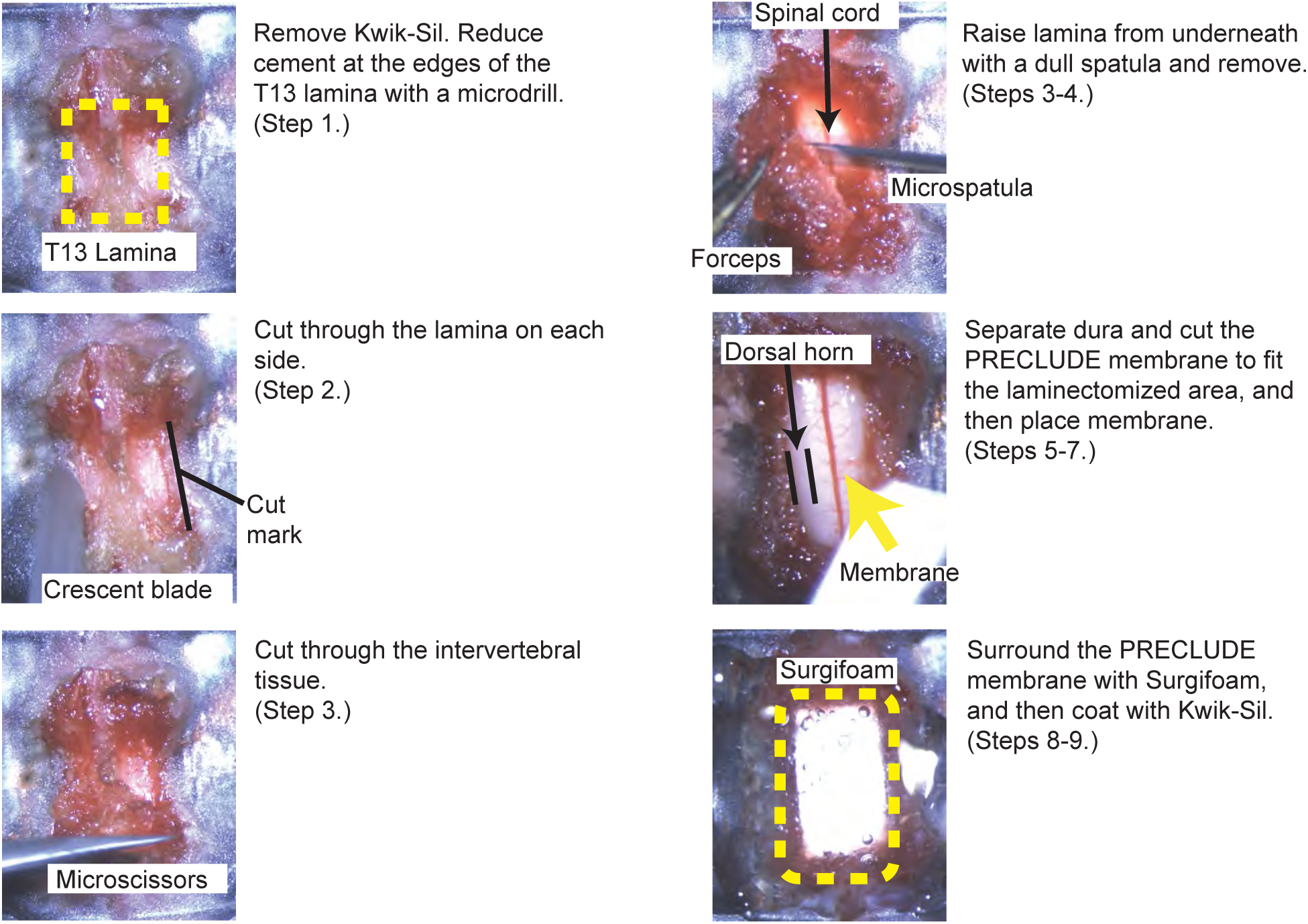
Key frames of the second procedure - Laminectomy. 1 week or more after the bar-implantation, a laminectomy is performed in 9 steps.

**Supplementary Fig. 3.**
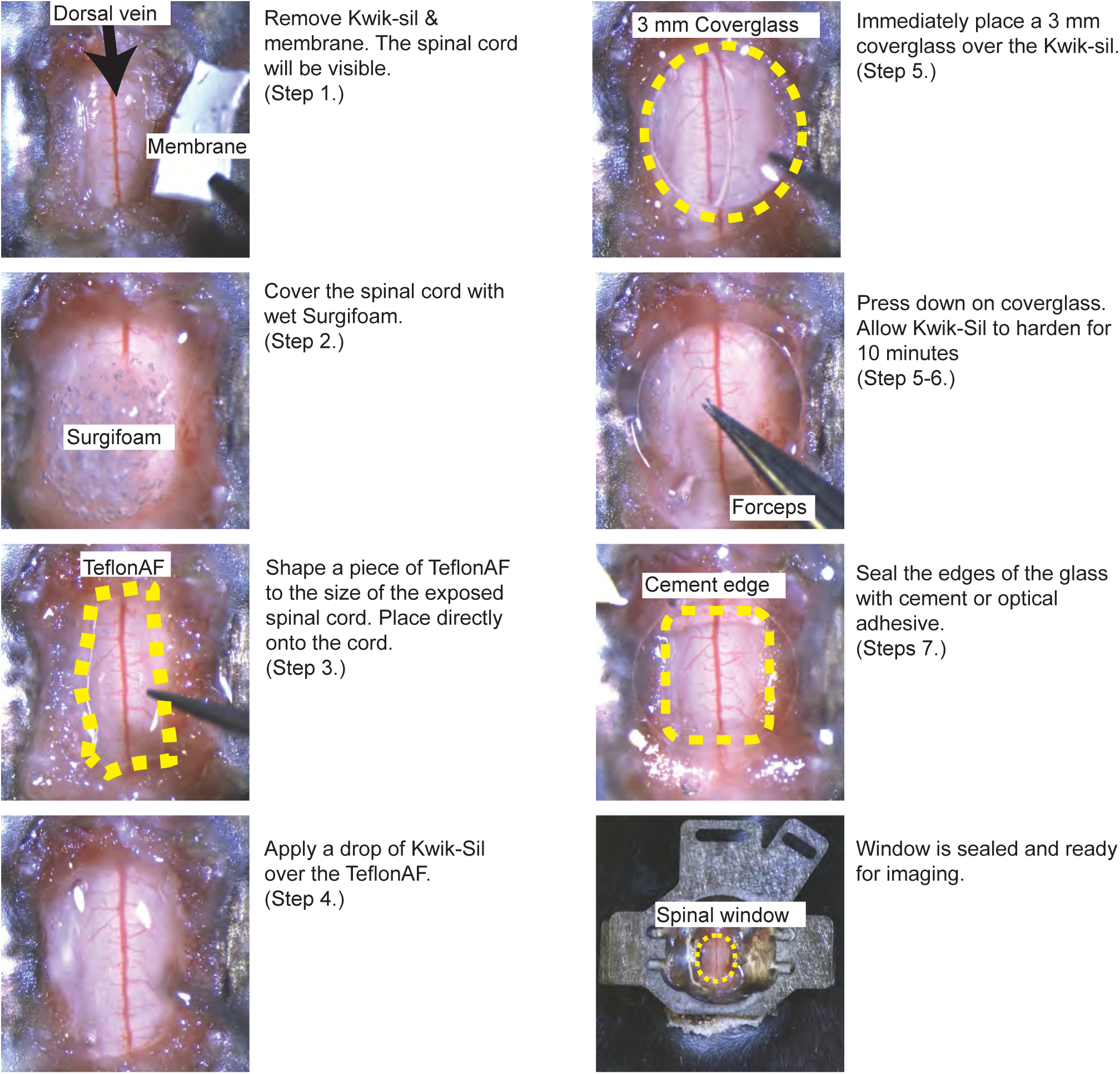
Key frames of the third procedure - Spinal cord window placement. A glass window is installed by following these 7 steps.

## SUPPLEMENTARY TABLES

**Supplementary Table 1.**
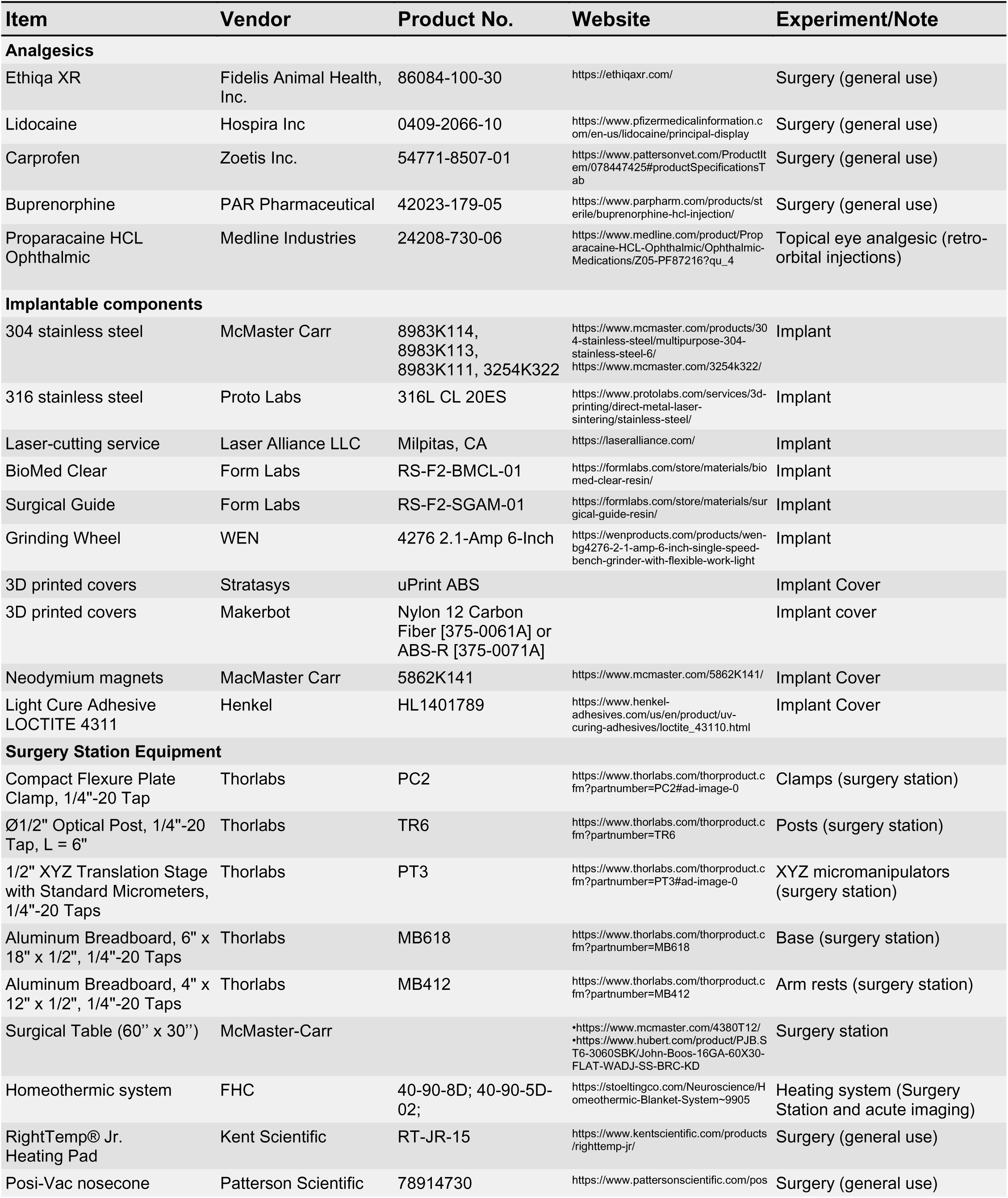
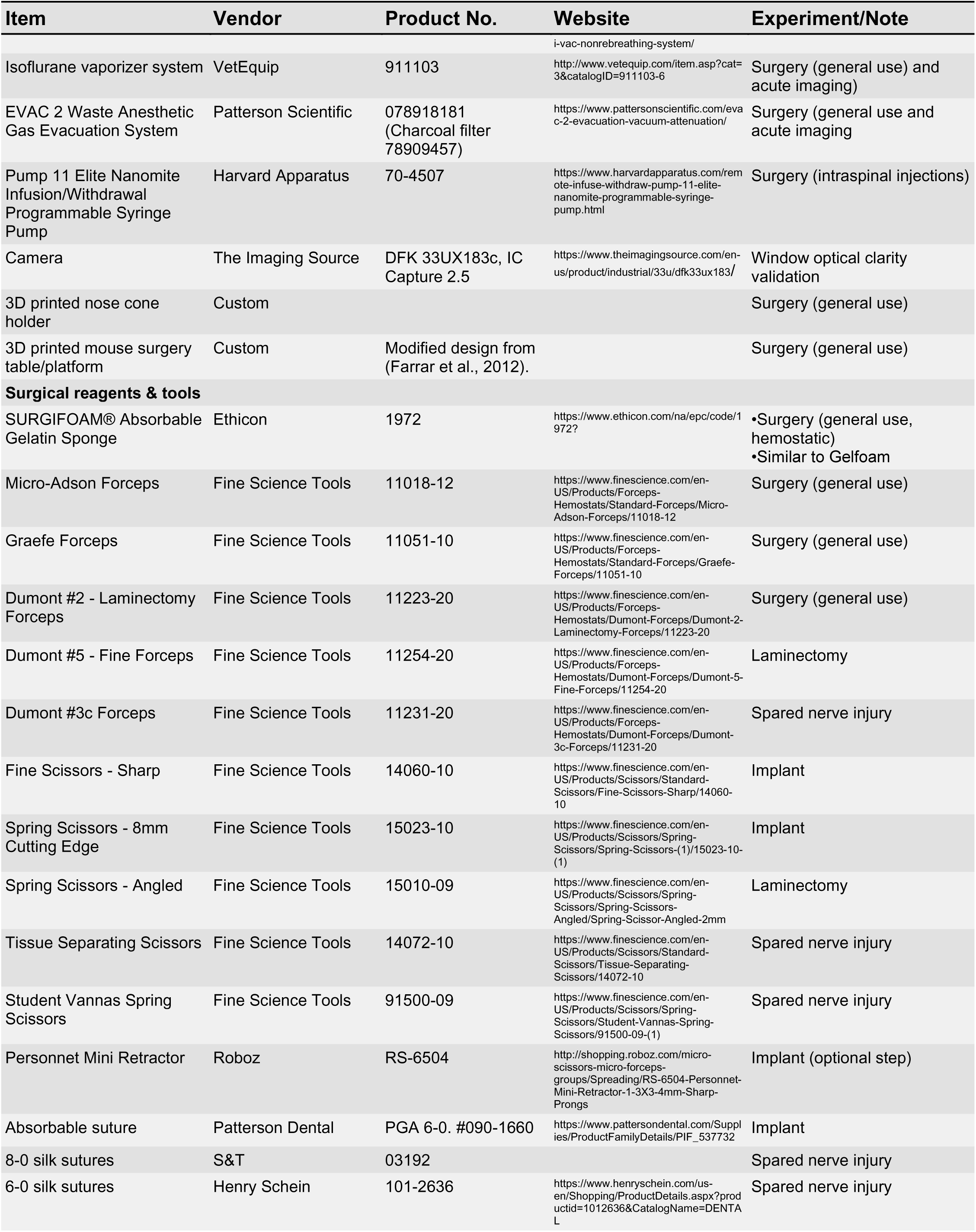
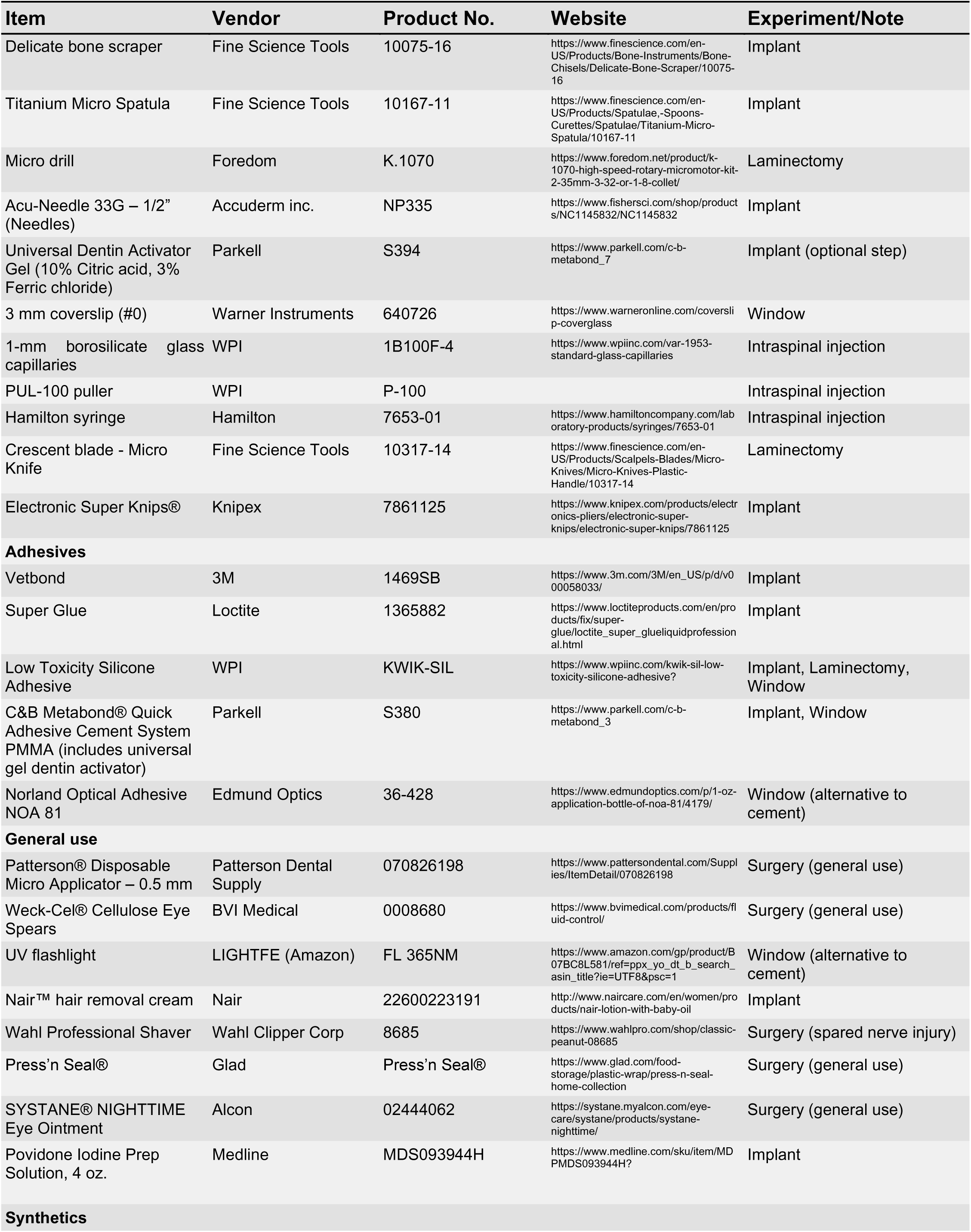
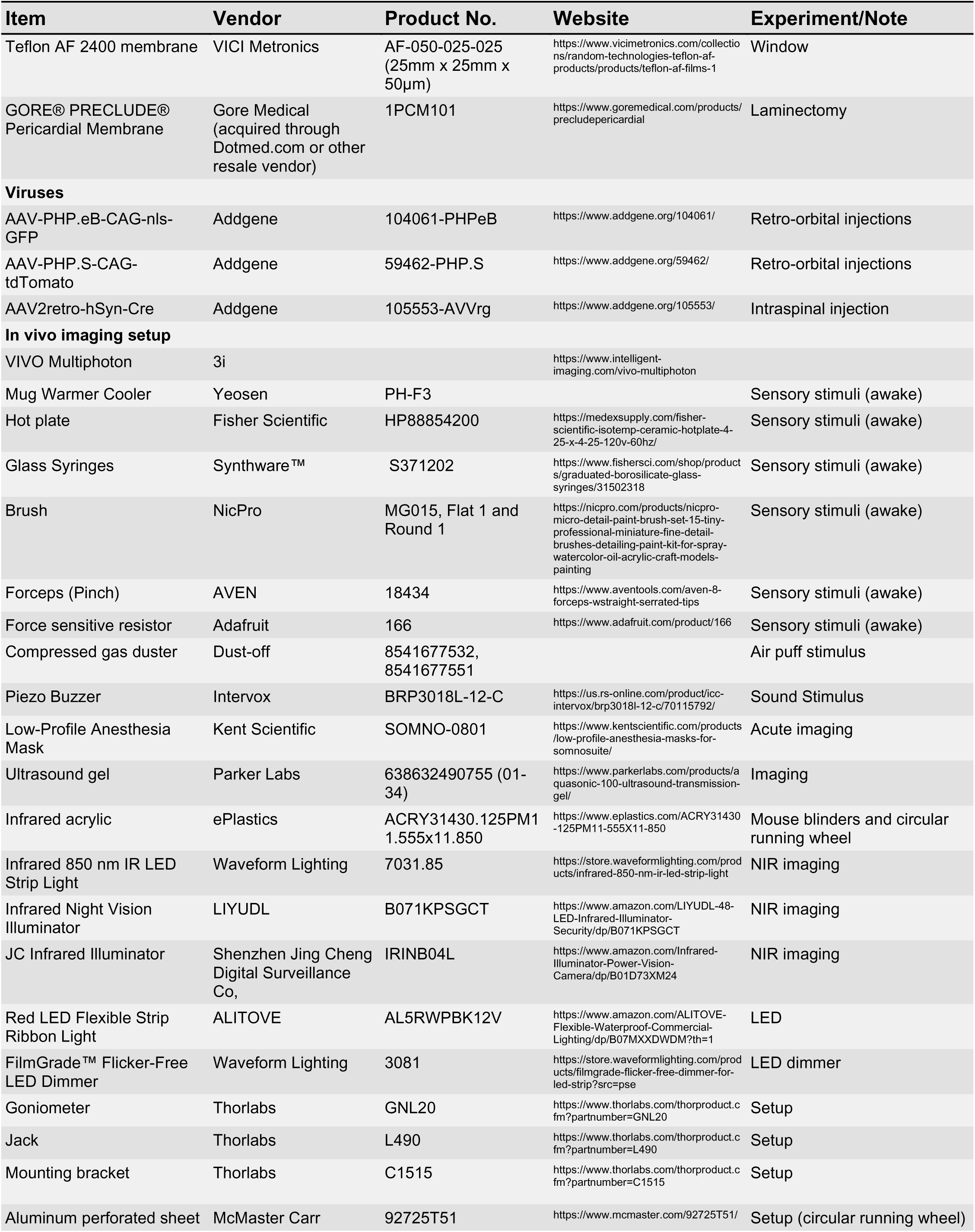
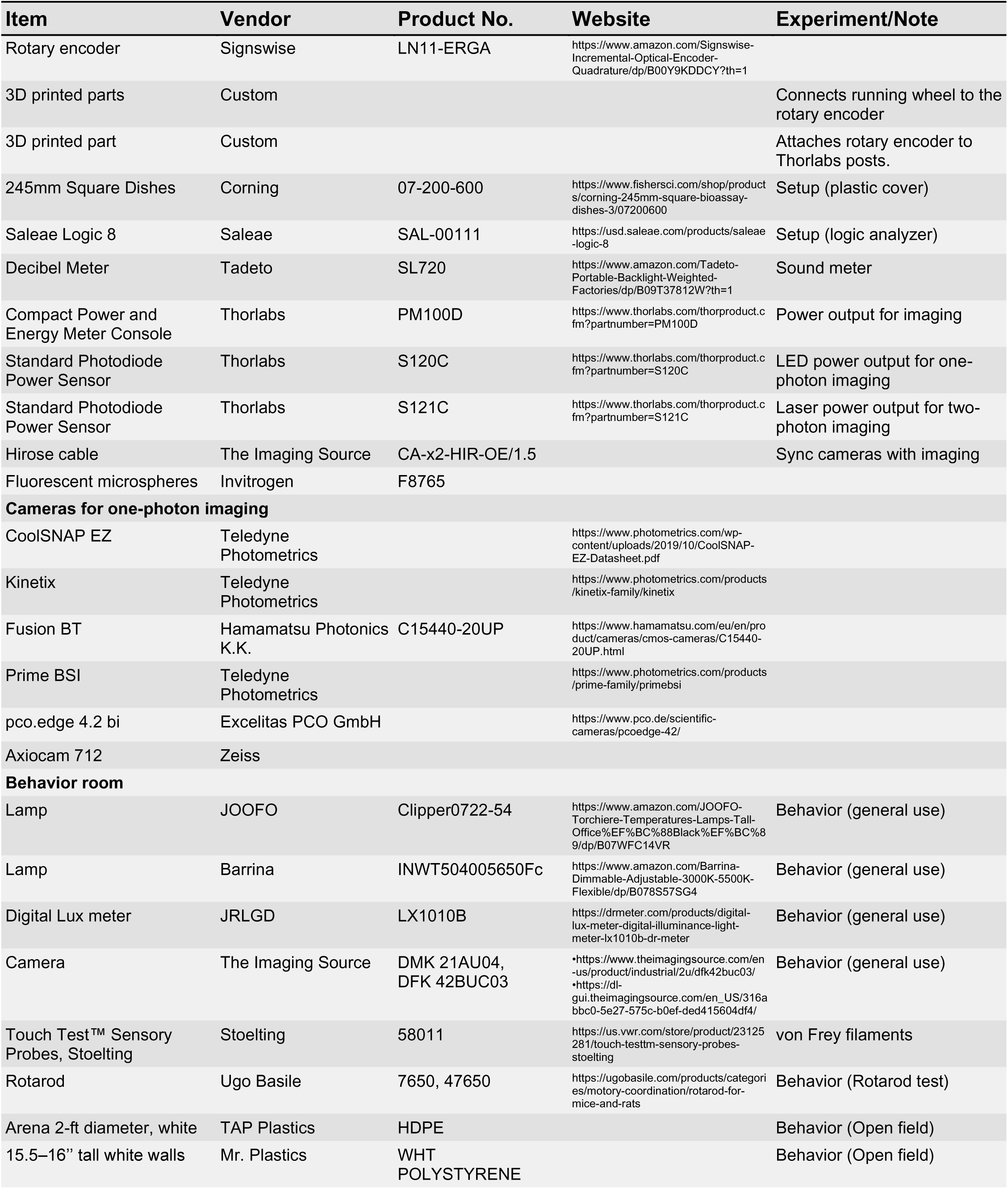
Materials used for in vivo imaging of the mouse spinal cord

